# Comparison of bibliographic data sources: Implications for the robustness of university rankings

**DOI:** 10.1101/750075

**Authors:** Chun-Kai (Karl) Huang, Cameron Neylon, Chloe Brookes-Kenworthy, Richard Hosking, Lucy Montgomery, Katie Wilson, Alkim Ozaygen

## Abstract

Universities are increasingly evaluated, both internally and externally on the basis of their outputs. Often these are converted to simple, and frequently contested, rankings based on quantitative analysis of those outputs. These rankings can have substantial implications for student and staff recruitment, research income and perceived prestige of a university. Both internal and external analyses usually rely on a single data source to define the set of outputs assigned to a specific university. Although some differences between such databases are documented, few studies have explored them at the institutional scale and examined the implications of these differences for the metrics and rankings that are derived from them. We address this gap by performing detailed bibliographic comparisons between three key databases: Web of Science (WoS), Scopus and, the recently relaunched Microsoft Academic (MSA). We analyse the differences between outputs with DOIs identified from each source for a sample of 155 universities and supplement this with a detailed manual analysis of the differences for fifteen universities. We find significant differences between the sources at the university level. Sources differ in the publication year of specific objects, the completeness of metadata, as well as in their coverage of disciplines, outlets, and publication type. We construct two simple rankings based on citation counts and open access status of the outputs for these universities and show dramatic changes in position based on the choice of bibliographic data sources. Those universities that experience the largest changes are frequently those from non-English speaking countries and those that are outside the top positions in international university rankings. Overall MSA has greater coverage than Scopus or WoS, but has less complete affiliation metadata. We suggest that robust evaluation measures need to consider the effect of choice of data sources and recommend an approach where data from multiple sources is integrated to provide a more robust dataset.

## 1. Introduction

### 1.1 Why should we care?

Bibliometric statistics are commonly used by university leadership, governments, funders and related industries to quantify academic performance. This in turn may define academic promotion, tenure, funding and other functional facets of academia. This obsession with *excellence* is highly correlated to various negative impacts on both academic behaviour and research bias (Anderson et al., 2007; Fanelli, 2010; van Wessel, 2016; Moore et al., 2017). Furthermore, these metrics (such as citation counts and impact factors) are often derived from one of the large bibliographic sources such as Web of Science (WoS), Scopus or Google Scholar (GS). Given the potential differences between their coverages of the scholarly literature, quantitative evaluations of research based on a single database present a risky basis on which to make policy decisions.

In a related manner, these bibliographic sources and metrics are also used in various university rankings. For example, Scopus is utilised by *QS University Rankings* and *THE World University Rankings* for citation counts, while *Academic Ranking of World Universities* makes use of WoS for a similar purpose^1^. These rankings, and others, have been driving systematic transformations to higher education, including increased focus on student satisfaction, and changes in consumer behaviour. A focus on performance according to the narrow set of measures reflected in university rankings comes with a number of side effects, such as institutional homogenization, distorting disciplinary balance and altering institutional focus (Shin & Toutkoushian, 2011; Hazelkorn, 2007). As a result of heavy criticism by the scientific community, university rankings (together with impact factors) have recently been boycotted by some academic stakeholders (Stergiou & Lessenich, 2014). This also includes domestic rankings^2^. Nevertheless, they are still widely marketed and used, without necessarily being carefully comprehended by decision makers (e.g., policymakers, students).

Bibliographic data sources evidently make a significant impact on the academic landscape. This makes the selection and use of such databases essential to various stakeholders. As such, a number of important research questions arise:

1. Are there differences across bibliographic databases?
2. If there are differences, can we characterise them?
3. Do these differences matter? How do they matter?
4. And, to who these differences matter?

Answers to these questions may shed light on better and more robust ways to understanding scholarly outputs. For all of these questions our concern is how these different analytical instruments differ in the completeness, comparability and precision of information they provide at the institutional level. Our focus is not on reconstructing a ‘true’ view of scholarly outputs but in a comparison of this set of tools.

### 1.2 Literature review

Citation indexing of academic publications began in the 1960s, with the introduction of the *Science Citation Index* (SCI) by Eugene Garfield. This was followed by the annual release, starting from 1975, of *Impact Factors* through *Journal Citation Reports*. This was initially developed to select additional journals for inclusion in the SCI. At that stage, much of the citation extraction was done manually (e.g., using punched cards as input to primitive computers) and results were restricted to a niche selection of articles and journals. However, with the explosion of the Internet in the 1990s, citation indexing became automated and led to the creation of *CiteSeer* (Giles et al., 1998), the first automatic public citation indexing system.

The rapid up-scaling of citation records created opportunities for new research explorations and bibliographic services. The former is often driven by citation analysis in the fields of bibliometrics and scientometrics, where quantitative evaluations of the academic literature play major roles. The latter is evidenced by the rise of large bibliographic and citation databases. Some of the most popular databases include WoS, Scopus, GS, and, more recently, Microsoft Academic (MSA).

WoS was the only systematic source for citation counts until 2004, when Scopus and GS were introduced. One of the earliest comparisons of these three sources was done by Jacsó (2005). The article reported on search results for citations to an article, citations to a journal and citations to top 30 most cited papers in a particular journal. At that time, WoS had the highest number of records simply because of its longer time span, Scopus had the widest coverage for more recent years, and GS had the lowest number of records with very limited search functions and incoherent metadata records.

Other early studies showed that Scopus offered 20% more coverage (than WoS) in citations, while GS (although with good coverage) had inconsistent accuracy in its results (Falagas et al., 2008). A number of studies have shown that the average citation counts across disciplines varied by source (Bakkalbasi et al., 2006; Yang & Meho, 2006; Kulkarni et al., 2009). It was also shown that, for a small list of researchers, the *h-index* calculated from these three sources gave very different results (Bar-Ilan, 2008). The latest large scale comparison showed that GS had significantly more coverage of citations than WoS and Scopus, though the rank correlations were high (Martín-Martín et al., 2018). Interestingly, Archambault et al. (2009) also showed that the rankings of countries by number of papers and citations were highly correlated between results extracted separately from WoS and Scopus.

Mongeon & Paul-Hus (2016) found that the journal coverages of both WoS and Scopus were biased towards Natural Sciences, Engineering and Biomedical Research. More importantly, their overall coverages differed significantly. Similar findings were obtained by Harzing & Alakangas (2016) when GS was added to the comparison, although for a much smaller sample of objects. Franceschini et al. (2016) also studied database errors in both Scopus and WoS, and found that the distributions of errors were very different between these two sources.

MSA was re-launched (in beta version) in 2016 as the newly improved incarnation of the outdated Microsoft Academic Services. MSA obtains bibliographic data through web pages crawled by Bing. MSA’s emergence and fast growth (at a rate of 1.3 million records per month, according to Hug & Brändle, 2017) has spurred its use in several bibliometrics studies (De Domenico et al., 2016; Portenoy et al., 2016; Sandulescu & Chiru, 2016; Wesley-Smith et al., 2016; Vaccario et al., 2017; Portenoy & West, 2017; Effendy & Yap, 2017). At the same time, various papers have tracked changes in the MSA database and compared it to other bibliographic sources (Paszcza, 2016; Harzing, 2016; Harzing & Alakangas, 2017a; Harzing & Alakangas, 2017b; Hug & Brändle, 2017). Its rapid development, especially in correcting some vital errors, over the past two years and strength in coverage have been very encouraging.

Tsay et al. (2017) indicated that MSA had similar coverage to GS and the Astrophysics Data System for publications of a sample of Physics Nobel Laureates from 2001 to 2013, with MSA having a much lower internal overlap percentage than that of GS. MSA has also recently been used to predict *Article Influence* scores for open access (OA) journals (Norlander et al., 2018). Hug et al. (2017) and Thelwall (2018), using samples of publications, showed there was uniformity between citation analyses done via MSA and Scopus. Harzing & Alakangas (2017a) also showed, for individual researchers, that the citation counts by MSA were similar to or higher than Scopus and WoS, varying across disciplines.

### 1.3 What is different in this study?

As discussed by Neylon & Wu (2009), using a singular article-level or journal-level metric as a filter for scientific literature is deeply flawed and incorporating diverse effective measurement tools is a necessary practice. In a similar vein, using a single bibliographic source for evaluating specific aspects of academia can be very misleading. Given the immense social and academic impacts of the results of such evaluations, and the unlikeliness of them (as either part of research quantification or rankings) being completely discarded anytime soon, one ought to be cautious in both interpreting and constructing such evaluation frameworks. With this in mind, we aim to provide a deep exploration in comparing the coverage of research objects with DOIs (digital object identifiers) in WoS, Scopus and MSA^3^, in terms of both volume and various bibliographic variables, at the institutional level. In particular, a sample list of fifteen universities is selected (ranging in geography, prestige and size) and data affiliated with each university are drawn from all three sources (from 2000 to 2018). Less detailed data are also collected for another 140 universities to be used as a supplementary set where applicable. An automated process is used to compare the coverage of the sources and the discrepancies in publication year recorded. On the other hand, manual online searches were deployed to validate affiliation correctness and plausibility for samples of DOIs. The focus on DOIs also provides broader opportunities for cross-validation of bibliographic variables, such as OA status and document types from Unpaywall^4^, and citations data from OpenCitations^5^. This will assist in further understanding of the differences between these sources and the kind of biases that they may lead to.

Previous studies that compared WoS, Scopus and MSA were limited to publications linked to an individual researcher, a small group of researchers, or one university. These comparisons were also mostly drawn in relation to citation counts. This article extends the literature by expanding the study set to include several universities and drawing institutional comparisons across a larger selection of characteristic and measures. The study further includes analyses of potential effects in the exclusive selection of one source for evaluating a set of bibliographic metrics, i.e, potential effects on the ranking of universities. The use of secondary data sources, i.e., Unpaywall and OpenCitations, to construct metrics for OA and citations is another variation from some of the previous work. This gives standardised contrasting sets of records for comparisons across bibliographic sources and potentially reduces the level of dissimilarity caused by internal bias. The results lead up to the main message that it is essential to integrate diverse data sources in any institutional evaluation framework.

The remainder of this article is structured as follows: Section 2 gives an overview of some global characteristics across the various bibliographic databases. Section 3 provides detailed descriptions of our data collection and manual cross-validation processes. All analyses and results are presented in Section 4. Sections 5 and 6 are discussions on limitations and conclusions, respectively.

## 2. Global comparison of features and characteristics across WoS, Scopus & MSA

WoS and Scopus are both online subscription-based academic indexing services. WoS was originally produced by the Institute for Scientific Information (ISI), but was later acquired by Thomson Reuters, and then Clarivate Analytics (formerly a part of Thomson Reuters). It contains a list of several databases, where access (full or partial) to each depends on the selection of subscription models. The search functionalities can also vary according to which databases are selected (for example, the “Organization-Enhanced” search option is not available when all WoS databases are included). On the other hand, Scopus (provided by Elsevier) seems to offer one unified database of all document types (the only exception is data on patents, which pops up as a separate list in search results). A quick manual online search would reveal a wider variety of document types in WoS. For example, it contains items listed as “poetry”, which does not seem to fit into any of the types in Scopus.

MSA is open to the public through the Academic Knowledge API, though both a rate limit and a monthly usage cap apply to this free version^9^. The subscription version is documented as relatively cheap at $0.32 per 1000 transactions^10^. Its semantic search functionality and ability to cater for natural language queries are amongst the main differences from the other two bibliographic sources. Its coverage in patents has greatly increased through the recent inclusion of Lens.org metadata^11^. As a preliminary examination, we take a look at some global characteristics and features across the three sources. Table 1 provides an overview of coverage and comparative strengths in each source.

**Table 1:** Coverage and features of WoS, Scopus and MSA.

|  | WoS Core | WoS All | Scopus | MSA |
| --- | --- | --- | --- | --- |
| **Subject Focus** | Sciences and Technology | Similar to WoS Core but with significant increases in Social Sciences | All Sciences, with no subject indexing for Arts & Humanities | Sciences, but with significantly more coverage for Social Sciences and Arts & Humanities |
| **Total Count<sup>6</sup>** | 62,602,202 | 73,808,358 | 72,170,639 | 206,252,196 |
| **Time Span<sup>7</sup>** | From 1972 | From 1950 | From 1858 | From 1800 |
| **Coverage<sup>9</sup>** | >20,300 Journals, Books and Conference Proceedings | >34,200 Journals, Books, Proceedings, Patents and Data Sets | 24,130 Journals<br>245 Conference Series | 47,989 Journals<br>4,029 Conference Series<br>Includes books, book chapters and patents |
| **Updating Frequency** | Daily (Mon to Fri) | Ranges from daily to monthly | Daily | Weekly |
| **Strengths** | • Comprehensive search options, including affiliation, DOIs, year, etc.<br>• Organisation-enhanced search<br>• Provides detailed OA information as per Unpaywall | • Provides detailed OA information as per Unpaywall<br>• Provides coverage of patents and data sets.<br>• Increased regional coverage, e.g., Russia, Latin America and China | • Simple API access available through Python<br>• Comprehensive search options, including affiliation, DOI, year, etc. | • API available through languages such as R and Python<br>• A strong coverage of all subject areas<br>• Includes patents data from Lens.org<br>• semantic search |
| **Weaknesses** | • Lack of coverage for Arts & Humanities | • Does not seem to provide affiliation search (and various other queries that are available for WoS Core) | • Fewer details of OA information are provided<sup>8</sup><br>• Little to no coverage of Arts & Humanities | • No apparent record of OA information<br>• Very few search options through the web service<br>• Completeness and accuracy of metadata less studied |

WoS has several databases from which it extracts data. The most commonly used version is WoS Core, which allows for more functionality. On the other hand, WoS All Databases includes all databases listed by WoS (with increased coverage for Social Sciences and local languages, for example), but due to varying levels of availability of information it functionalities are limited, e.g., less search query options. Scopus does not seem to index Art & Humanities, while MSA appears to have significantly more coverage in Social Sciences and Arts & Humanities than WoS Core and Scopus. With higher coverages for journals and conferences, MSA tracks a significantly larger set of records. It is also interesting to note that MSA had approximately 127 million documents only a couple of years ago (Herrmannova & Knoth, 2016).

The annual total numbers^13^ of objects for the various sources from 1970 to 2017 are displayed in Figure 1. In comparison to Jascó (2005), and other studies mentioned earlier, there seems to be significant increases in both Scopus and WoS, in terms of both growth over time and backfilling. However, both sources still have significantly less total counts than that of MSA. The figure also shows a high degree of correlation between Scopus, WoS Core and WoS All. However, this figure does not provide any information on internal or external overlaps across the sources (which we shall explore).

**Figure 1:**
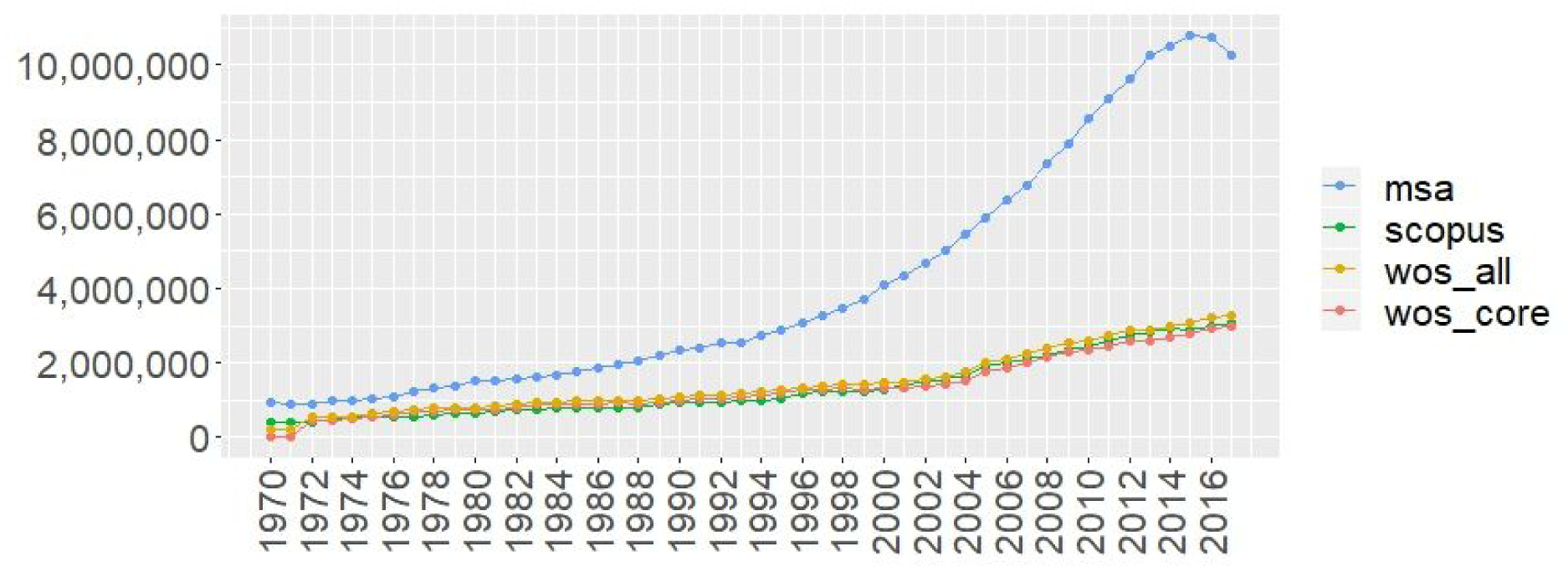
Annual total item counts for Scopus, MSA, WoS Core and WoS All from 1970 to 2017^12^.

To get a better overview of research disciplines covered by each source, the percentage spread of objects across disciplines, for each source, is displayed in Figure 2. Evidently all sources are dominated by the sciences, as commonly noted in the literature. However, MSA does seem to have relatively higher proportions for both Social Sciences and Arts & Humanities.

**Figure 2:**
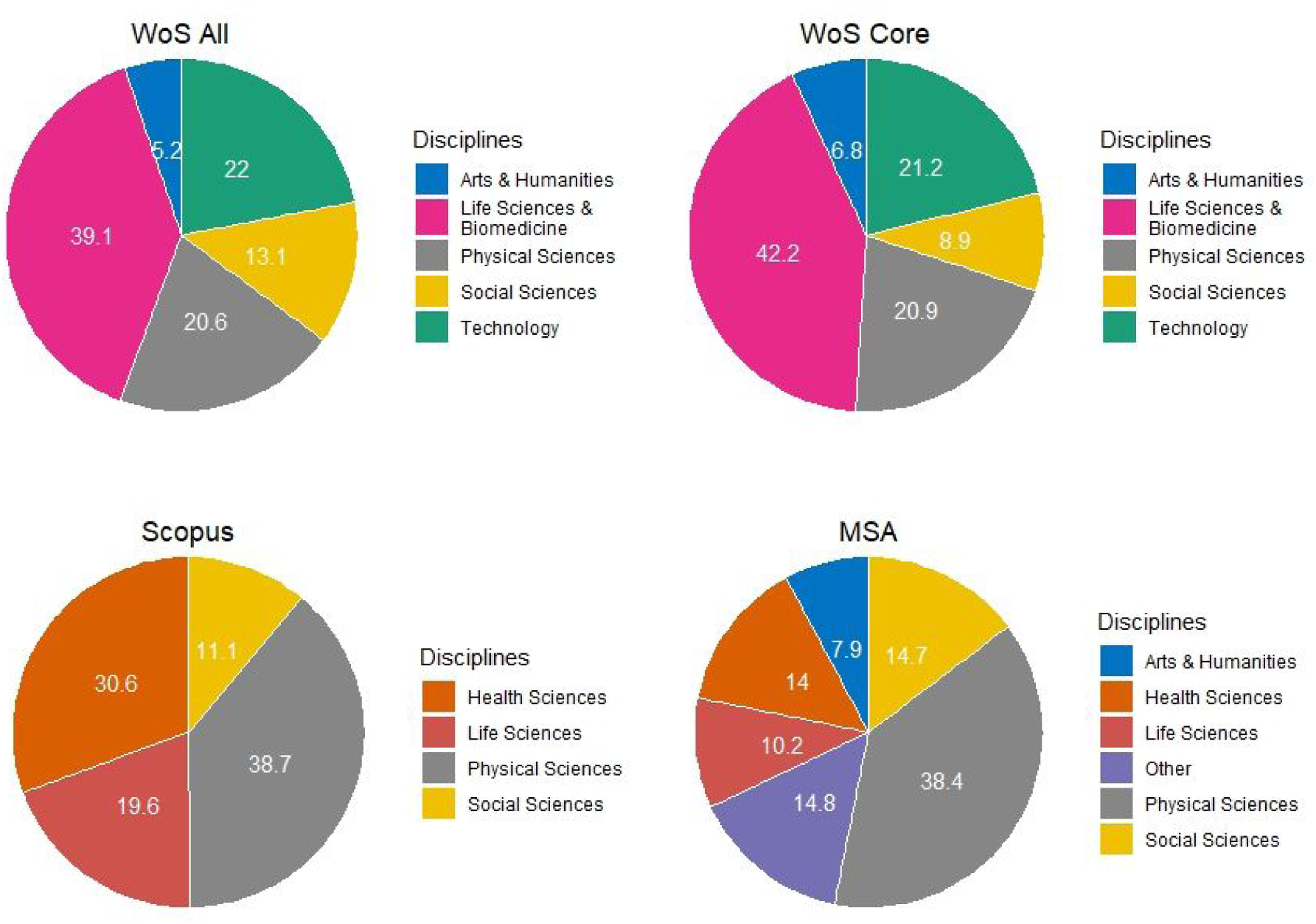
Distributions of objects in WoS All^14^, WoS Core^15^, Scopus^16^ and MSA^17^ among disciplines.

## 3. Methodology & data

To perform a more detailed comparison of sources, we gather output for a selected set of fifteen universities (which range in geography, prestige and size) from each bibliographic source, i.e, WoS, Scopus and MSA. This is done through the use of APIs for each source. We extract records for the years from 2000 to 2018 via affiliation IDs (in the case of Scopus and MSA) and organization-enhanced search terms (for WoS)^18^. The results form 3 sets of data (one from each source) for each university. Subsequently, DOIs of objects (for those that do have them) are extracted from each set. A further 140 universities are also included as a supplementary set to be used where necessary. Our strategy is to explore various bibliographic characteristics related to these DOIs at the overall level (all years and all institutions) and then contrast that with the corresponding results for individual universities focusing on a single year (i.e., 2016). Where applicable, the analysis for 2016 is also extended to the full set of 155 universities. We are mainly interested in the following characteristics:

1. Distributions (e.g., Venn diagrams) of DOIs across sources
2. Discrepancies in publication year recorded by each source
3. Document types across various parts of the Venn diagrams of DOIs
4. Citation counts (as per OpenCitations) calculated across sources
5. OA levels (as per Unpaywall) calculated across sources
6. Plausibility of assigned affiliation for DOIs exclusively indexed by a single source

Characteristics 1 to 5 are mostly automated, with data collected into Google Cloud via APIs of the WoS^19^, Scopus and MSA. Two additional data dumps are also used. These are Unpaywall and OpenCitations data. Unpaywall is used to query the OA status of (Crossref) DOIs and document type. For this article, we only require the general OA status and not the type of OA (e.g., gold OA, green OA, etc.). Hence, we only use the “is_oa” field in the Unpaywall metadata to determine the OA status of DOIs in our data. Document type is determined via the data field “genre”. OpenCitations records citation links between Crossref DOIs. By querying and merging all links to a DOI, it allows us to determine the number of citations this DOI receives. We gather this information for a set of DOIs of interest (e.g., DOIs from WoS affiliated to one university) and obtain total citation counts for this set. This total can then be divided by the number of (Crossref) DOIs affiliated to this university to produce an average citation count^20^.

A manual process is followed for checking characteristic 6. The procedure for the manual validation is focused on the non-overlapping parts of the three sources (i.e., shaded sections in Figure 3). The overlapping parts indicate agreement by at least two sources, over both affiliation and publication year records (when filtered down to a particular year). Given the different ways in which the sources gather data, the reliability of information for these parts is much more convincing. In contrast, the non-overlapping sections are not validated by other sources (at least appears so through the data gathering process). This leads to the need for the manual validation process.

**Figure 3:**
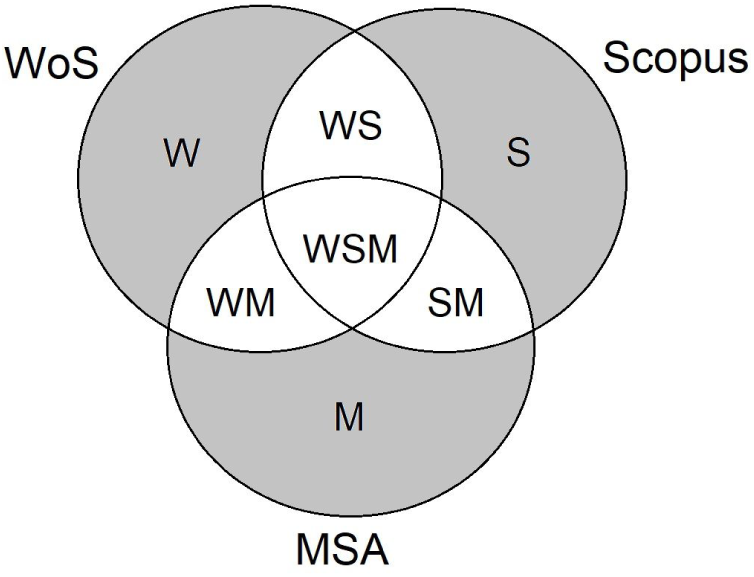
Non-overlapping sections (in grey) of the spread of DOIs from 3 sources for an institution in a particular year.

The publication year can be a reason for the discrepancy of coverage due to inconsistencies in which date is recorded. For example, in the case of a journal article, a source may choose to record either the date of the journal issue, publication date for the article, or the date for which the article first appears online. Hence, our first step is to check whether DOIs from the non-overlapping sections are indeed in another source but fall in a different year. After removing these DOIs that were identified via comparison to adjacent years, we sample the remaining DOIs from each non-overlapping section for manual validation (Figure 3). This is processed for DOIs from 2016.

The process that leads to the manual validation is summarised in the flowchart given in Figure 4^21^. Once DOIs are sampled from each non-overlapping section, they are compared against the other two sources (via DOI and title searches on each source’s webpage) and also the original document (online versions)^22^.

**Figure 4:**
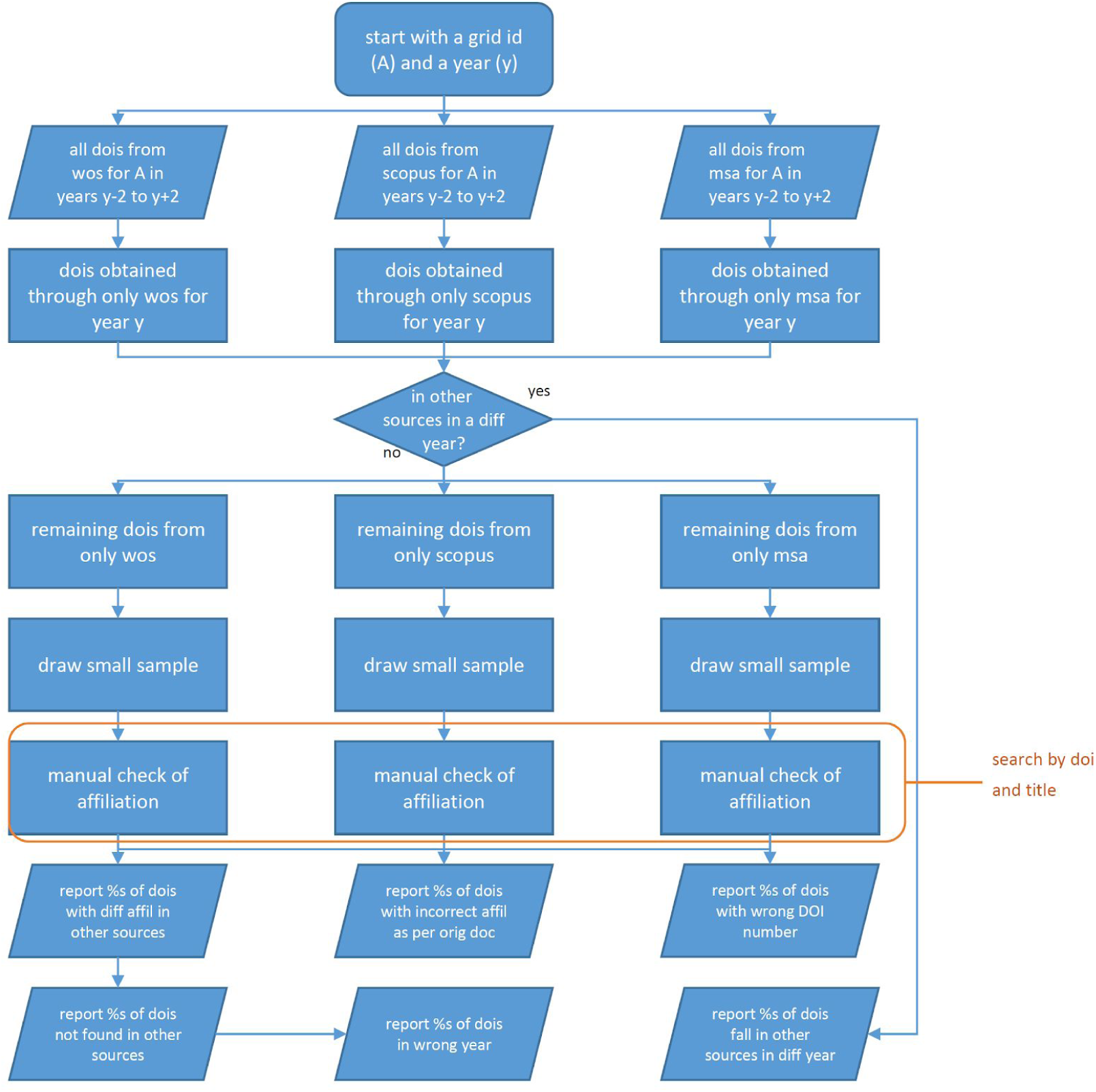
Flowchart of the process leading to manual validation.

Assume we have three sources A, B and C, and the current set of DOIs are from source A. The following questions are asked as part of the manual checking process (with a likewise procedure used for DOIs from the other two sources):

1. *Is this DOI found in the metadata record in source B?*
2. *Is the title associated with this DOI found in source B?*
3. *Is the exact affiliation phrase found in the metadata record in source B?*
4. *If not, is the affiliation plausible?*
5. *Is this DOI found in the metadata record in source C?*
6. *Is the title associated with this DOI found in source C?*
7. *Is the exact affiliation phrase found in the metadata record in source C?*
8. *If not, is the affiliation plausible?*
9. *Is the DOI correctly recorded in source A (as per original document or doi.org)?*
10. *Is the exact affiliation phrase found on the original document?*
11. *If not, is the affiliation plausible?*

The numbers of DOIs to be sampled for each institution are 30, 30 and 40 from (exclusively) WoS, Scopus and MSA, respectively, after removal of DOIs that are found in another source for a different year.

Table 2 presents the total number of unique DOI records we have obtained from each source, the combined number of unique DOIs and how many of these DOIs are recorded in Unpaywall, for each institution for 2016. The coverage of DOIs by Unpaywall is very high, as expected. The only slight exception is DUT, where a significantly higher portion of Scopus DOIs were not recorded by Unpaywall. A quick exploration^23^ finds most of these DOIs to be registered with China National Knowledge Infrastructure (CNKI) or the Institute of Scientific and Technical Information of China (ISTIC), whereas Unpaywall currently only index Crossref DOIs^24^.

**Table 2:** The spread of DOIs (for 2016) across various sources for 15 different institutions and percentages of them recorded in Unpaywall.

|  | WoS |  | Scopus |  | MSA |  | Combined |  |
| --- | --- | --- | --- | --- | --- | --- | --- | --- |
| Instituion<sup>25</sup> | Total Count | Unpayw all<sup>26</sup> | Total Count | Unpayw all | Total Count | Unpayw all | Total Count | Unpayw all |
| Cairo | 2629 | 98.4% | 2454 | 98.7% | 2761 | 98.7% | 3793 | 97.4% |
| Curtin | 3168 | 99.2% | 2963 | 98.8% | 3150 | 98.8% | 4070 | 98.2% |
| DUT | 3552 | 97.9% | 3789 | 82.8% | 3582 | 99.5% | 5091 | 86.1% |
| IISC | 2002 | 98.6% | 1964 | 98.9% | 2464 | 97.8% | 3156 | 97.8% |
| ITB | 585 | 99.1% | 1040 | 99.7% | 1027 | 96.8% | 1744 | 97.8% |
| LU | 1419 | 97.8% | 1357 | 98.6% | 1564 | 99.2% | 2014 | 97.5% |
| MIT | 8053 | 99.3% | 6702 | 99.1% | 7457 | 97.6% | 10889 | 98.6% |
| MSU | 5480 | 99.2% | 5107 | 99.5% | 5719 | 99.3% | 7362 | 98.8% |
| UNAM | 4754 | 98.6% | 4258 | 98.8% | 5401 | 96.8% | 7056 | 96.5% |
| UCL | 13615 | 99.0% | 11255 | 99.1% | 9924 | 98.7% | 17230 | 98.4% |
| UCT | 3079 | 98.8% | 2852 | 98.8% | 3189 | 98.3% | 4206 | 97.9% |
| Giessen | 1871 | 98.7% | 1638 | 99.4% | 1545 | 99.2% | 2354 | 98.2% |
| USP | 11451 | 98.3% | 10923 | 99.4% | 13664 | 96.8% | 17732 | 96.5% |
| Tokyo | 9640 | 99.0% | 8810 | 99.1% | 9789 | 97.9% | 12848 | 97.9% |
| WSU | 2717 | 98.7% | 2041 | 99.2% | 2794 | 99.1% | 3569 | 98.2% |
| Overall<sup>27</sup> | 71709 | 98.8% | 65017 | 98.2% | 72386 | 98.5% | 100456 | 97.2% |

## 4. Analysis & discussion

In this section, we proceed with the comparisons across sources. We will start with exploring the coverage of DOIs by each source. This is followed by examining the amount of agreement, or disagreement, of publication year recorded by each bibliographic source. The document types, citation counts and OA percentages, as per source, are the subsequent analyses. Lastly, a manual cross-validation procedure is employed for samples extracted from non-overlapping sections of the Venn diagrams for each institution in our sample of 15 institutions.

### 4.1 Coverage and distribution of DOIs

Here we take an exploration of the spread of the DOIs across the sources. Figure 5 shows the Venn diagrams of DOI counts for our initial set of 15 universities combined from 2000 to 2018 and for just 2016^28^, respectively. Evidently, the central regions (overlap of all three sources) have the highest count in each Venn diagram. These are DOIs that have been indexed by all three sources and, given the intended global coverage of these sources, the relatively higher counts here are not at all surprising. However, there are also significant portions of DOIs exclusively accessed via a single source in both Venn diagrams. This gives rise to the potential biases in any bibliometric measure to be calculated from a single source.

**Figure 5:**
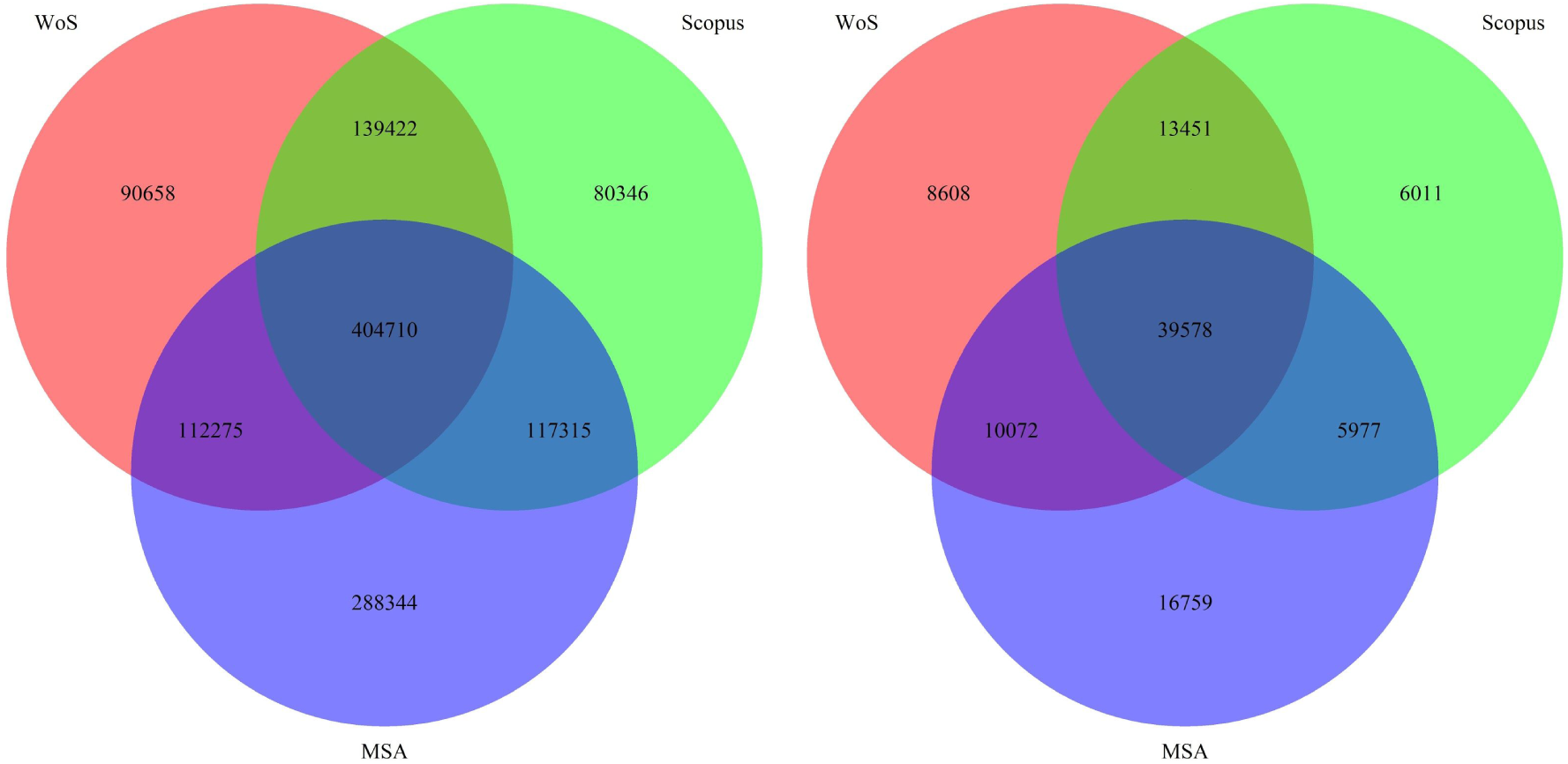
Venn diagrams of DOIs from all 15 institutions for years 2000-2018 (left) and for only 2016 (right).

This pattern of difference in coverage is mirrored at the institutional level. Appendix 3 contains two Venn diagrams for each institution, both for 2016. In each case, the Venn diagram on the left records all DOIs as per bibliographic source and the one on the right is a subset of these DOIs that are also in the Unpaywall database. It is noted that the two Venn diagrams for each institution are quite similar due to the high coverage of these DOIs by Unpaywall. The only exception being the Scopus coverage of DOIs for DUT, for which the DOIs exclusively indexed by Scopus significantly decreased when moving from the left Venn diagram to the one on the right. This is consistent with what we observed earlier, with many these DOIs (provided by agencies other than Crossref) not indexed in Unpaywall. The overall pattern is that there appears to be significant portions of DOIs only indexed by a single source. Hence, pulling together these sources can greatly enhance coverage. Interestingly, for most institutions, MSA has the most number of exclusively indexed DOIs. The only exception being UCL.

To have a better overview of how coverages of these three sources vary across institutions, we perform several analyses as follows. First, we identify each institution with the seven different counts as per its own Venn diagram of all DOIs (the Venn diagrams on the left in Appendix 3). We also include another 140^29^ universities for comparison. We view each (GRID ID, DOI) pair as a distinct object. Hence, we obtain a 155 by 7 contingency table. Each column of this table represents the number of DOIs falling in the respective section of the Venn diagram, e.g., column 1 is the number of DOIs in section WSM of the Venn diagram (refer to Figure 3). We can also convert these counts to proportions through dividing them by the total number of DOIs for each institution. Figure 6 shows the distribution of these proportions for each section of the Venn diagram. The higher proportion in the central region (section WSM) of the Venn diagram is again observed. The general pattern having emerged is that, for all sections of the Venn diagram, there appears to be a concentrated central location with many extreme cases (excess kurtosis of 2.29, 9.72, 5.96, 1.82, 22.24, 11.49 and 6.88, from sections WSM, WS, WM, SM, W, S and M, respectively) and substantial skewness.

**Figure 6:**
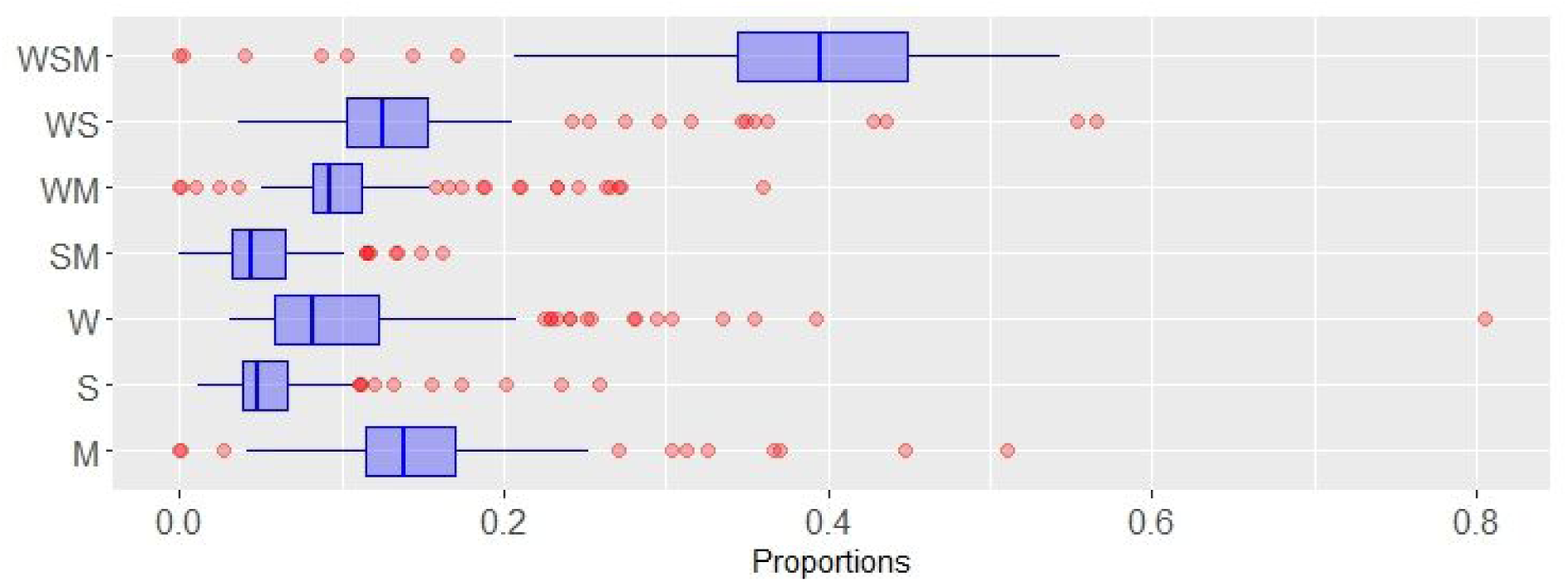
Boxplots of proportions of DOIs that fall in each section of the Venn diagram across 155 universities for 2016.

We can also concatenate the respective sections to get the proportion of DOIs covered by each bibliographic source. The spreads of these proportions are summarised in Figure 7 as histograms.

**Figure 7:**
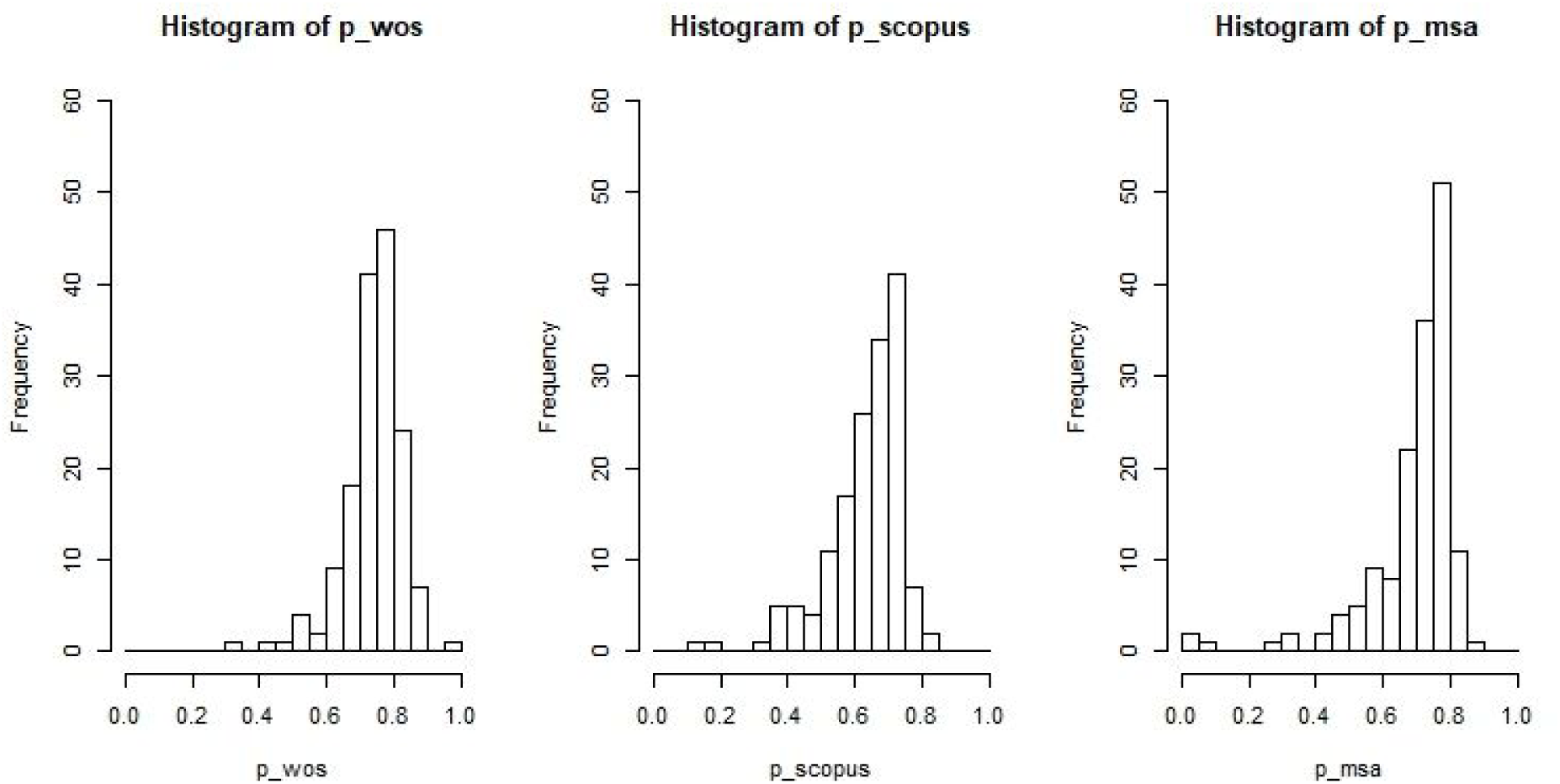
Histograms of proportions of DOIs in WoS, Scopus and MSA for 2016 (across 155 universities).

Again, the pattern of high central peak, skewness and heavy tails are observed. The peakedness and heavy tails are confirmed by the excess kurtosis of 4.29, 3.34 and 8.60 for WoS, Scopus and MSA respectively. The skewness to the left with number of extreme cases highlights the low degree of coverage for some universities. Meanwhile, a correlation analysis of the proportions for the three sources is quite intriguing (see Table 3). Both Spearman’s rank correlation and Pearson’s correlation matrices are presented here. There appears to be a negative correlation between coverage by WoS and coverage by MSA, i.e., when there is a high proportion of coverage by WoS, the coverage by MSA is relatively low. There is also a low correlation between WoS and Scopus. While much of these may be attributed to the different methodological structure and focus across WoS, Scopus and MSA, the degree of non-alignments is still quite a surprise.^30^

**Table 3:** Spearman’s rank correlation and Pearson’s correlation matrices of proportions of DOIs covered by each bibliographic source.

|  | Spearman |  |  | Pearson |  |  |
| --- | --- | --- | --- | --- | --- | --- |
| Sources | p_wos | p_scopus | p_msa | p_wos | p_scopus | p_msa |
| p_wos | 1 | 0.07 | -0.50 | 1 | 0.08 | -0.39 |
| p_scopus | 0.07 | 1 | 0.10 | 0.08 | 1 | 0.06 |
| p_msa | -0.50 | 0.10 | 1 | -0.39 | 0.06 | 1 |

We further perform tests of homogeneity across institutions to check whether the spread of DOIs across individual Venn diagrams come from the same probability distribution. The results of these tests are provided in Table 4. It is evidenced that the chance of rejecting homogeneity is very high. Bootstrapped samples from sample sizes 10 to 155, in increments of 5, all gave similar results as well.

**Table 4:** P-values for tests of homogeneity across institutions in terms of the distribution of DOIs.

| Sample | Chi square<sup>31</sup> | Chi square MC<sup>32</sup> | G test |
| --- | --- | --- | --- |
| 15 universities | <0.0001 | 0.0002 | <0.0001 |
| 155 universities | <0.0001 | 0.0002 | <0.0001 |

It is also expected that these Venn diagrams are not symmetrical (in the sense of equal coverage by each source), which is observable from the Venn diagrams of our initial sample of 15 universities in Appendix 3. However, to obtain further insight into the symmetry of a large number of Venn diagrams (i.e., all 155 universities), we introduce 3 related measures. Let *p_i_* be the proportion of DOIs that fall in part *i* of a Venn diagram and define the following three measures:

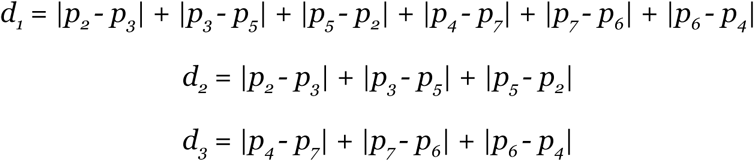

where *d_1_* is the sum of absolute differences across the whole Venn diagram, *d_2_* is sum of inner differences and *d_3_* is sum of differences across the outer regions of the Venn diagram. We calculate values for these three measures for each university’s Venn diagram and compare their distributions to those produced by randomly generated Venn diagrams. Firstly, they are compared to randomly generated symmetrical Venn diagrams^33^. The resulting distributions are presented in Figure 8. It is quite obvious that the results from our data do not correspond to those of generated symmetrical Venn diagrams. As further contrasts, we also compare these measures against Venn diagrams generated from various other distributions (see Appendix 5). As expected, our data is better represented by other distributions rather than that produced by symmetrical Venn diagrams. Furthermore, there appears to be some differences in distributions across *d_1_*, *d_2_* and *d_3_*, which we do not further examine and leave for future exploration.

**Figure 8:**
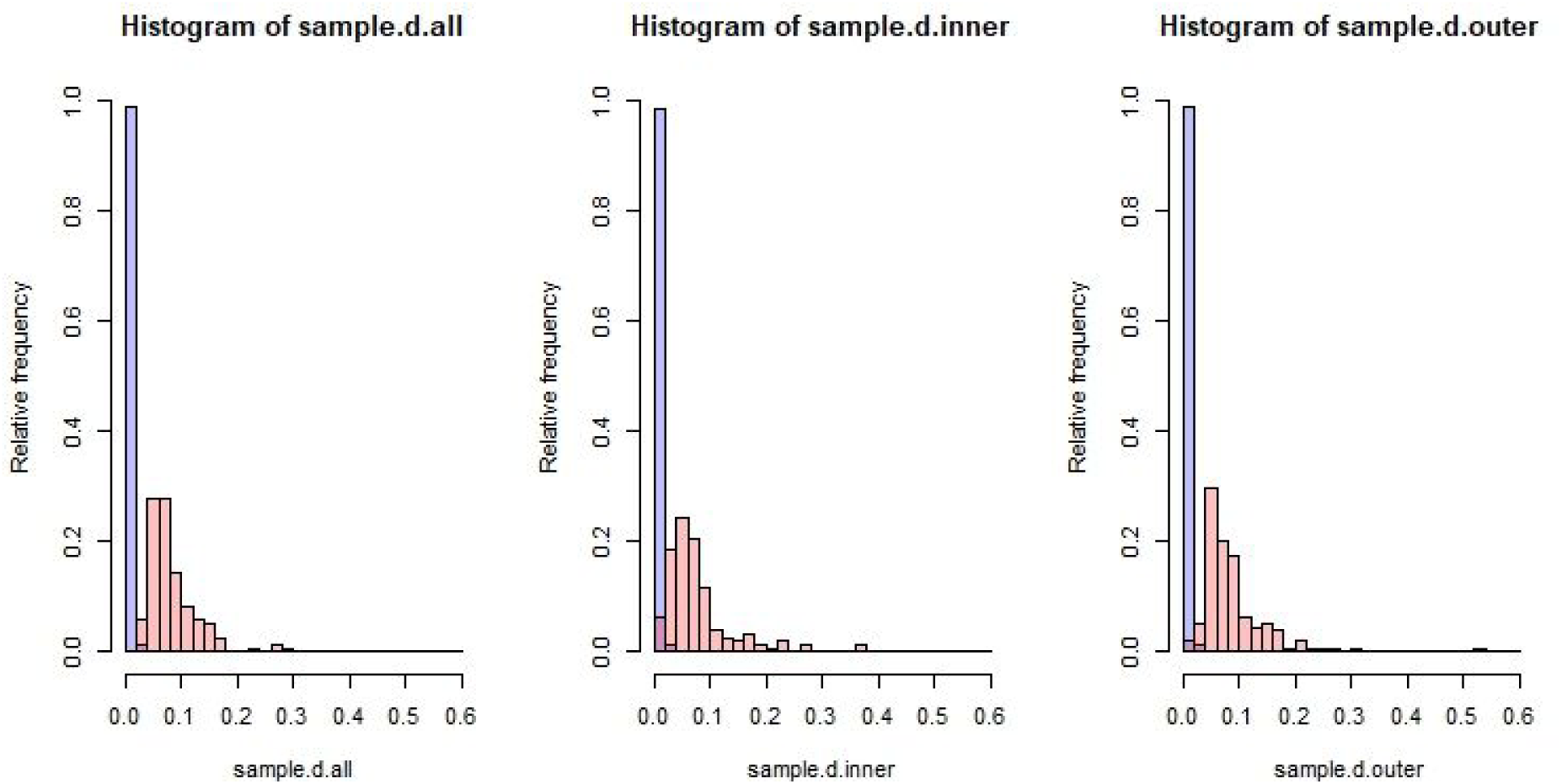
Histograms of ***d_1_***, ***d_2_*** and ***d_3_*** (left to right, respectively) for our data of 156 universities (in red) and for randomly generated symmetrical Venn diagrams (in purple).

Now that we have confirmed the differences in DOI distributions across institutions and negative to low correlations between the non-symmetrical coverages by the three bibliographic sources, a follow-up question may be whether there are groupings amongst these universities. We proceed with a hierarchical cluster analysis for both the sample of 15 universities and for all 155 universities, using dissimilarities between the proportions of the

Venn diagrams as clustering criteria^34^. At the same time, we also colour code the universities by their regions and rank positions on the 2019 THE World University Rankings. Some of these are presented in Appendix 6. While no striking patterns emerge, there does appear to be some interesting groupings. For example, there seems to be a block of European and American universities towards the left of the dendrogram coloured by region. Perhaps unsurprisingly, around the same area for the dendrogram coloured by THE ranking, there is also a rough cluster of the most highly ranked universities.

The contrasts may be more apparent for the smaller sample of 15 universities. An example of this is presented in Figure 9. ITB is clearly an outlier from the rest of the group (we shall come across this again later) and the two highest ranked universities are placed quite close to each other. Seven of the universities ranking from 201 and above are placed on the right of the dendrogram (perhaps in 2 clusters). One of these also consists mainly of universities from non-English speaking regions (Loughborough being the exception). In general, there appear to be some general patterns of prestige and regional clustering. However, we may need a bigger set of universities for a full analysis.

**Figure 9:**
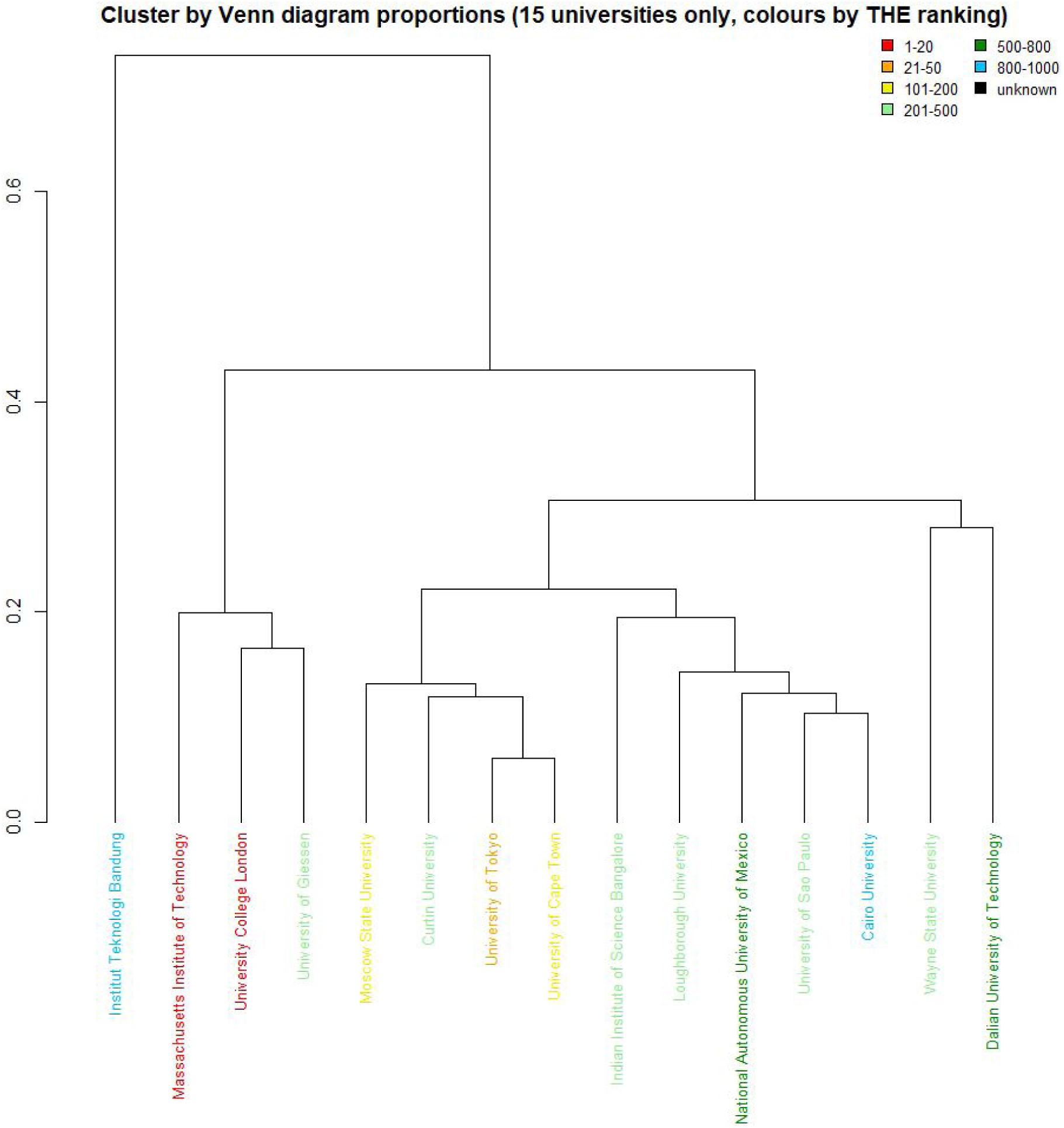
Dendrogram showing clustering of 15 universities by Venn diagram proportions vs rank position on 2019 THE World Universities Rankings.

### 4.2 Comparison of publication years

As mentioned earlier, discrepancies in publication year recorded by different bibliographic sources is possible, given there is no universal standard to the definition of *publication year* (or *publication date* for that matter). It could potentially refer to various dates linked to a research output. This poses a problem when one would like to combine sources to evaluate and track a bibliometric variable (or metric) over time. If not dealt with, a DOI can be double-counted, i.e., counted two or more times in different years via different sources. In the following, we explore the amount of agreement (or disagreement) on publication years by WoS, Scopus and MSA. The overall numbers are presented in Table 5, covering all DOIs for 15 institutions and years from 2000 to 2018.

**Table 5:** The amount of agreements of “publication year” across bibliographic sources for 15 universities combined (from 2000 to 2018).

| Sources | WoS/Scopus | WoS/MSA | Scopus/MSA | All 3 sources |
| --- | --- | --- | --- | --- |
| Total agreement on year | 541501 | 508269 | 509797 | 397560 |
| Total overlap of DOIs<sup>35</sup> | 544133 | 516986 | 522026 | 404710 |
| % of agreement | 99.5% | 98.3% | 97.7% | 98.2% |

In this table, the number of DOIs jointly indexed by pairs of bibliographic sources (columns 2 to 4) and by all three bibliographic sources (column 5) are recorded (row 3). The corresponding numbers and percentages of DOIs for which the sources agree on publication years are given in rows 2 and 4, respectively. It should be noted that these percentages are calculated over different sets of DOIs (i.e., different denominators). For example, number of DOIs common to all three sources (i.e., 404710) is less than number of DOIs common to only Scopus and MSA (i.e., 522026).

It is clear that the overall levels of agreements are very high. However, two follow-up questions are: 1) for DOIs that does lie in a different source for a different year, what is the spread of these DOIs over years? 2) while the overall agreement of publication years is high, does that carry over to individual institutions?

To answer these questions, we now focus our attention to the year 2016 and DOIs that are exclusively indexed by a single source for that year. Figure 10 displays the spread of such DOIs from a particular sources when matched against the other sources for different years. These are again DOIs from our sample of 15 institutions combined. The majority of the discrepancies are within one year (i.e., falling in 2015 and 2017), while going a further one year period in both directions covers almost all remaining cases. We also note some differences across the sources. The amount of discrepancies between WoS and Scopus are relatively smaller compared to those involving MSA. This may be the likely result of MSA defaulting their record of publication date to when a document first appears online^36^.

**Figure 10:**
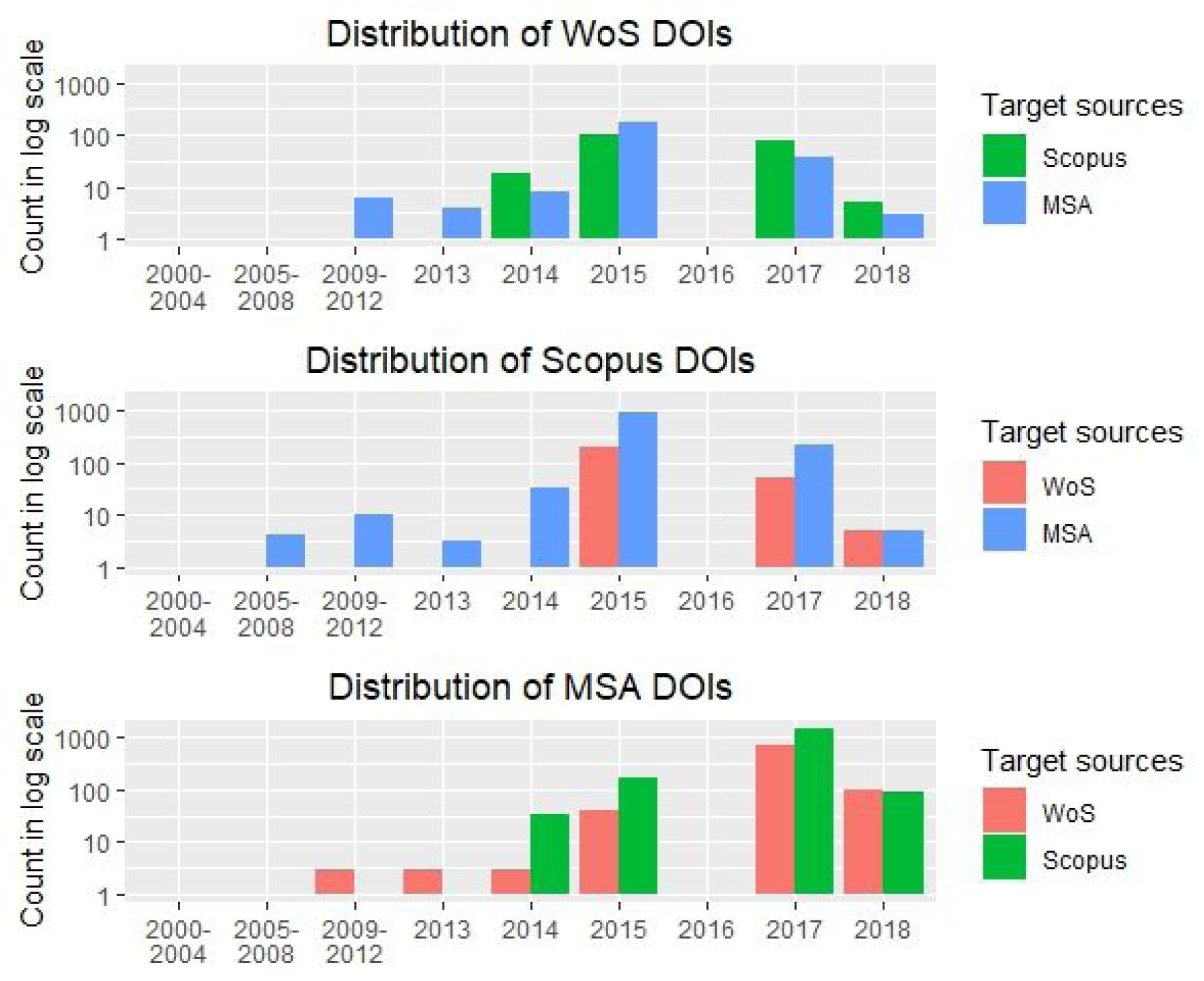
Number (in log scale) of 2016 DOIs from each source (exclusively) that falls in another source but in a different year (15 universities combined).

Next we explore how these discrepancies of the publication year are distributed for individual institutions. Table 6 records, for each source, the percentages of DOIs from 2016 that lies in other two sources but differ by a year and two years, respectively. For WoS, the percentage of matches over one year is consistently small for all institutions, ranging from 0.8% to 2%. This also significantly decreases when moving to the two year gap. In contrast, Scopus and MSA seem to have more varied results for the one year gap across institutions and with generally higher percentages than those of WoS.

**Table 6:** Percentage of 2016 DOIs^37^, per bibliographic source, listed in the other two sources but a year away (i.e., 2015 and 2017) and two years away (i.e., 2014 and 2018).

|  | All DOIs from WoS vs other two sources |  | All DOIs from Scopus vs other two sources |  | All DOIs from MSA vs other two sources |  |
| --- | --- | --- | --- | --- | --- | --- |
| Institution | 1 year | 2 years | 1 year | 2 years | 1 year | 2 years |
| Cairo | 1.5 | 0.2 | 6.0 | 0.2 | 5.4 | 0.4 |
| Curtin | 1.2 | 0.2 | 2.3 | 0.0 | 2.6 | 0.3 |
| DUT | 1.3 | 0.1 | 2.5 | 0.1 | 3.1 | 0.3 |
| IISC | 1.3 | 0.2 | 4.9 | 0.2 | 6.0 | 0.1 |
| ITB | 1.0 | 0.0 | 24.6 | 0.0 | 25.6 | 0.0 |
| LU | 1.6 | 0.1 | 2.7 | 0.1 | 3.4 | 0.3 |
| MIT | 1.2 | 0.2 | 1.6 | 0.1 | 2.6 | 0.4 |
| MSU | 0.8 | 0.1 | 0.7 | 0.1 | 1.3 | 0.1 |
| UNAM | 1.5 | 0.1 | 1.5 | 0.0 | 1.7 | 0.2 |
| UCL | 0.8 | 0.1 | 1.2 | 0.2 | 2.3 | 0.3 |
| UCT | 1.8 | 0.1 | 2.0 | 0.1 | 2.8 | 0.1 |
| Giessen | 1.1 | 0.1 | 1.4 | 0.4 | 1.8 | 0.1 |
| USP | 1.5 | 0.1 | 2.2 | 0.1 | 2.3 | 0.2 |
| Tokyo | 1.2 | 0.2 | 2.1 | 0.1 | 2.7 | 0.3 |
| WSU | 2.0 | 0.2 | 2.4 | 0.2 | 2.1 | 0.2 |

The one standout case is ITB, an Indonesian university situated in the City of Bandung. Its results for WoS is similar to other institutions, but one year comparisons from Scopus and MSA yielded 24.6% and 25.6%, respectively. We believe that this may be due to two reasons. Firstly, WoS has a significantly less coverage for ITB (see Venn diagrams for ITB in Appendix 3) than those of Scopus and MSA. There is also a much lower number of DOIs exclusively indexed by WoS. Secondly, Indonesia has an extraordinary large number of local journals owned by universities and many of them OA (with or without OA license). This is largely driven by government policy that requires academics and students to publish research results and theses in academic journals^38^. Many of these journals are also linked to conference output. This may have resulted in a systematic difference on how publication years (or dates) are recorded (or defined). The other two cases that stand out, although less extreme, are Cairo and IISC.

In Appendix 7, the directions of the comparisons are displayed in more detail for the three standout cases (i.e., Cairo, IISC and ITB). The comparisons are also narrowed down to just Scopus and MSA. It is immediately clear that the difference between Scopus and MSA are the main contributors to these standout cases. Also, it appears that MSA tends to record the publication year once year earlier than Scopus. This is in line with our earlier comments regarding MSA recording date of first online and the publishing venues in Indonesia.

Let us now focus on the outer parts of the Venn diagrams (i.e., DOIs that appear to be exclusively indexed by a single source). Results for these sets of DOIs are presented in Table 7. Columns 2, 5 and 8 lists the number of 2016 DOIs exclusively indexed by WoS, Scopus and MSA (compare these again with Venn diagrams in Appendix 3), without checking against DOIs listed in other years. The subsequent columns list the percentages of these DOIs that can be matched against DOIs in other sources for a one year and a two year gap, respectively. Consistent with Table 6, significantly higher portions of DOI matches occur after incorporating the first one year gap, as compared to including a further one year on both sides. Relatively, the most impacted university is ITB, which corresponds to the observation made in Table 6. In general, the effect on these exclusive sets of DOIs varies considerably across institutions and sources (more so than observed in Table 6, as expected).

**Table 7:** Percentages of 2016 DOIs exclusively from each bibliographic source that is indexed by other sources within 1 and 2 year (before and after 2016) gaps, respectively.

|  | DOIs excl. from WoS |  |  | DOIs excl. from Scopus |  |  | DOIs excl. from MSA |  |  |
| --- | --- | --- | --- | --- | --- | --- | --- | --- | --- |
| Institution | original | 1<sup>39</sup><br>year | 2<sup>40</sup><br>year | original | 1<br>year | 2<br>year | original | 1<br>year | 2<br>year |
| Cairo | 340 | 2.4 | 0.3 | 261 | 47.9 | 0.0 | 660 | 21.2 | 1.8 |
| Curtin | 220 | 5.9 | 2.3 | 198 | 28.8 | 0.5 | 502 | 13.3 | 1.4 |
| DUT | 187 | 8.6 | 0.5 | 794 | 9.4 | 0.0 | 533 | 18.4 | 1.9 |
| IISC | 126 | 6.3 | 0.8 | 177 | 49.2 | 0.0 | 712 | 19.2 | 0.3 |
| ITB | 61 | 3.3 | 0.0 | 410 | 62.4 | 0.0 | 545 | 47.5 | 0.0 |
| LU | 127 | 4.7 | 0.0 | 138 | 18.8 | 0.0 | 307 | 15.0 | 1.6 |
| MIT | 1309 | 2.8 | 0.3 | 396 | 20.2 | 0.8 | 1784 | 8.8 | 1.6 |
| MSU | 530 | 3.0 | 0.2 | 206 | 8.3 | 0.5 | 1148 | 5.7 | 0.6 |
| UNAM | 522 | 6.5 | 0.4 | 460 | 8.7 | 0.2 | 1532 | 5.2 | 0.8 |
| UCL | 2680 | 1.7 | 0.2 | 1193 | 6.8 | 0.8 | 1735 | 11.4 | 1.4 |
| UCT | 252 | 6.0 | 0.4 | 202 | 12.9 | 1.0 | 681 | 11.6 | 0.3 |
| Giessen | 234 | 2.6 | 0.0 | 110 | 11.8 | 3.6 | 303 | 7.6 | 0.7 |
| USP | 1258 | 4.2 | 0.5 | 932 | 15.9 | 0.5 | 4067 | 7.1 | 0.6 |
| Tokyo | 732 | 8.9 | 0.1 | 565 | 26.7 | 1.4 | 1793 | 13.2 | 0.8 |
| WSU | 265 | 7.9 | 1.1 | 122 | 23.0 | 2.5 | 595 | 8.1 | 0.8 |

### 4.3 Document Types

Another important bibliographic variable is the document types (e.g., journal articles, proceedings, book chapters, etc) that relate to each DOI. In particular, the coverage of different document types can lead to insights into potential disciplinary biases in data sources and differences in institutional focuses on output types.

For this study, we use the “genre” variable in Unpaywall metadata to determine the document type of each DOI. These are Crossref-reported types for all DOI objects in the Crossref database^41^. Table 8 provides the counts of each document type within each part of the Venn diagram between WoS, Scopus and MSA (for all 15 institutions from 2000 to 2018 combined)^42^. An immediate observation is that journal articles make up (by far) the highest proportion of the DOIs. This is true overall and for individual parts of the Venn diagram, but not unexpectedly so. The scenario is again more interesting when we consider the outer parts of the Venn diagram (sections W, S and M of the Venn diagram). The set of DOIs exclusive to MSA contains significantly more book chapters and proceeding-papers relative to any other parts. It also provides almost all thesis entries in our data and is the only source to provide posted-contents. On the other hand, Scopus seems to provide many books and monographs not indexed by the other two sources.

**Table 8:** Document types of all DOIs, recorded in Unpaywall, for all 15 universities combined from 2000 to 2018.

| All 15 universities combined |  |  |  |  |  |  |  |
| --- | --- | --- | --- | --- | --- | --- | --- |
| Section of Venn diag.<br><sup>43</sup> | WSM | SM | WM | M | WS | S | W |
| book-chapter | 241 | 5745 | 4699 | 37138 | 50 | 6709 | 1523 |
| journal-article | 393524 | 96600 | 95127 | 190768 | 144634 | 63244 | 85850 |
| proceedings-article | 1849 | 12172 | 6201 | 61546 | 1504 | 4766 | 3270 |
| reference-entry | 191 | 96 | 78 | 1107 | 89 | 122 | 252 |
| report | 1 | 2 | NA | 216 | NA | 8 | 7 |
| book | NA | 272 | NA | 514 | NA | 4189 | 29 |
| component | NA | 2 | NA | 106 | 2 | 59 | 37 |
| journal | NA | 7 | NA | 9 | 1 | 8 | NA |
| journal-issue | NA | 1 | 3 | 200 | 9 | 20 | 26 |
| monograph | NA | 64 | NA | 144 | NA | 505 | 1 |
| other | NA | 14 | 1 | 36 | NA | 220 | 64 |
| dataset | NA | NA | NA | 7 | NA | NA | 2 |
| dissertation | NA | NA | NA | 341 | NA | 5 | NA |
| posted-content | NA | NA | NA | 1391 | NA | NA | NA |
| reference-book | NA | NA | NA | 1 | NA | 26 | 1 |
| report-series | NA | NA | NA | 116 | NA | NA | NA |
| book-section | NA | NA | NA | NA | NA | 4 | NA |
| book-set | NA | NA | NA | NA | NA | 3 | NA |
| proceedings | NA | NA | NA | NA | NA | NA | 1 |

<sup>42</sup> Note here the total number of DOIs are slightly lower in each part of the Venn diagram as compared to the left Venn diagram in Figure 5. This is because here we are only including DOIs that are also recorded in Unpaywall.
<sup>43</sup> See Figure 3 for the labelling of the Venn diagram.

Again we would like to examine how the situation plays out for individual institutions. After filtering the sets of DOIs to each institution and to the year 2016, we follow the same procedure as above to produce the spread of document types across each part of an institution’s Venn diagram. These are recorded in Appendix 8. As we have observed for the combined data set, journal articles make up the highest portion of the DOIs for each institution. The next two most common document types are book chapters and proceeding papers. The only exception being ITB, where there are slightly more proceeding papers than journal articles. Interestingly, there are a few universities with more book chapters than proceeding papers, i.e., Curtin, UNAM, UCL, UCT, Giessen and WSU.

There are high proportions of book chapters indexed exclusively by MSA for all institutions. MSA also have the highest proportion of exclusively indexed journal articles, except for MIT, UCL and Giessen (WoS has the highest such proportion for these three institutions). It is also observed that MSA and Scopus seem to bring in more additional proceeding papers than WoS (only exception being UNAM where all three sources have similar exclusive coverage on proceeding papers). Scopus also seem to often add books and monographs not indexed by the other two sources. For all universities, journal articles make up the majority of DOIs exclusively indexed by WoS. In contrast, the document types of DOIs exclusively indexed by Scopus or MSA are more diverse. Overall, we observe that each source has a different exclusive coverage of document types and this coverage also varies across institutions.

### 4.4 Citation counts

One set of commonly used bibliographic metrics, in the evaluation of academic output, are those that relate to citation counts. These include metrics such as h-index, impact factor and eigenfactor. However, these citation metrics can also be calculated via different sources. WoS, Scopus and MSA all record and maintain their own citation data. While some research have shown that the citation counts across these sources showed high correlations at the author level and journal level (Harzing, 2016; Harzing & Alakangas, 2017a; Harzing & Alakangas, 2017b), the corresponding effects on a set of universities remain relatively unknown. These analyses were also performed using internal citation counts of each source. In this study, we rather prefer to use a standard set of citation links to be applied to all three sources of DOIs. As such, we introduce a further reference set of data from OpenCitations. We match each DOI against the list of DOI citation links in OpenCitations and obtain (if exist) its total citation count. In Table 9, we present the results combining DOIs for our initial set of 15 universities and for all years from 2000 to 2018.

**Table 9:** Total citations for all 15 institutions from 2000 to 2018, as per OpenCitations.

|  | WoS | Scopus | MSA | Combined |
| --- | --- | --- | --- | --- |
| **Number of DOIs<sup>44</sup>** | 735832 | 734515 | 907239 | 1202032 |
| **Total citations<sup>45</sup>** | 9670953 | 9581710 | 9122420 | 13060486 |
| **Citations per output** | 13.1 | 13.0 | 10.1 | 10.9 |

The results show that the total number of citations to MSA DOIs are slightly lower than WoS and Scopus. This is in addition to an already larger set of (Unpaywall/Crossref) DOIs. Hence, MSA resulted in a lower average citation number from the OpenCitations citation links.

As a further analysis, we would like to investigate how the change of bibliographic source influences the perceived performance of an institution. Figure 11 presents two different charts for total citations and ranks by average citations for each of the sample of 15 universities. UCL and MIT experience the biggest changes in total citation counts: decreases of 34% and 38% respectively (left side chart in Figure 11), when shifted from WoS to MSA. While the remaining universities’ total citation counts seem to have changed at a lesser degree across sources, the differing coverage of DOIs (i.e., different number of DOIs recorded) by each source can still significantly change the average citation counts. This is evidenced in the second chart of Figure 11. Only 4 universities’ rankings remain unchanged across sources (top 3 and last place). All other universities’ positions have shifted at least once across the three sources, with the biggest changes affecting IISC, USP and UNAM.

**Figure 11:**
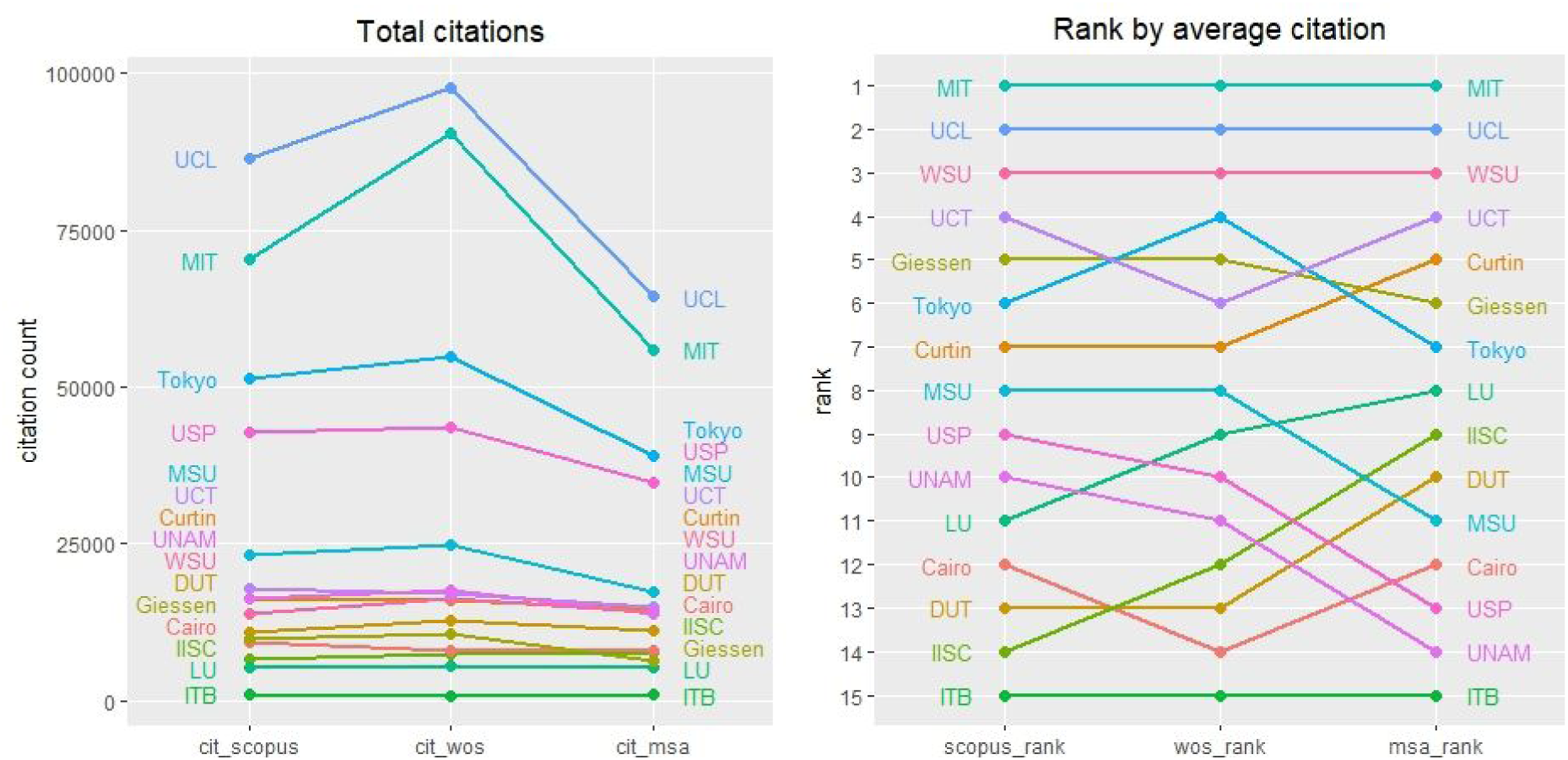
2016 Total citations and rankings in average citations for 15 institutions, as per bibliographic source^46^.

For a more general view, we now include the ranking results for the large set of 155 universities in Figure 12. The results related to universities that have shifted by at least 20 positions across the three sources are highlighted in colour, with universities from English speaking regions in red and non-English ones in orange. This includes 45 universities, 27 in red and 18 in orange. That means almost one-third of the universities have shifted 20 or more positions. The most extreme cases include: Charles Sturt University dropping 146 when moved from WoS to Scopus; Universitat Siegen and University of Marrakech Cadi Ayyad dropping 143 and 112 positions when moved from WoS to MSA.

**Figure 12:**
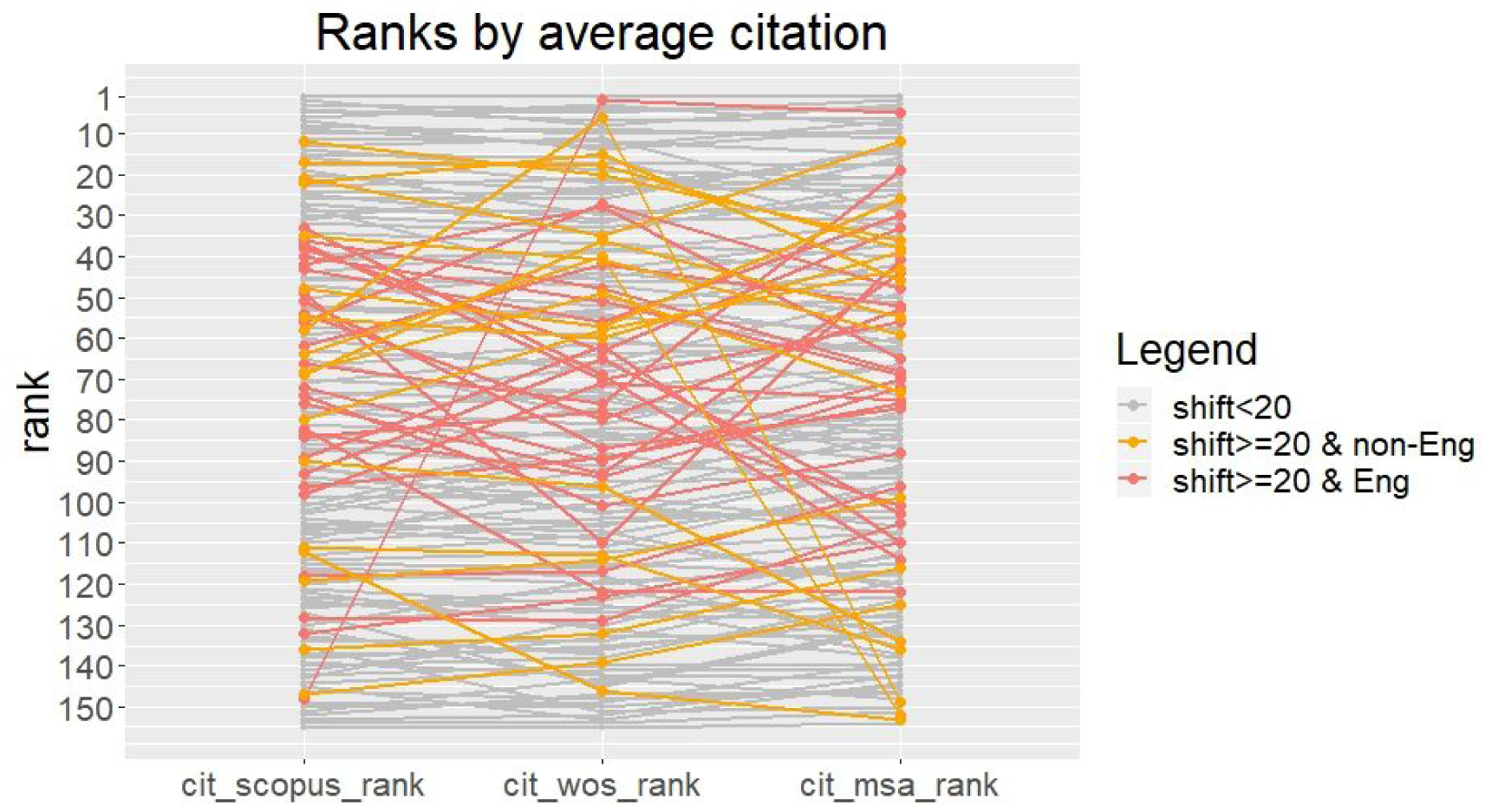
2016 ranking by average citations for 155 universities, as per bibliographic source (with those shifting at least 20 positions displayed in colour)

For further insight into the distribution of shifts across sources, we summarise the pairwise changes to average citations and rankings by average citations into box plots in Figure 13. The median change to average citations when moving from WoS to Scopus is just below zero, while the corresponding medians for WoS to MSA and Scopus to MSA are both just above zero. The corresponding mean values are −0.2, 1.2 and 1.3, respectively. As for the changes to rankings, the median and mean values are all close to zero. The distributions of these box plots are characterised by a concentrated centre with long tails. Again, signifying the existence of two contrasting groups: those universities that were less affected by shifts in bibliographic sources, and those that can have their performance levels, in terms of average citations, greatly altered depending on the choice of source.

**Figure 13:**
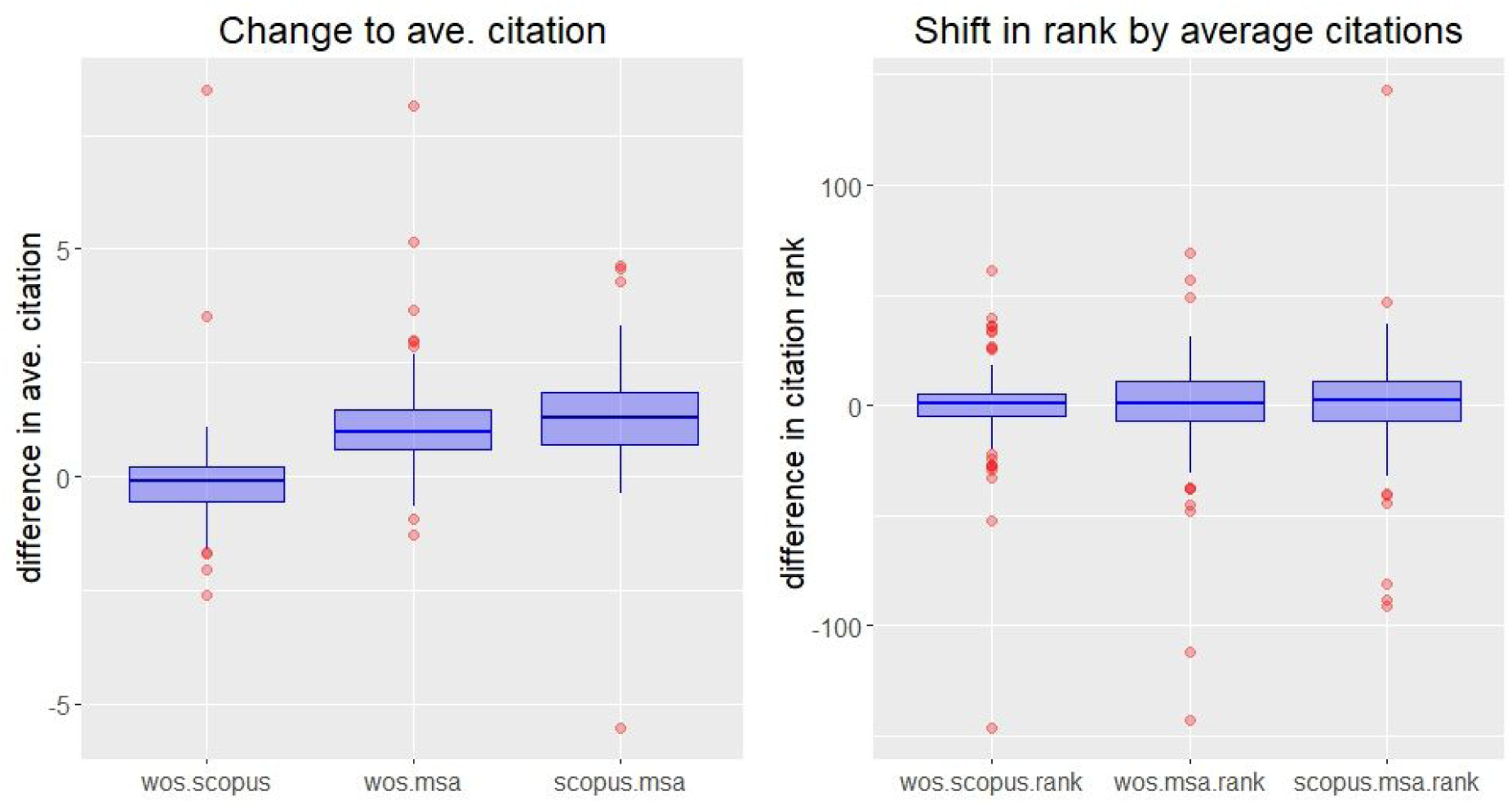
Changes to 2016 average citations (left) and rank by average citations (right) when moving from one source to another for 156 universities.

### 4.5 OA status

A recent topic of interest is the amount of OA publications produced at different levels of the academic system. In particular, universities may wish to evaluate their OA standings for compliance with funder policies and OA initiatives. For objects with DOIs (and, in particular, Crossref DOIs), various information on accessibility can be queried through Unpaywall^47^. We match all DOIs from the sample of 15 universities to the Unpaywall metadata and calculated the percentage of OA output across each bibliographic source and for all (unique) DOIs combined. This is presented in Table 10.

**Table 10:** Total level of OA for all DOIs in our sample of 15 universities, from 2000 to 2018, as per bibliographic source.

|  | WoS | Scopus | MSA | Combined |
| --- | --- | --- | --- | --- |
| **Number of DOIs<sup>48</sup>** | 735832 | 734515 | 907239 | 1202032 |
| **OA count** | 317021 | 294655 | 367100 | 498929 |
| **%OA** | 43.1 | 40.1 | 40.5 | 41.5 |

There does not appear to be substantial changes to the overall OA percentage when shifting across sources for the combined sets of DOIs. However, we should keep in mind that there are significant differences in each source’s DOI coverage, as observed earlier.

To see whether such consistency in OA percentages carries over to the institutional level for 2016, we again filter the data down to each university. Figure 14 provides the percentages of OA output and the corresponding relative ranks for each institution, as per set of 2016 DOIs indexed by each source and also recorded in Unpaywall. It is observed that, for quite a few universities, the OA percentages considerably vary depending on which source is used to obtain the sets of DOIs. The most extreme case is again ITB, which had about a 20% drop when moving from WoS to Scopus. Also, the direction of OA percentage changes differ across universities. For example, OA percentage for MIT decreased when moving from Scopus to MSA, but the opposite occurred for USP. This is especially critical if one is to compare the relative OA status across universities, which can vary according to source of DOIs used. As for OA ranks, it seems to indicate a group of universities not affected by changing source, while the other group have their ranks shifted significantly. The most affected cases seem to be USP, ITB and UNAM.

**Figure 14:**
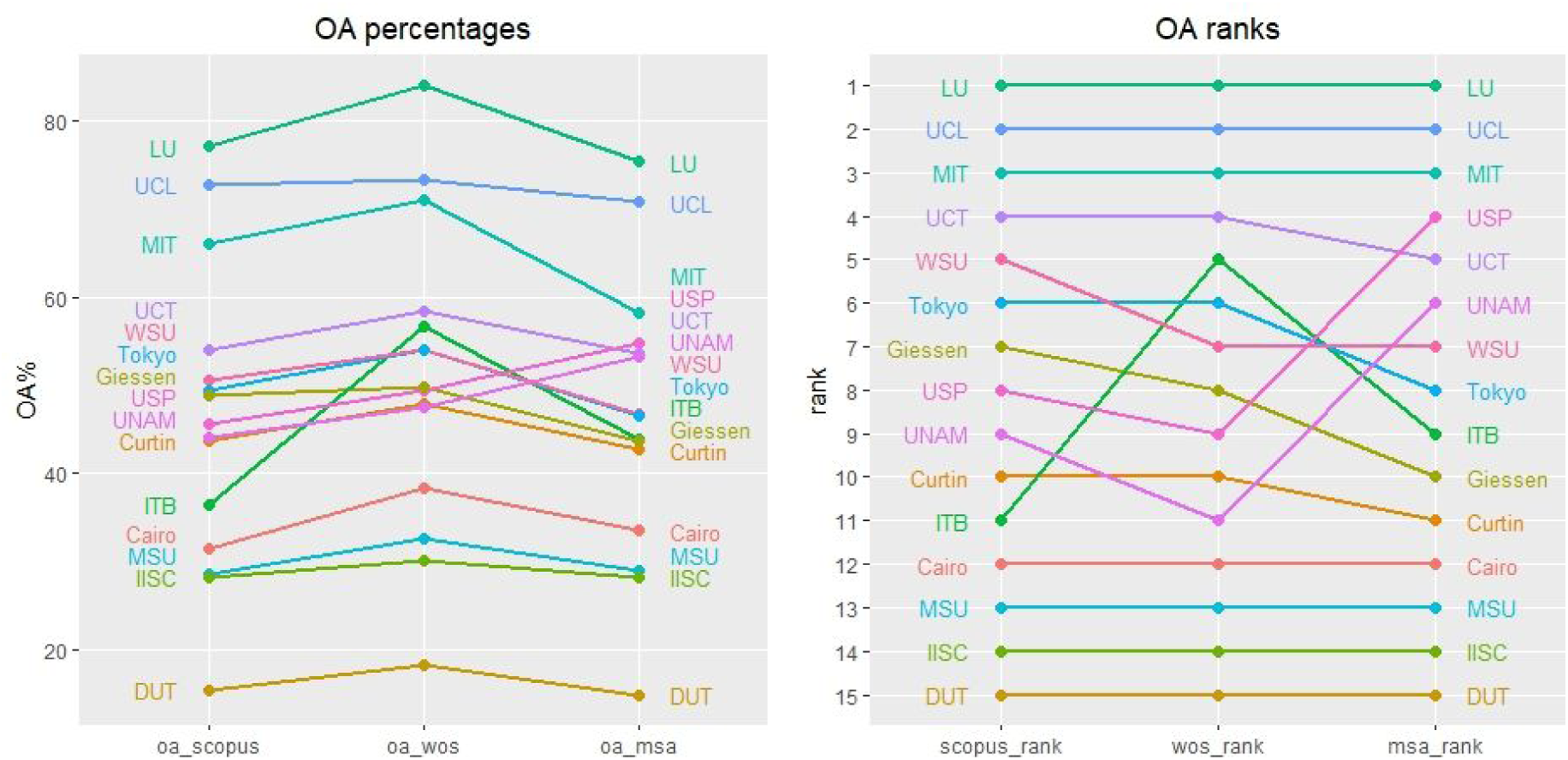
2016 Total OA percentages (left) and OA rankings (right) for 15 institutions, as per bibliographic source^49^.

The effects on OA levels and ranks are more difficult to express directly for the larger set of 155 universities. Again, instead of labelling the full set of universities, we highlight only those that have shifted by 20 positions or more at least once. This is displayed in Figure 15. There are 24 out of 155 universities that have shifted at least 20 positions in OA ranking when moved across sources. Seventeen of these are from non-English speaking regions, including six Latin American universities (out of seven in the full set). This is an indication of the potential difference in coverage of the three sources due to language.

**Figure 15:**
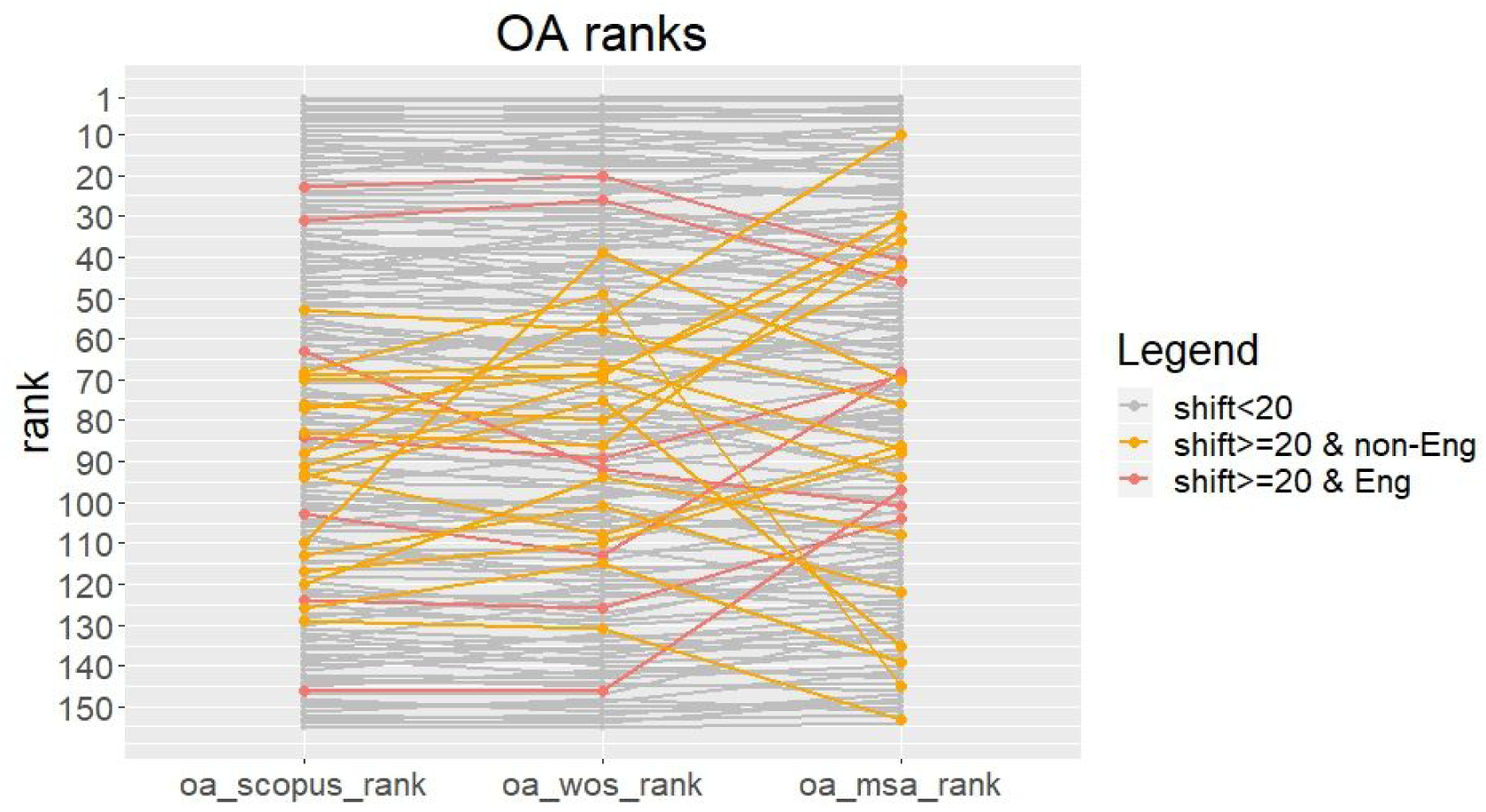
2016 OA rankings for 155 universities, as per bibliographic source (with those shifting at least 20 positions displayed in colour)

Analogous to the earlier analysis on citations, we calculate differences in OA percentages and OA ranks when shifting from one source to another and present these in a number of box plots in Figure 16.

**Figure 16:**
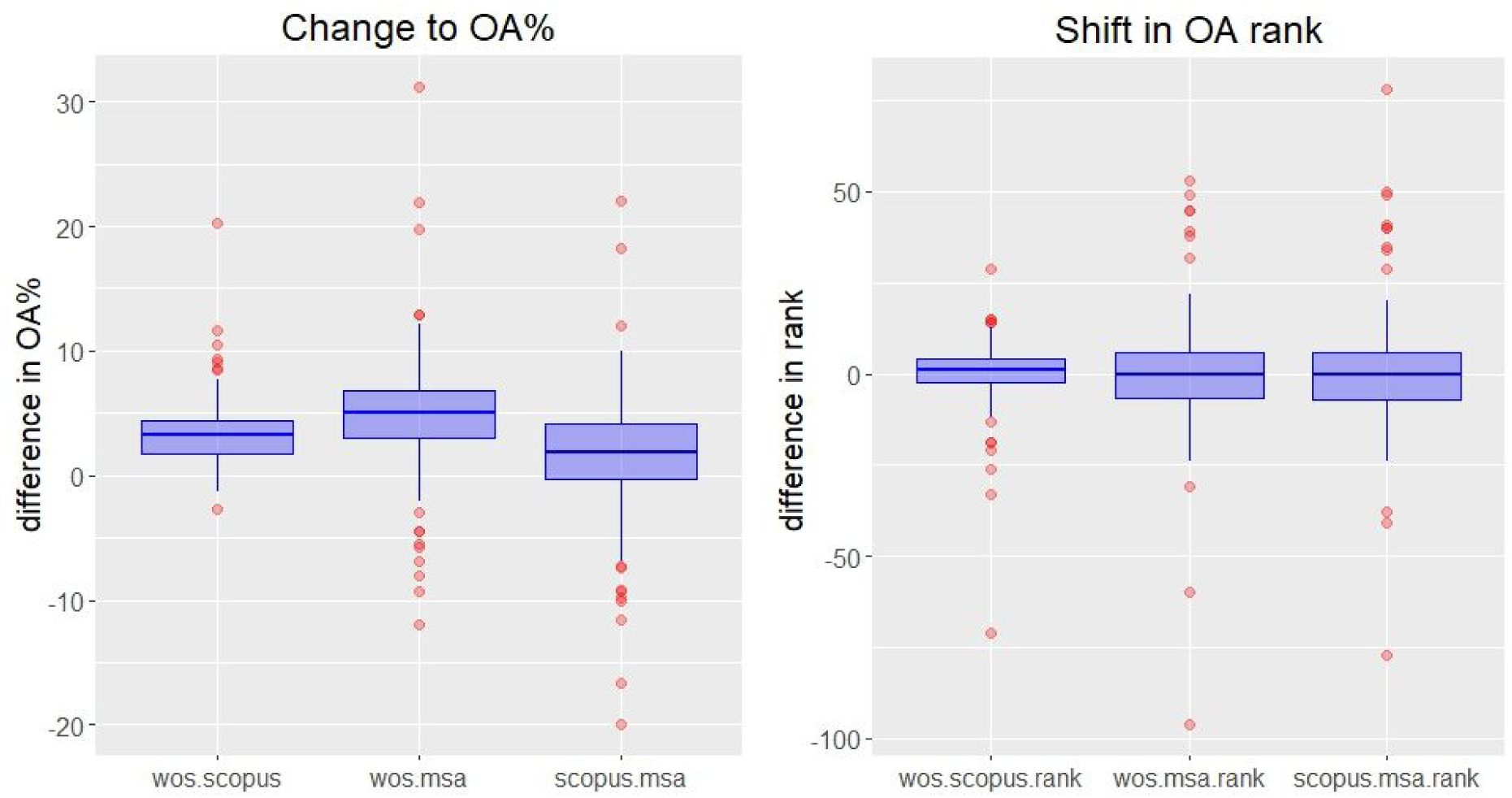
Changes to 2016 OA% (left) and OA rank (right) when moving from one source to another for 155 universities.

Evidently, the median OA% changes when shifting from WoS to Scopus, WoS to MSA, and Scopus to MSA are all positive. The corresponding mean changes are also positive at 3.4%, 4.9% and 1.5% respectively. The median and mean changes to rankings are all close to zero. However, in both OA% and OA rank changes, there are many recorded extreme points (including both negative ones and positive ones). These include an OA% change as large as 31.1% (moving from WoS to MSA) and an extreme drop in OA rank of 96 positions (MSA to WoS). The general distributions of both changes to OA% and changes to OA rankings are characterised by high central peaks and long tails. This implies that, while changes are small for a bulk of the universities, there is also a significant number of cases where universities are largely affected by shifts in data sources.

### 4.6 Manual cross-validation

This section provides a summary of our manual cross-validation results of DOIs exclusively indexed by each source. For each of the 15 institutions, we randomly sampled 40, 30 and 30 DOIs from their sets of 2016 DOIs exclusively indexed by WoS, Scopus and MSA, respectively (i.e., sections W, S and M from Venn diagram in Figure 3). This was done after the removal of DOIs that match-up to other sources in a different year (this includes the neighbouring two years, i.e., 2014, 2015, 2017 and 2018). Subsequently, these lists of DOIs go through a thorough manual cross-validation process. Various questions were asked against each DOI and compared across the three bibliographic sources. These are summarised into a table in Appendix 9.

In the following, we shall highlight some of the main findings in a few simple charts, with further detailed analysis provided in Appendix 9. Firstly, we focus on the plausibility of affiliation associated with each DOI.

In Figure 17, we present results related to affiliation of each DOI as per source. For each DOI, the target affiliation is checked against its online original document^50^. When the original document is not accessible (e.g., not OA), the affiliation is matched against the other two sources. The decision is made to indicate the affiliation as *plausible* when the target affiliation (i.e., affiliation as per our data collection process) appears exact (including obvious versions of the university name) on the document, a plausible affiliation name variant^51^ appears on the document, or the affiliation is confirmed by at least one of the other two bibliographic sources (if found). This should (roughly) inform us about whether each source have correctly assigned these DOIs to the target affiliations.

**Figure 17:**
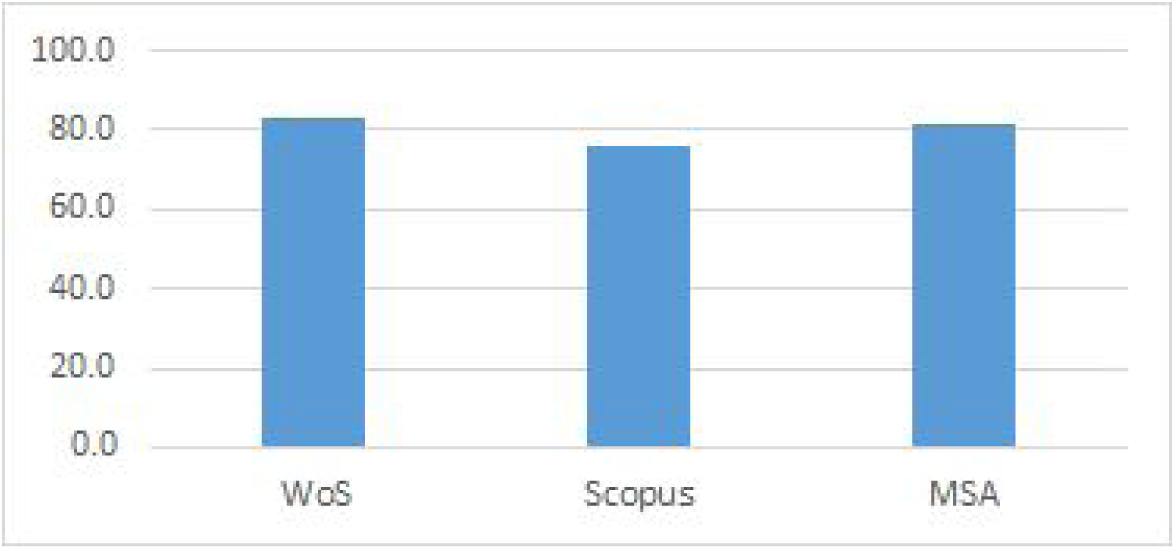
Percentage of DOIs with plausible affiliation as per matching against original document or the other two sources.

**Figure 18:**
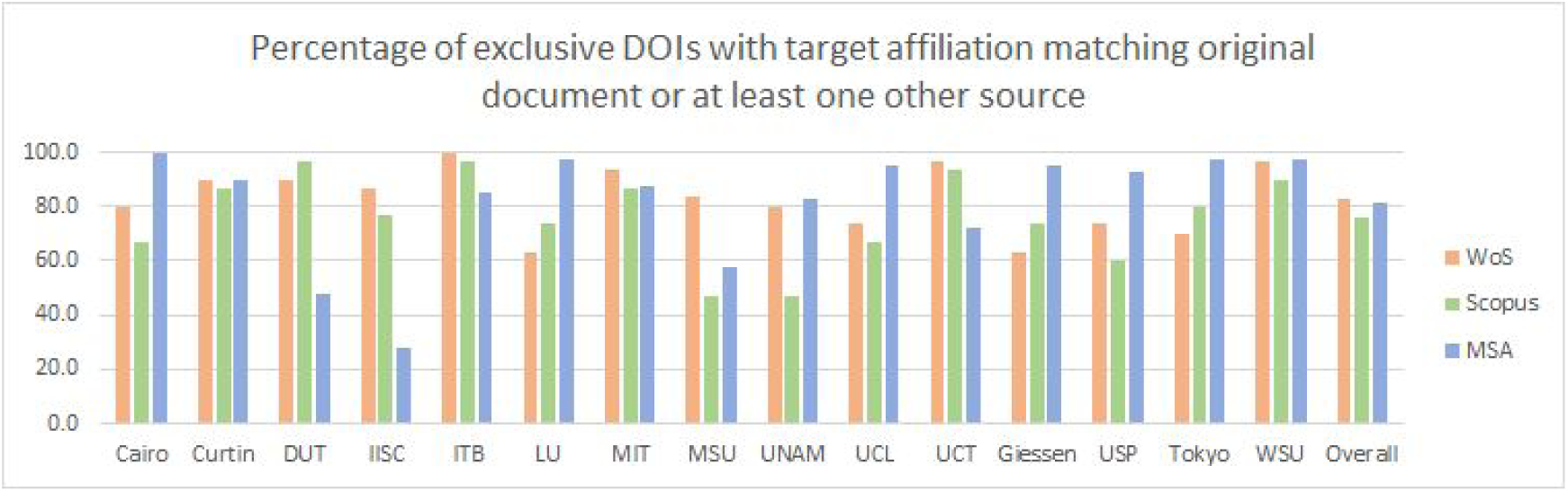
Percentage of exclusive DOIs from one source that has a plausible match to target affiliation as per original document or at least one other source.

The result shows that all sources have only correctly assigned roughly 80% of their respective DOIs from our sample to the target affiliations, with very little difference in performance across the sources. When this filtered down as per university, we see a more varied performance across universities.

Interestingly, not all percentages are very high across the universities. This is especially apparent for DUT and IISC, where MSA seems to have affiliated many DOIs to these two institutions without the target affiliations actually appearing on the original documents or confirmed by another source. Similarly, for DOIs that were assigned to MSU and UNAM by Scopus, only 46.7% (for both institutions) have a plausible affiliation match.

We have also checked each DOI against the DOI string actually recorded as per original document (where applicable) or via doi.org. These percentages (of correct DOIs) are 93.1%, 98.2% and 96.7 for WoS, Scopus and MSA respectively (with all 15 institutions combined). While these numbers are relatively high, the significant number of errors suggests that DOIs are not being systematically checked against authoritative sources such as Crossref which we find surprising. In addition the nature of these errors which in some cases appear to be transcription or OCR errors is concerning (see supplementary Information in Appendix 10).

We now take an overview of results from the DOI and title matching, given in Figure 19. As an initial analysis, no affiliation information is considered here and the results represent all DOIs for the 15 universities combined. Each bar represent the percentage of output corresponding to DOIs (that initially appear to be) exclusively indexed by one source that can be found in another source by DOI matching and title matching (via manual searches online). For example, the first bar corresponds to objects with DOIs sampled from Scopus. The height of the blue bar shows the percentage of these object that can be found in WoS by DOI matching. The orange bar then indicates how much more can be found by title matching.

**Figure 19:**
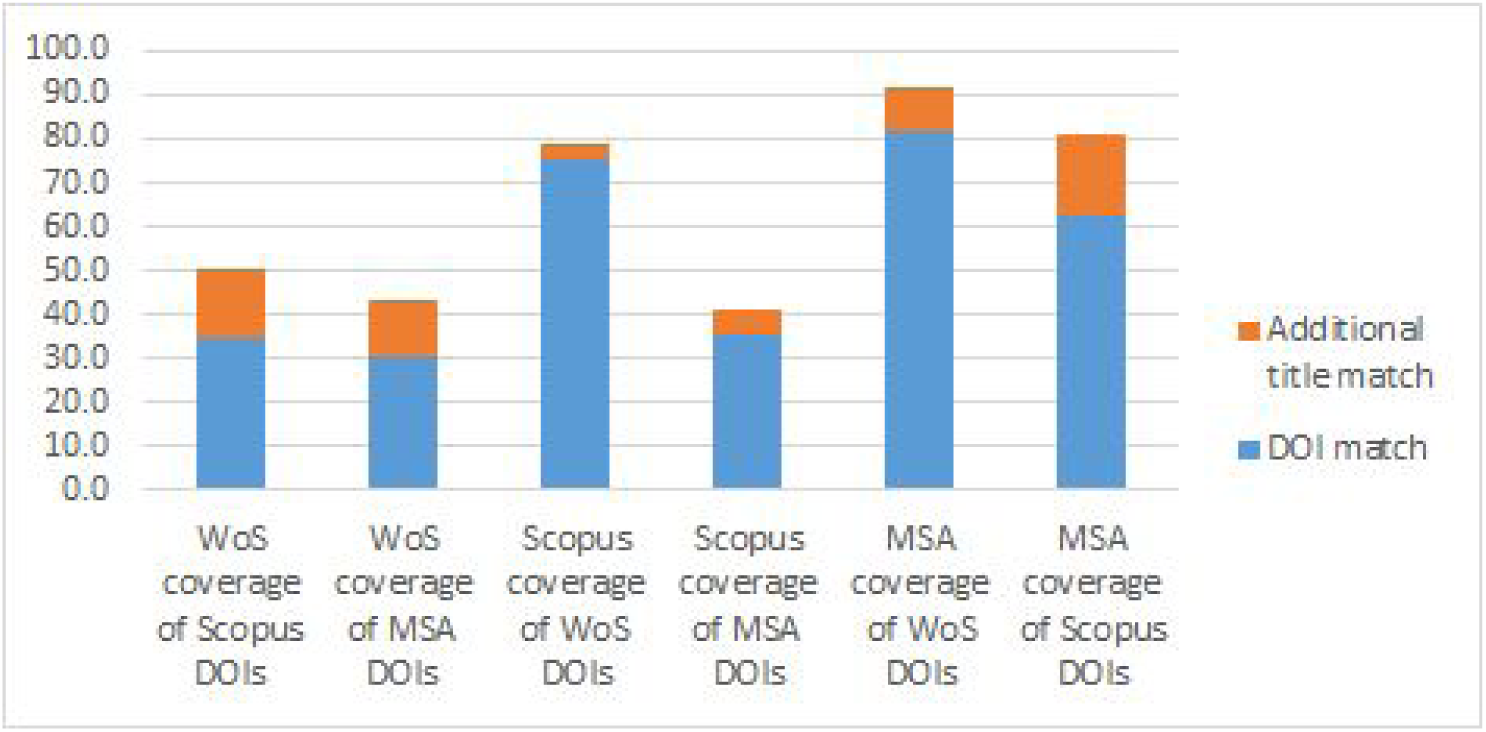
Percentage of DOIs found in another source by DOI and title matching.

We found that in all cases where there is a DOI match, there is also a title match. However, the opposite is not necessary true. Hence, title matching increases the coverage slightly in all scenarios. This does imply that all three sources have missing DOIs in their metadata, though there appears to be fewer cases for Scopus. Scopus also seems to have a good coverage of DOIs from WoS. More strikingly, very high proportion of DOIs and titles from WoS and Scopus are found in MSA. In contrast, much fewer MSA DOIs and titles are covered by WoS and Scopus.

In Figure 20, we added affiliation matching to the mix, i.e., check whether the target affiliation (i.e., affiliation as per our data collection process) appears in the metadata of the matching source, after an object is found by DOI or title match. This decreased the coverage in all cases, indicating the potential disagreement of affiliation across sources. MSA is the most affected out of the three sources.

**Figure 20:**
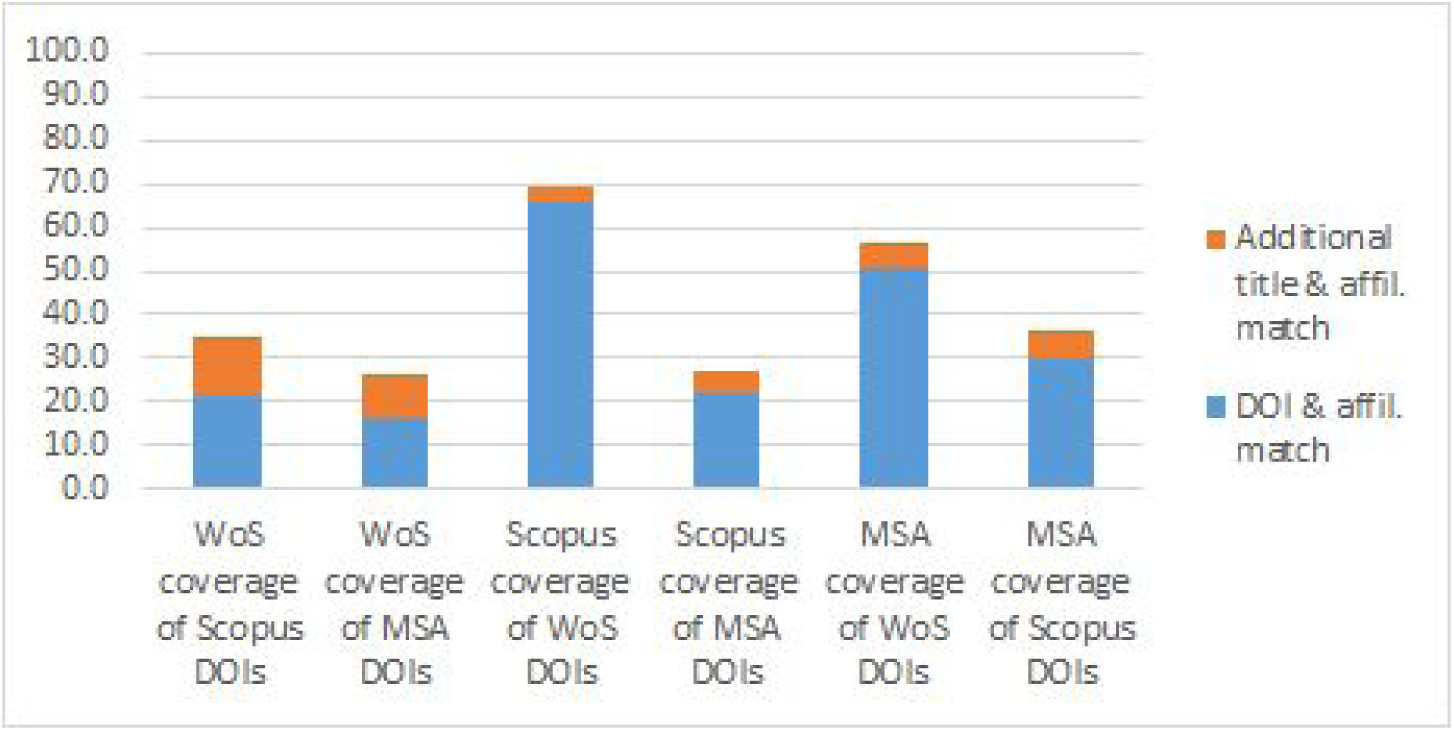
Percentage of DOIs found in another source by DOI and title matching, combined with affiliation matching.

The general picture that has emerged is that MSA seems to have good coverage of DOIs that initially appeared to be exclusively from WoS or Scopus. However, it falls short on getting the affiliation correct and recording the corresponding DOIs. MSA also seems to bring in more objects that genuinely appear to be exclusive to MSA. The correctness of affiliation metadata for these is high overall but tends to vary across institutions.

## 5. Limitations and challenges

One obvious limitation is our focus on DOIs and our dependence on the uniqueness of DOIs. We do note that there may be research objects with multiple DOIs and related objects may also be assigned a common DOI (e.g., books can fit in both cases). A related matter is the correctness of DOIs, i.e., whether they were recorded correctly (as per doi.org) in each source’s metadata. DOIs that did not generate Unpaywall returns could include such cases. While our manual cross-validation process did check our samples against doi.org, it is not clear what the scale of this issue is for the overall data.

Our manual cross-validation process is carried out over a number of months after the initial data collection process. This means that there may be potential discrepancies between metadata content at the time of collection and time of manual search. However, we expect such cases to be few given that we are focused on 2016 data and a number of manual spot checks did not reveal any obvious such cases.

Both API and manual searches for WoS and Scopus may be limited to the subscription model of the authors’ home institution at the time of access. On the other hand, matching identifiers have also proved to be challenging. For example, a few institutions have multiple Scopus IDs (e.g., multiple campuses), without an overarching ID. For the three cases we have encountered, out of the 155 universitieswe have selected what appeared to be the main campus IDs. Other challenges include: limit of Unpaywall and OpenCitations coverages to Crossref DOIs, manual cross-validation limited to DOI and title searches only, subjectivity in linking plausible affiliation names.

## 6. Conclusion

This article has taken on the task of comparing and cross-validating various bibliographic characteristics (including coverage, publication date, OA status, document type, citations and affiliation) across three major research output indexing databases, i.e., WoS, Scopus and MSA. This is done mainly with a focus on identifying institutional level differences and the corresponding effects of using different data sources in comparing institutions. Our data consists of all objects with DOIs extracted from the three bibliographic sources for an initial sample of 15 universities and a further supplementary 140 universities (used only where applicable).

Firstly, we found the coverage of DOIs not only differ across the three sources, their relative coverages are also non-symmetrical and the distribution of DOIs across the sources varied from institution to institution. This means that the sole use of one bibliographic source can potentially seriously disadvantage some institutions and advantage others in terms of total number of output. While the general level of agreement on publication year is high across sources, there were individual universities with large differences in coverage per year. The comparison of document types showed that different sources can systematically add coverage of selected research output types. This may be of importance when considering the coverage of different research discipline areas.

Our subsequent analyses further showed that while the aggregate levels (i.e., for 15 universities combined) in citation counts and OA levels varied little across sources, there are significant impacts at the institutional level. There were clear examples of universities shifting dramatically in both of these metrics when moving across sources, some in opposite directions. This makes any rank comparison of citations or OA levels strongly dependent on the selection of bibliographic source.

Finally, we implemented a manual cross-validation process to check metadata records for samples of DOIs that initially appeared to be exclusive from each source, for each of the 15 universities. The records were compared across the three bibliographic sources and against (where accessible) the corresponding online research documents. The process revealed cases of missing links between metadata and search functionalities within each database (for both affiliation and DOI). This means the real coverage of each source is unnecessarily truncated. Overall, it appears that MSA has the highest coverage of objects that initially appeared exclusive to other sources. However, it often has missing DOIs and affiliations that do not match with WoS, Scopus or online documents.

There is also strong evidence that the effects of shifting sources may be more prominent for non-English speaking and non-European universities. Similar signs were observable for universities that are medium-ranked in both citations and OA levels, while those that achieve high rankings in these measures show much smaller shifts in position when the data source is changed. Universities that are highly ranked on these measures also tend to be highly ranked in general rankings like the THES, suggesting a bias in reliability and therefore curation effort towards prestigious universities.

Our concluding message is: any institutional evaluation framework that is serious about coverage should consider incorporating multiple bibliographic sources. The challenge is in concatenating unstandardised data infrastructures that do not necessarily agree with each other. For example, one primary task would be to standardise the publication dates, especially for longitudinal study. This may be possible, to a certain level, using Crossref or Unpaywall metadata, as an external reference set. Such problems are by no means trivial. However, it has the potential to greatly enhance the delivery of fairer and more robust evaluation.

## Appendix 1 Affiliation IDs and search terms

Appendix 1: Affiliation IDs and search terms
| Institution | GRID ID | WoS search term | Scopus ID | MSA ID |
| --- | --- | --- | --- | --- |
| Cairo University | grid.7776.1 | Cairo University | 60010294 | 145487455 |
| Curtin University | grid.1032.0 | Curtin University | 60031226 | 205640436 |
| Dalian University of Technology | grid.30055.33 | Dalian University of Technology | 60004538 | 27357992 |
| Indian Institute of Science Bangalore | grid.34980.36 | Indian Institute of Science (IISc) - Bangalore | 60014097 | 59270414 |
| Institut Teknologi Bandung | grid.434933.a | Institute Technology of Bandung | 60069382 | 134635517 |
| Loughborough University | grid.6571.5 | Loughborough University | 60000891 | 143804889 |
| Massachusetts Institute of Technology | grid.116068.8 | Massachusetts Institute of Technology (MIT) | 60022195 | 63966007 |
| Moscow State University | grid.14476.30 | LOMONOSOV MOSCOW STATE UNIVERSITY | 60007457 | 19880235 |
| National Autonomous University of Mexico | grid.9486.3 | Universidad Nacional Autonoma de Mexico | 60032442 | 8961855 |
| University College London | grid.83440.3b | University College London | 60022148 | 45129253 |
| University of Cape Town | grid.7836.a | University of Cape Town | 60000356 | 157614274 |
| University of Giessen | grid.8664.c | Justus Liebig University Giessen | 60017134 | 200763008 |
| University of Sao Paulo | grid.11899.38 | Universidade de Sao Paulo | 60008088 | 17974374 |
| University of Tokyo | grid.26999.3d | University of Tokyo | 60025272 | 74801974 |
| Wayne State University | grid.254444.7 | Wayne State University | 60009408 | 185443292 |

GRID IDs for the set of 140 supplementary universities:

grid.1001.0, grid.1002.3, grid.1003.2, grid.1004.5, grid.1005.4, grid.1007.6, grid.1008.9, grid.1009.8, grid.1010.0, grid.1011.1, grid.1012.2, grid.1013.3, grid.1014.4, grid.1017.7, grid.1018.8, grid.1019.9, grid.1020.3, grid.1021.2, grid.1022.1, grid.1023.0, grid.1024.7, grid.1025.6, grid.1026.5, grid.1027.4, grid.1029.a, grid.1031.3, grid.1033.1, grid.1034.6, grid.1037.5, grid.10388.32, grid.1039.b, grid.1040.5, grid.1043.6, grid.1048.d, grid.10698.36, grid.10784.3a, grid.10858.34, grid.11135.37, grid.117476.2, grid.11843.3f, grid.11914.3c, grid.12295.3d, grid.12380.38, grid.14709.3b, grid.148374.d, grid.16463.36, grid.16750.35, grid.168010.e, grid.16821.3c, grid.170205.1, grid.17088.36, grid.177174.3, grid.189504.1, grid.19006.3e, grid.19188.39, grid.20861.3d, grid.21006.35, grid.21107.35, grid.213917.f, grid.21729.3f, grid.21925.3d, grid.252547.3, grid.252890.4, grid.254880.3, grid.25879.31, grid.262743.6, grid.264200.2, grid.266842.c, grid.266886.4, grid.267827.e, grid.27755.32, grid.278276.e, grid.35030.35, grid.35403.31, grid.35541.36, grid.35915.3b, grid.38142.3c, grid.410356.5, grid.410714.7, grid.411301.6, grid.411340.3, grid.411840.8, grid.411958.0, grid.412419.b, grid.412522.2, grid.412813.d, grid.412831.d, grid.413050.3, grid.417984.7, grid.419886.a, grid.42505.36, grid.4280.e, grid.4488.0, grid.4514.4, grid.461025.4, grid.4691.a, grid.4708.b, grid.47100.32, grid.47840.3f, grid.4830.f, grid.49481.30, grid.4991.5, grid.5012.6, grid.5037.1, grid.5110.5, grid.5132.5, grid.5252.0, grid.5333.6, grid.5335.0, grid.5380.e, grid.5475.3, grid.5477.1, grid.5510.1, grid.5596.f, grid.5601.2, grid.5606.5, grid.5836.8, grid.5841.8, grid.5842.b, grid.59025.3b, grid.5963.9, grid.6612.3, grid.67105.35, grid.6852.9, grid.7177.6, grid.7247.6, grid.7345.5, grid.7372.1, grid.7400.3, grid.7427.6, grid.7445.2, grid.7700.0, grid.7737.4, grid.77602.34, grid.8096.7, grid.8217.c, grid.8591.5, grid.89336.37, grid.9601.e, grid.9654.e.

## Appendix 2 WoS databases used in this study

Science Citation Index Expanded (SCI-EXPANDED) --1972-present

Social Sciences Citation Index (SSCI) --1972-present

Arts & Humanities Citation Index (A&HCI) --1975-present

Conference Proceedings Citation Index-Science (CPCI-S) --1990-present

Conference Proceedings Citation Index-Social Science & Humanities (CPCI-SSH) --1990-present

Book Citation Index– Science (BKCI-S) --2005-2012

Book Citation Index– Social Sciences & Humanities (BKCI-SSH) --2005-2012

Emerging Sources Citation Index (ESCI) --2015-present

Current Chemical Reactions (CCR-EXPANDED) --1985-present

(Includes Institut National de la Propriete Industrielle structure data back to 1840) Index Chemicus (IC) --1993-present

## Appendix 3 Venn diagrams of DOIs from 2016 for each institution^52^

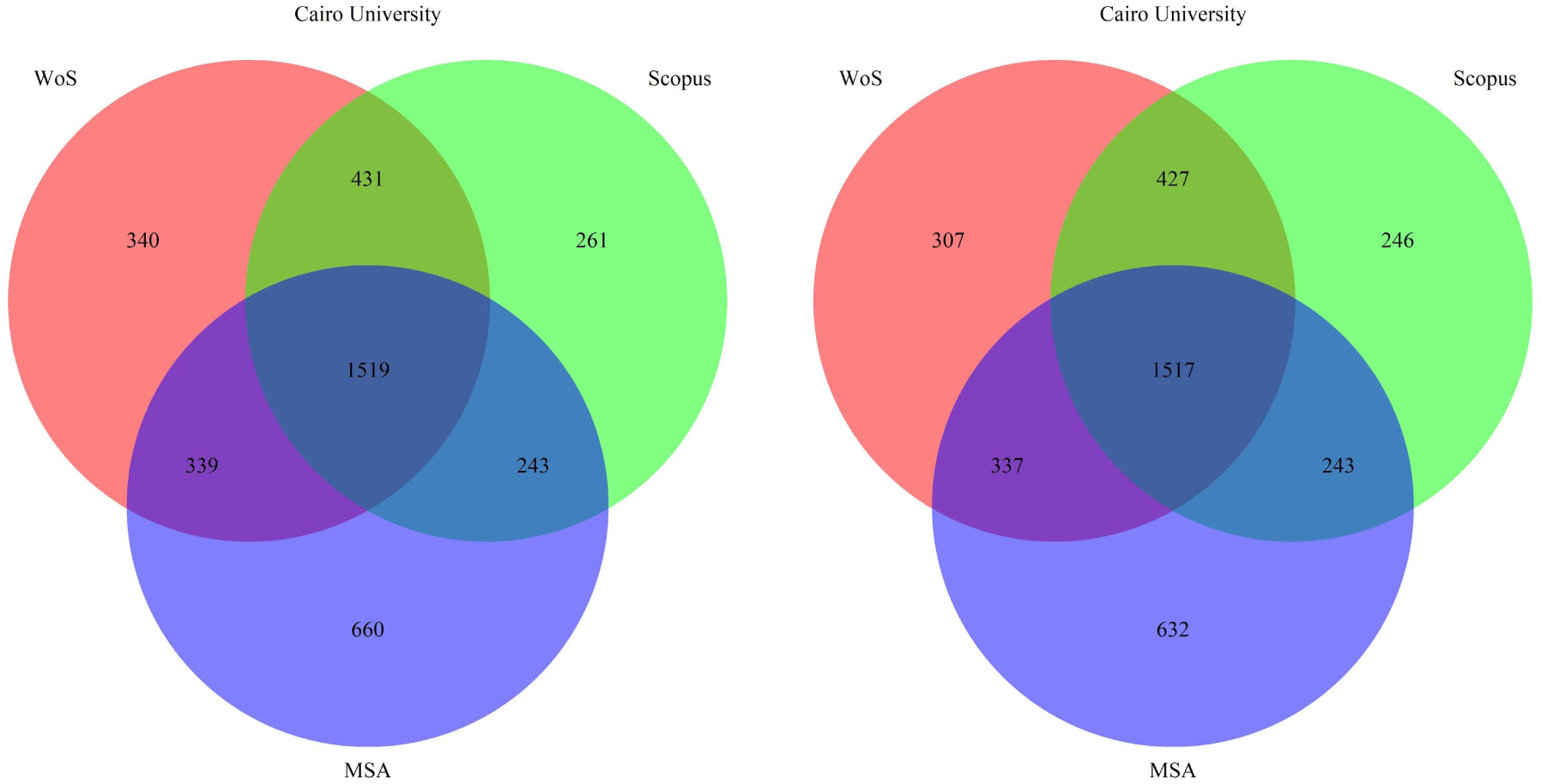
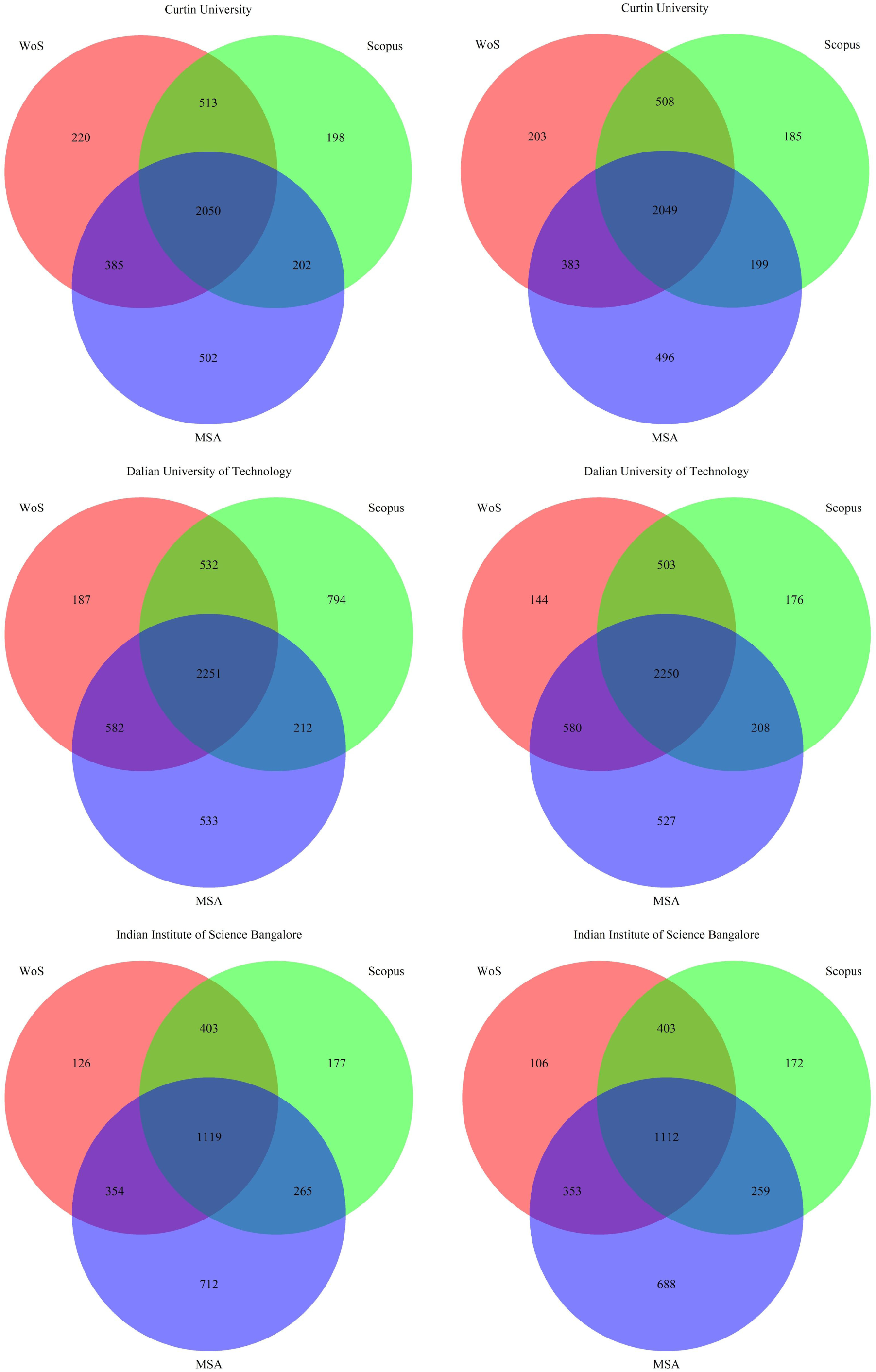
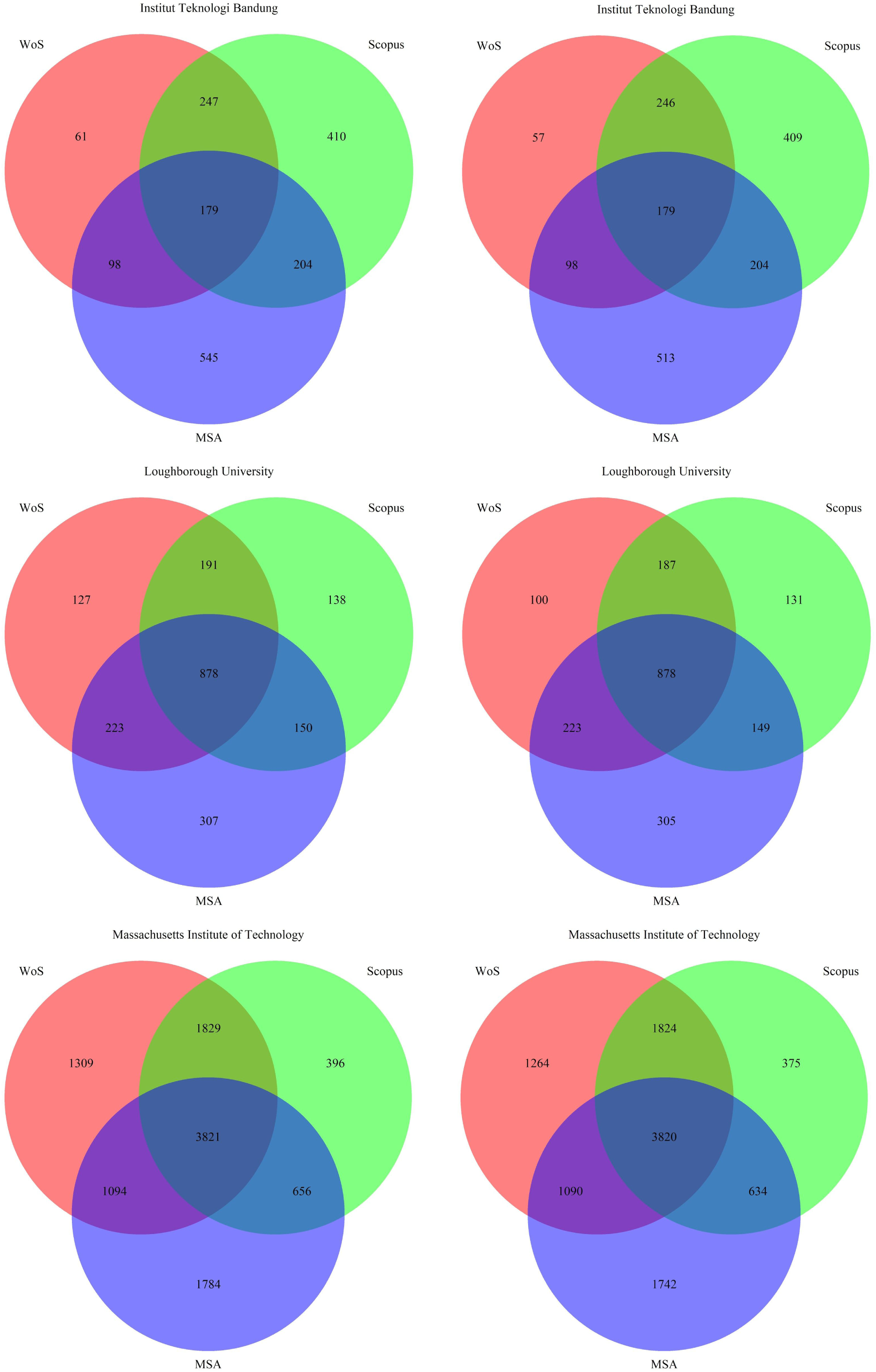
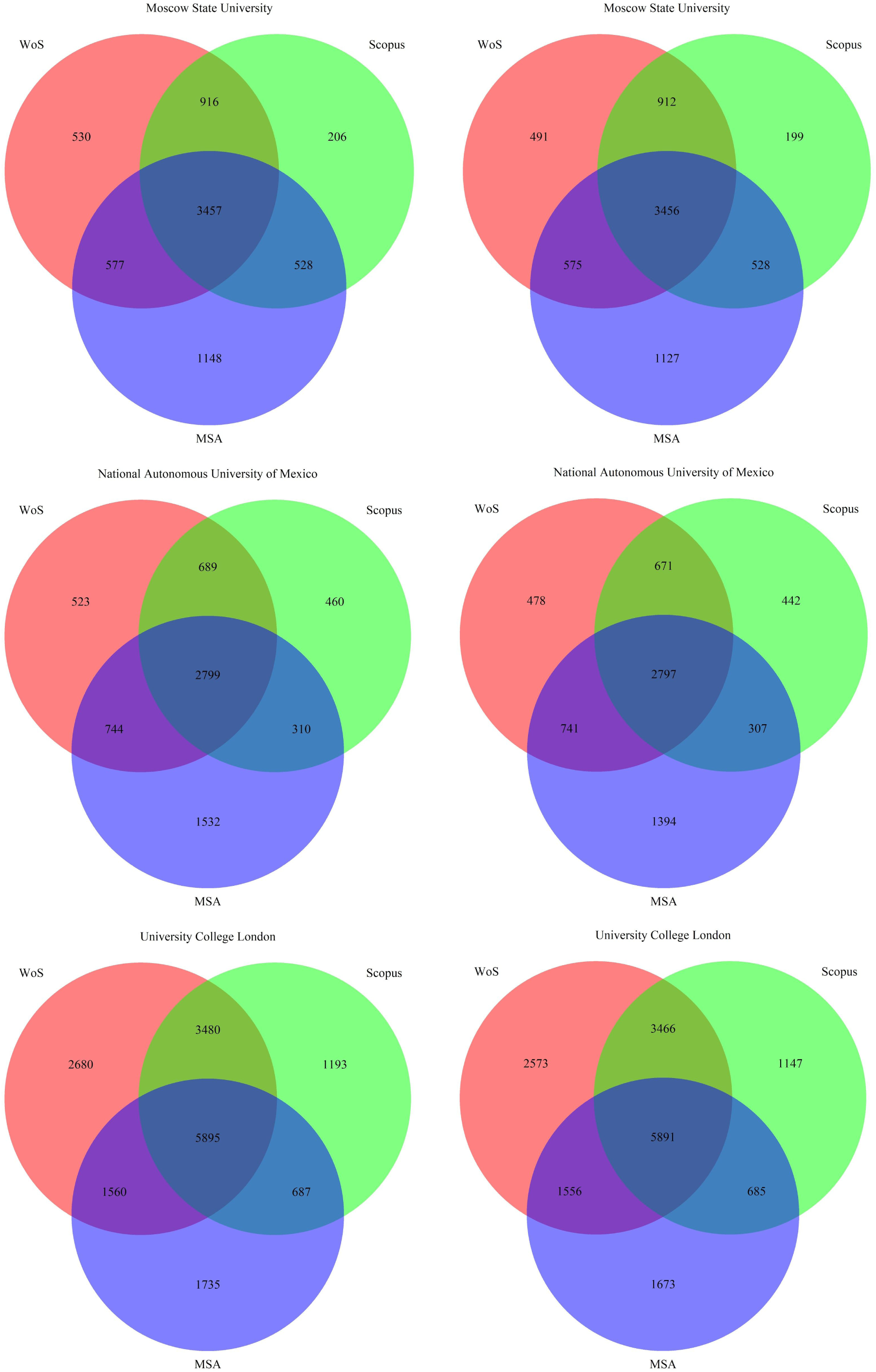
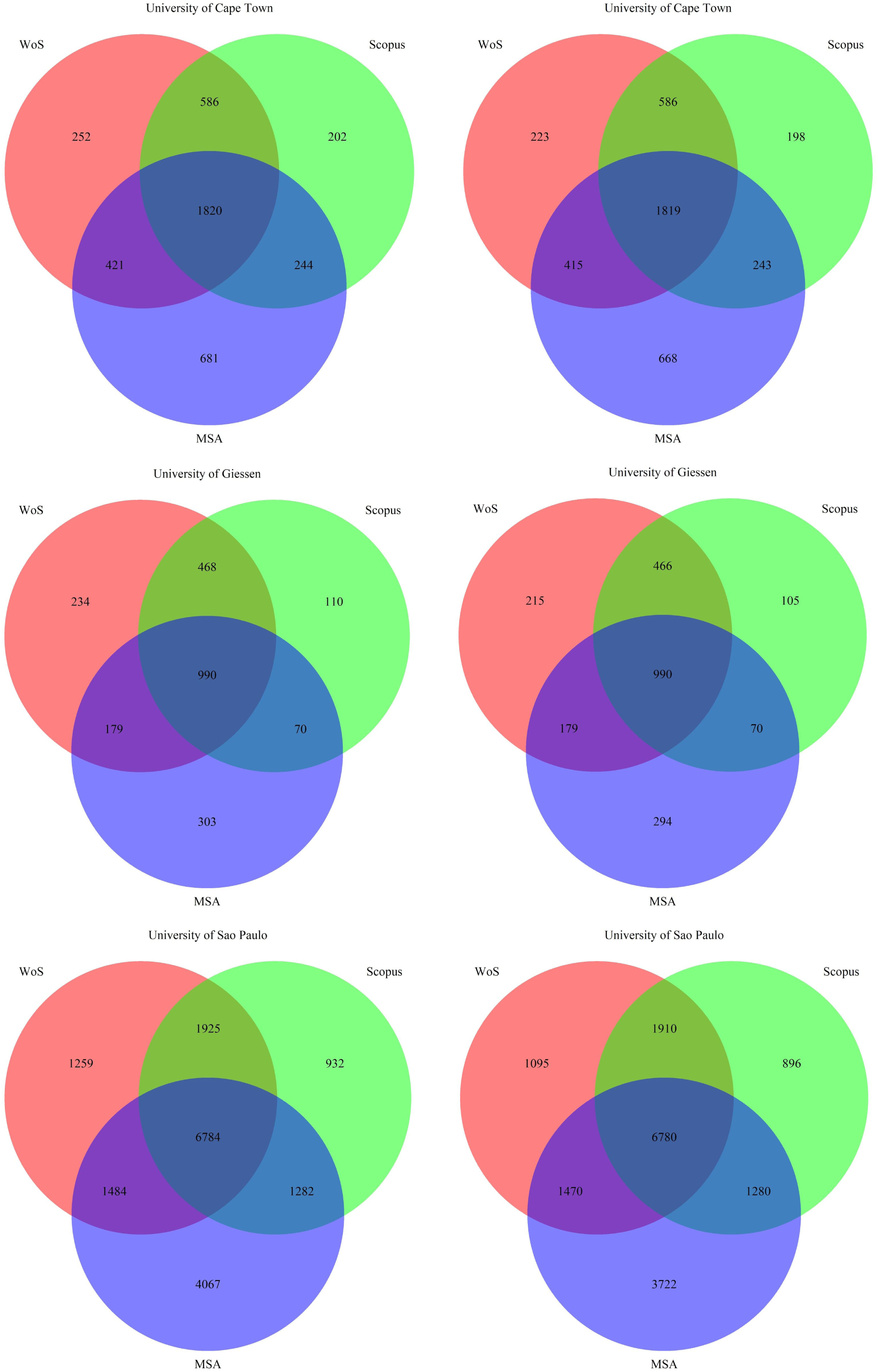
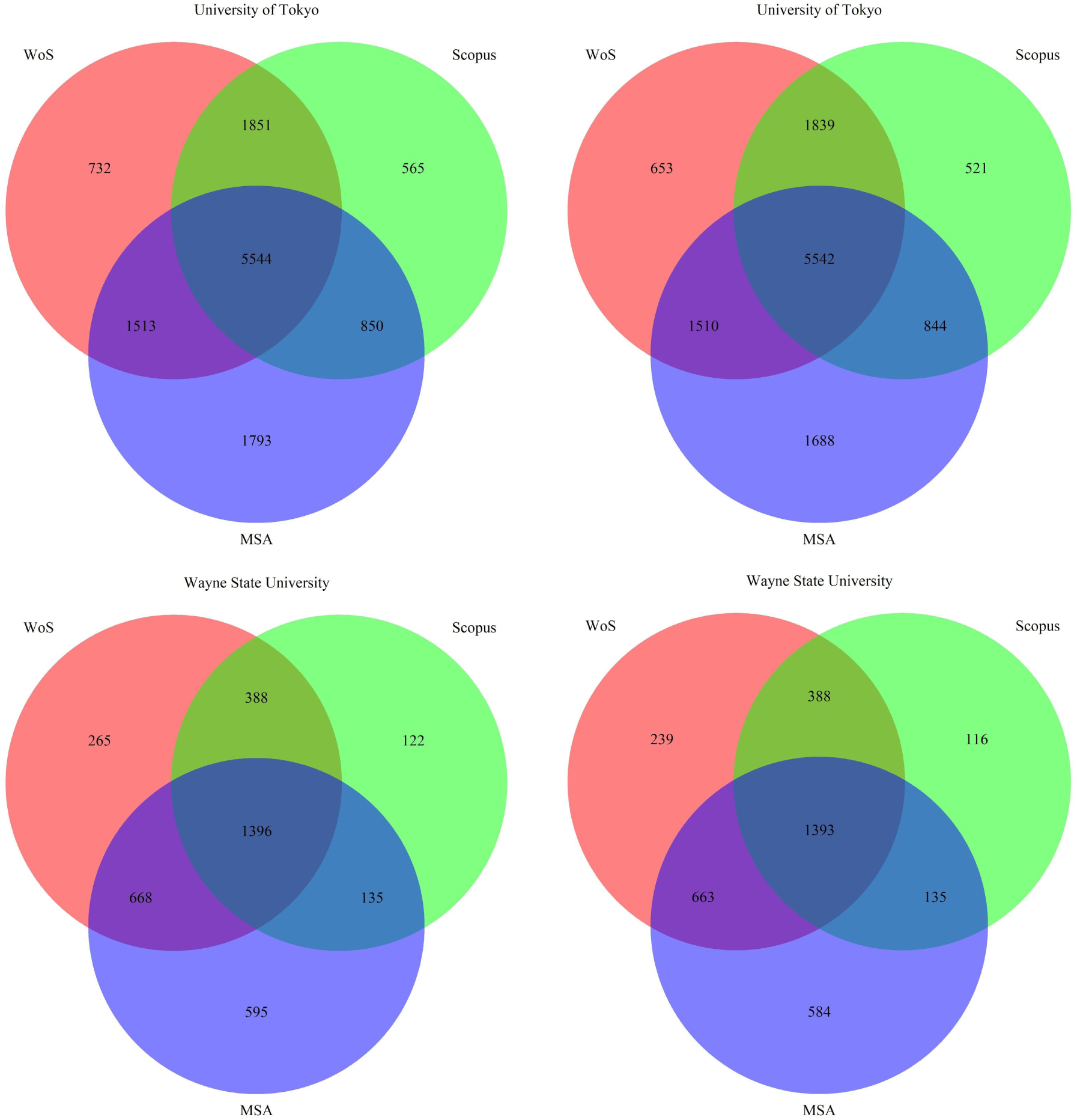

## Appendix 4 Scatterplots of proportions of coverage by WoS, Scopus and MSA^53^

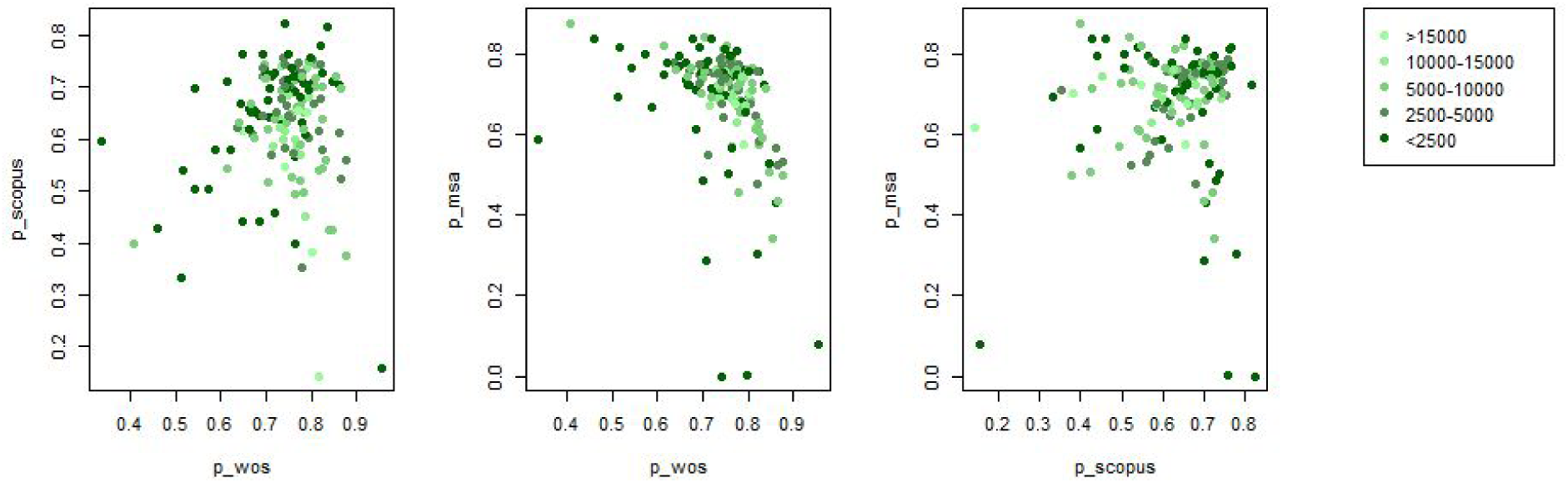

## Appendix 5 Comparison of distributions of *d_1_*, *d_2_* and *d_3_* against Venn diagrams generated from various distributions

**Figure A4.1:**
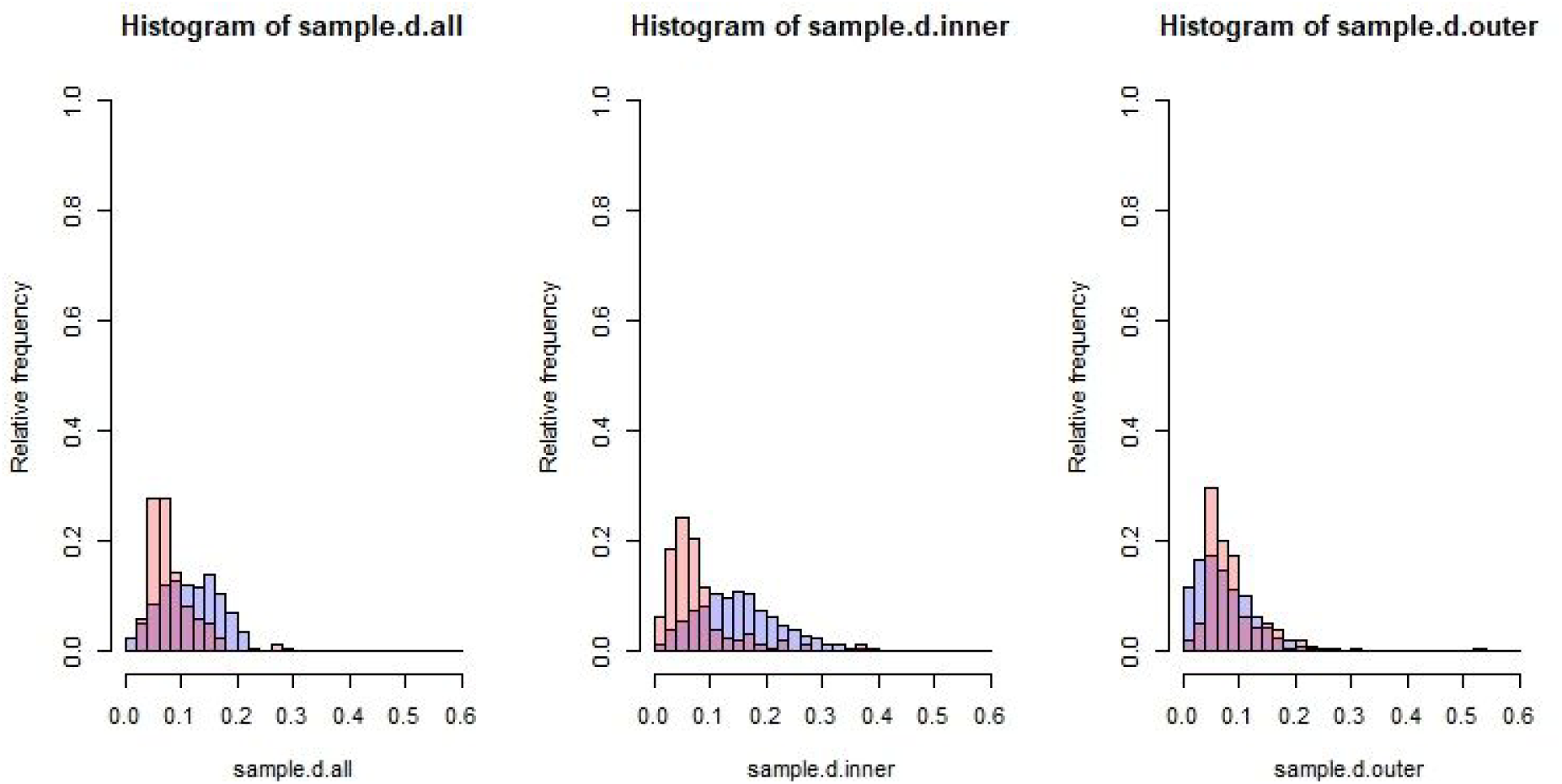
Histograms of *d_1_*, *d_2_* and *d_3_* (left to right, respectively) for our data (in red) and for randomly generated Venn diagrams (in purple) from independent uniform distributions.

**Figure A4.2:**
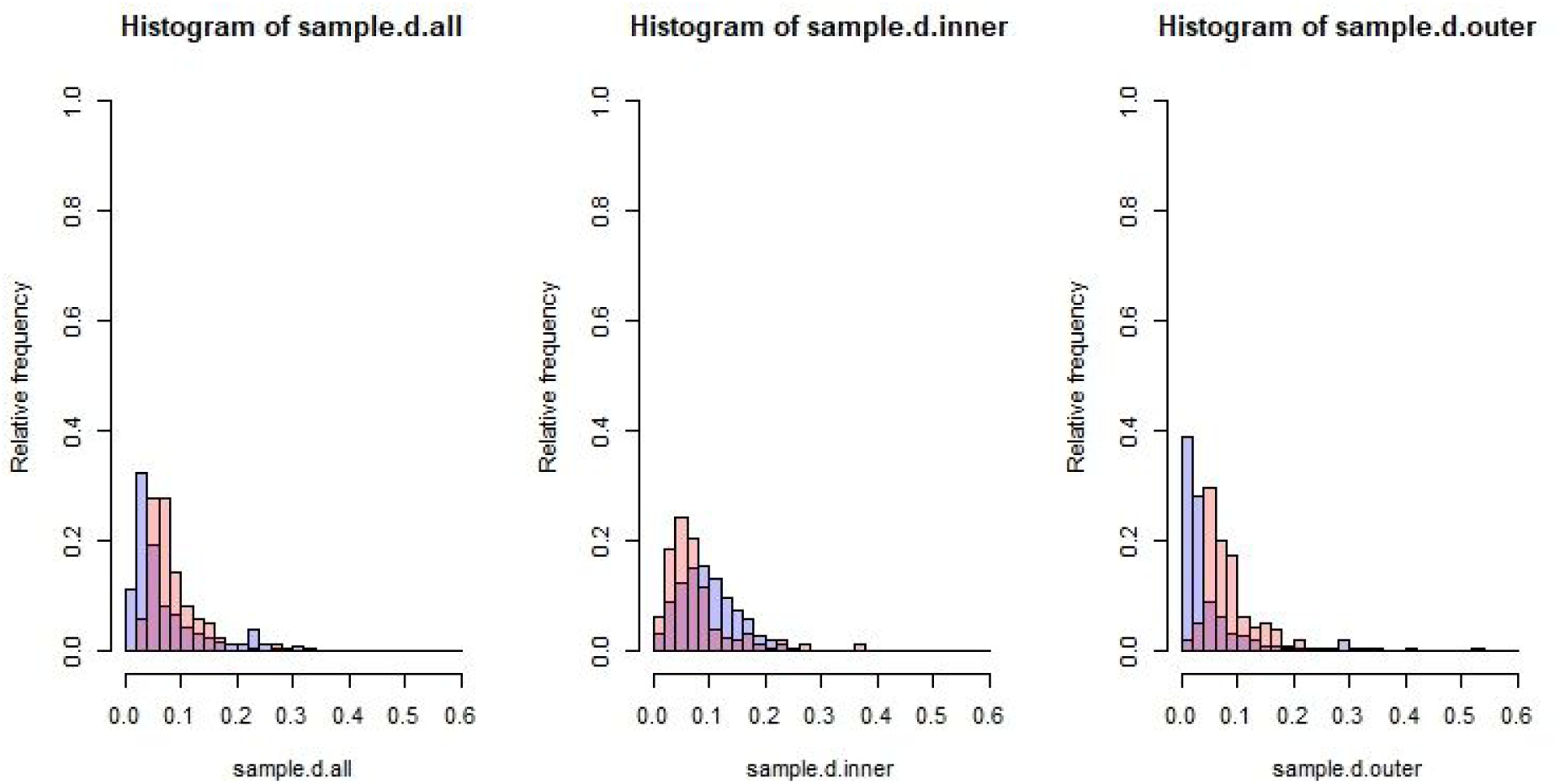
Histograms of *d_1_*, *d_2_* and *d_3_* (left to right, respectively) for our data (in red) and for randomly generated Venn diagrams (in purple) from multivariate skew Cauchy distribution^54^.

**Figure A4.3.**
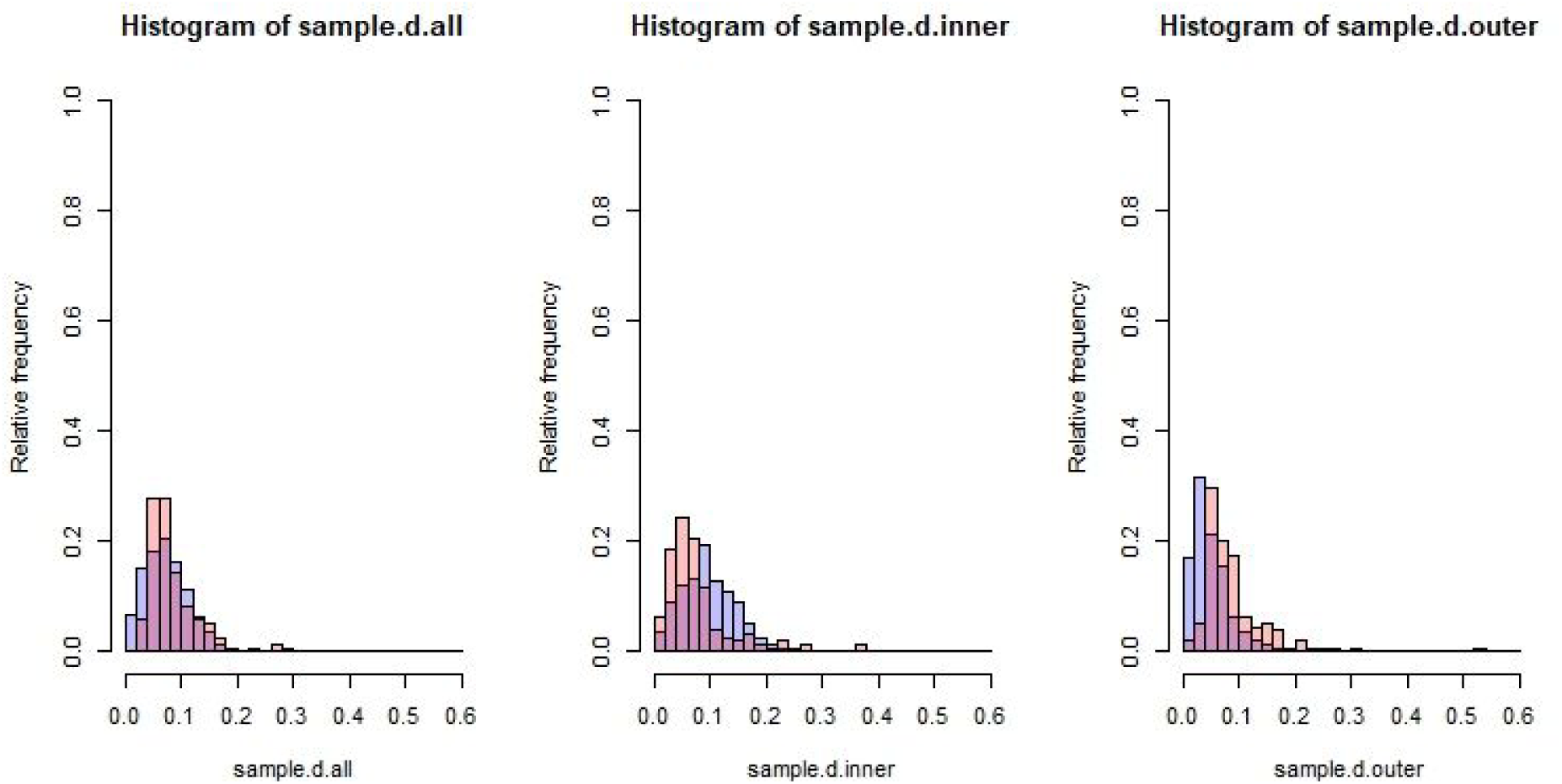
: Histograms of *d_1_*, *d_2_* and *d_3_* (left to right, respectively) for our data (in red) and for randomly generated Venn diagrams (in purple) from multivariate skew normal distribution^55^.

## Appendix 6 Hierarchical cluster analysis of institutions by Venn diagram proportions

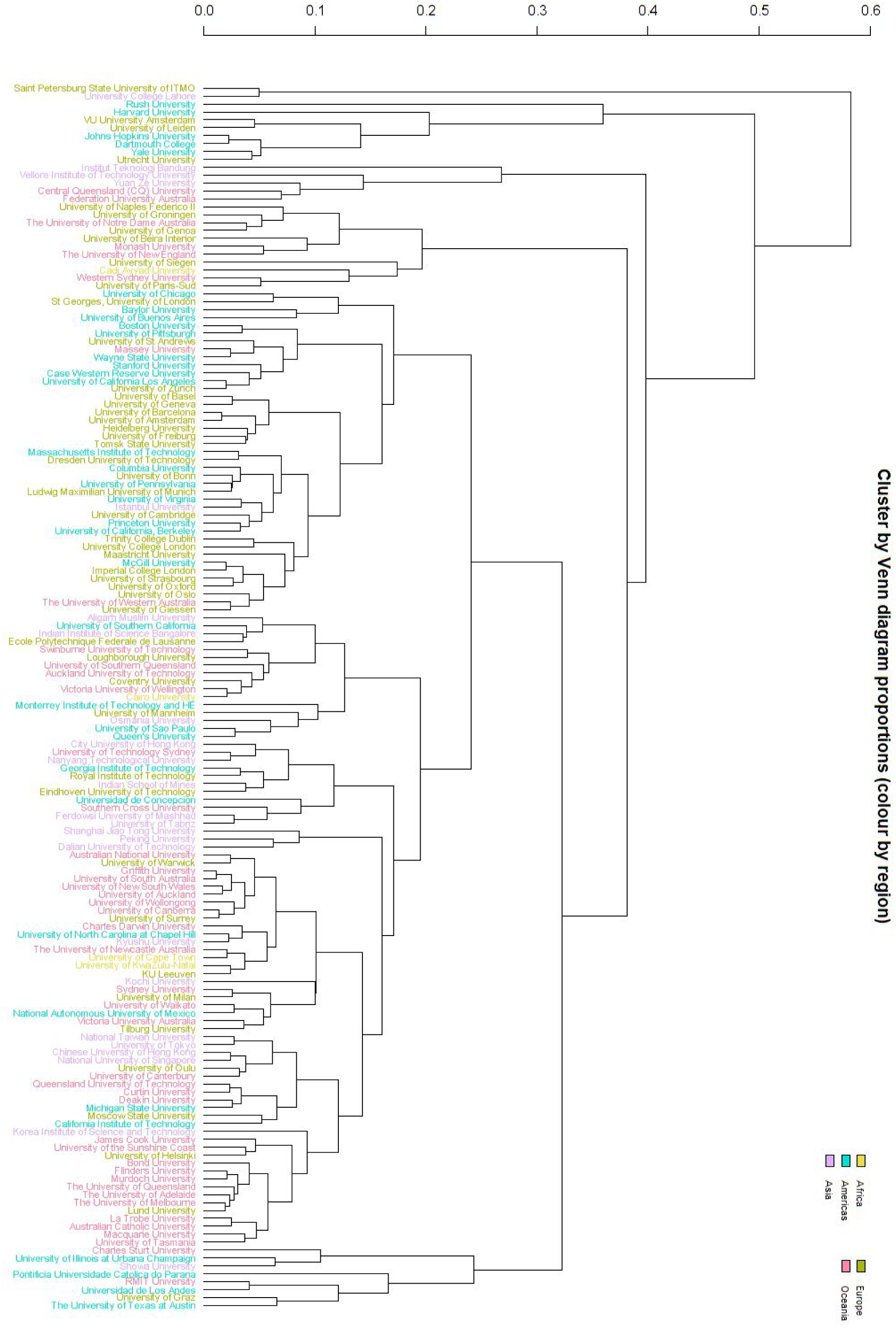
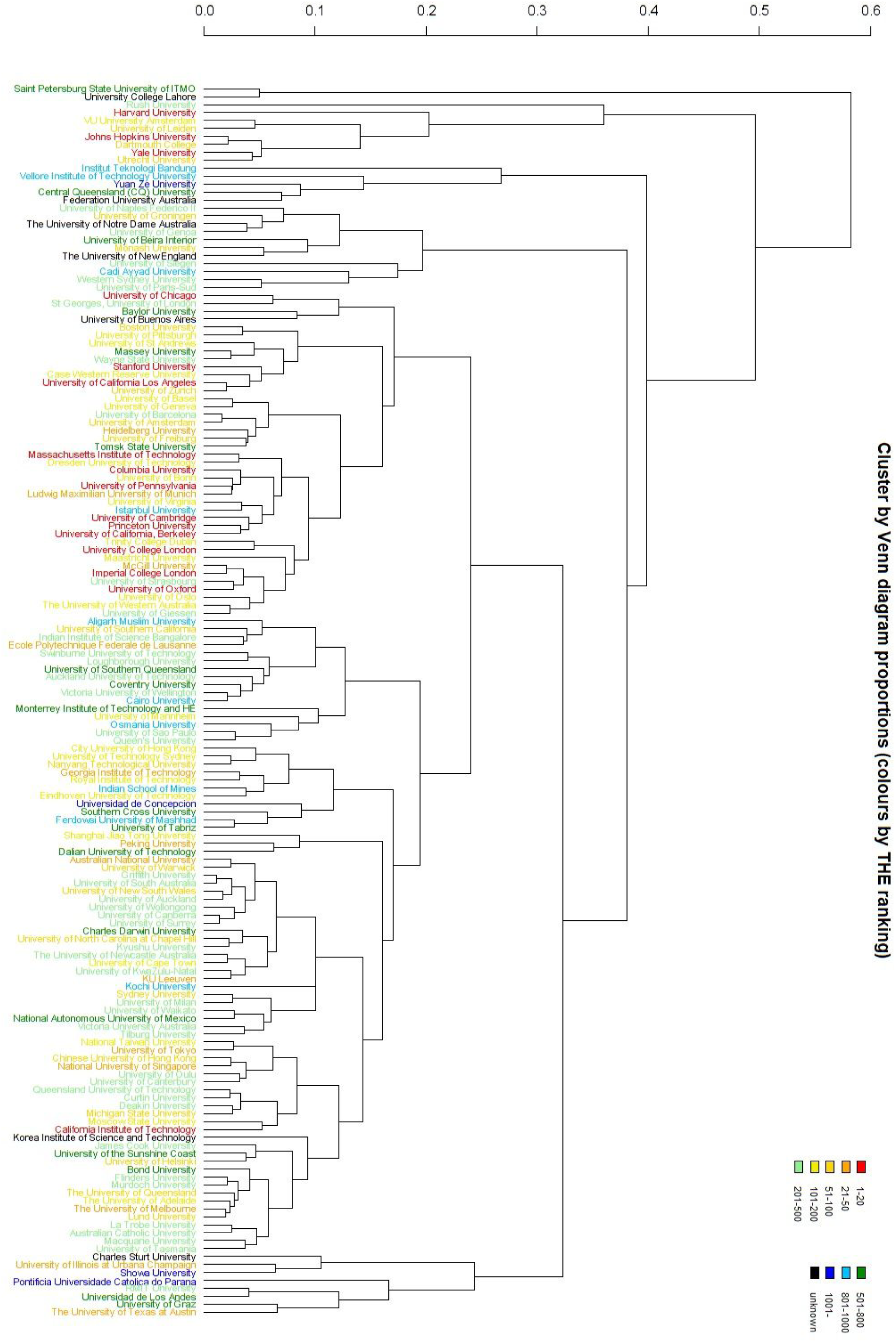
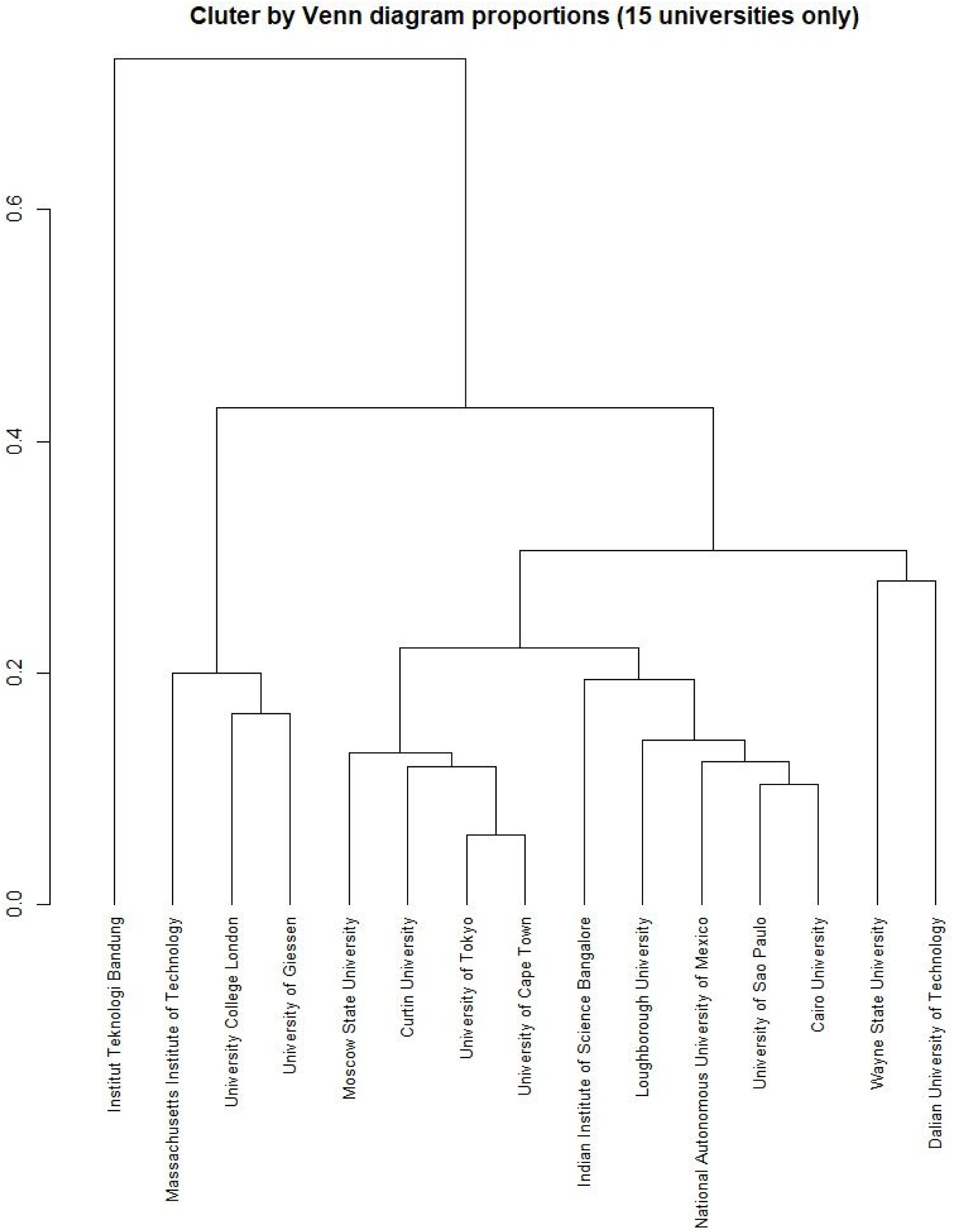

## Appendix 7 Standout cases in publication year comparison

Appendix 7: Standout cases in publication year comparison
|  | 2016 Scopus DOIs matched against MSA DOIs in other years |  |  |  | 2016 MSA DOIs matched against Scopus DOIs in other years |  |  |  |
| --- | --- | --- | --- | --- | --- | --- | --- | --- |
| Year diff. | -2 | -1 | +1 | +2 | -2 | -1 | +1 | +2 |
| Cairo | 0.1 | 5.5 | 0.3 | 0.0 | 0.0 | 0.5 | 4.2 | 0.3 |
| IISC | 0.2 | 4.6 | 0.3 | 0.0 | 0.1 | 0.5 | 5.1 | 0.0 |
| ITB | 0.0 | 24.5 | 0.1 | 0.0 | 0.0 | 0.4 | 25.0 | 0.0 |

## Appendix 8 Document types for individual institutions in 2016

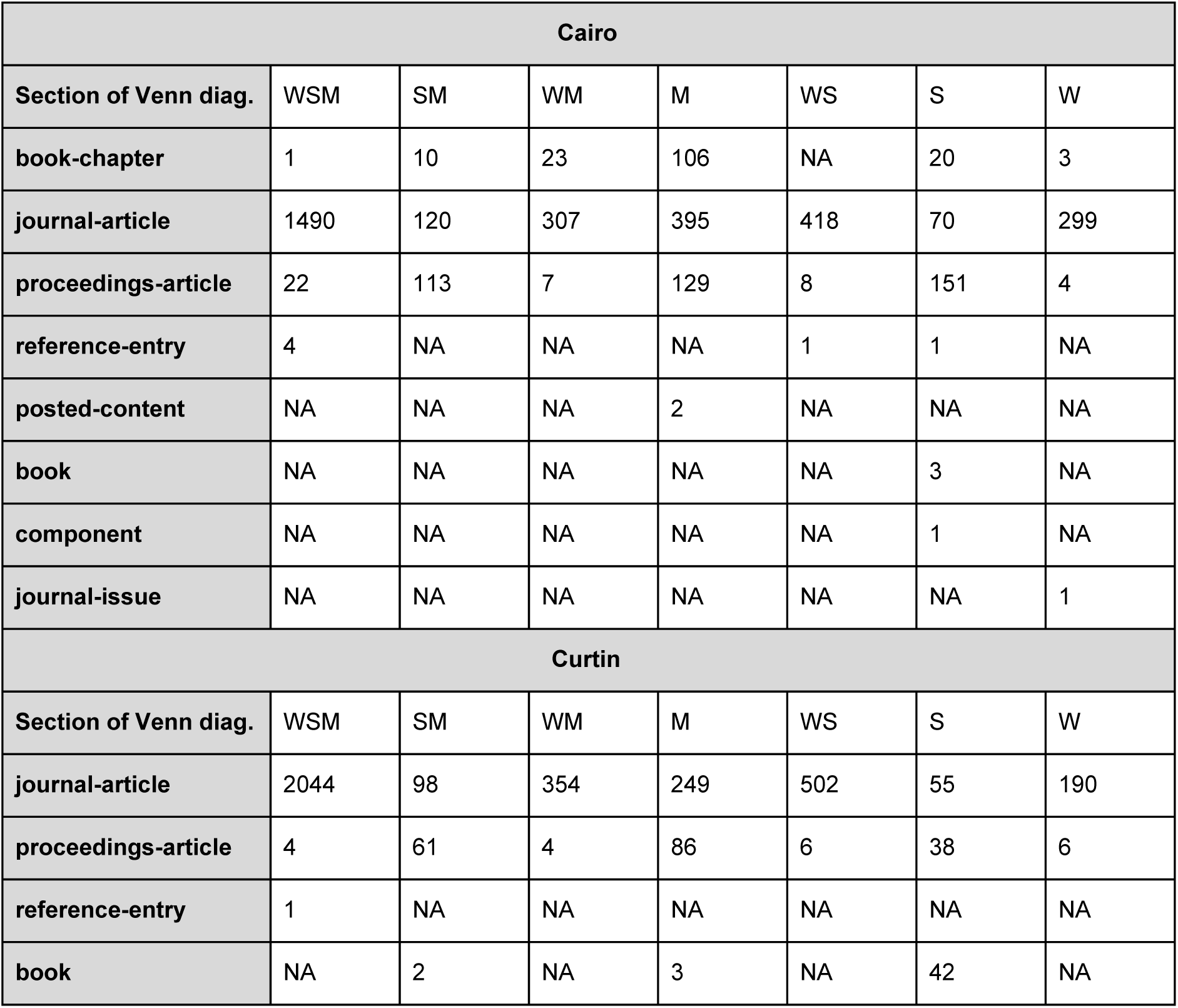
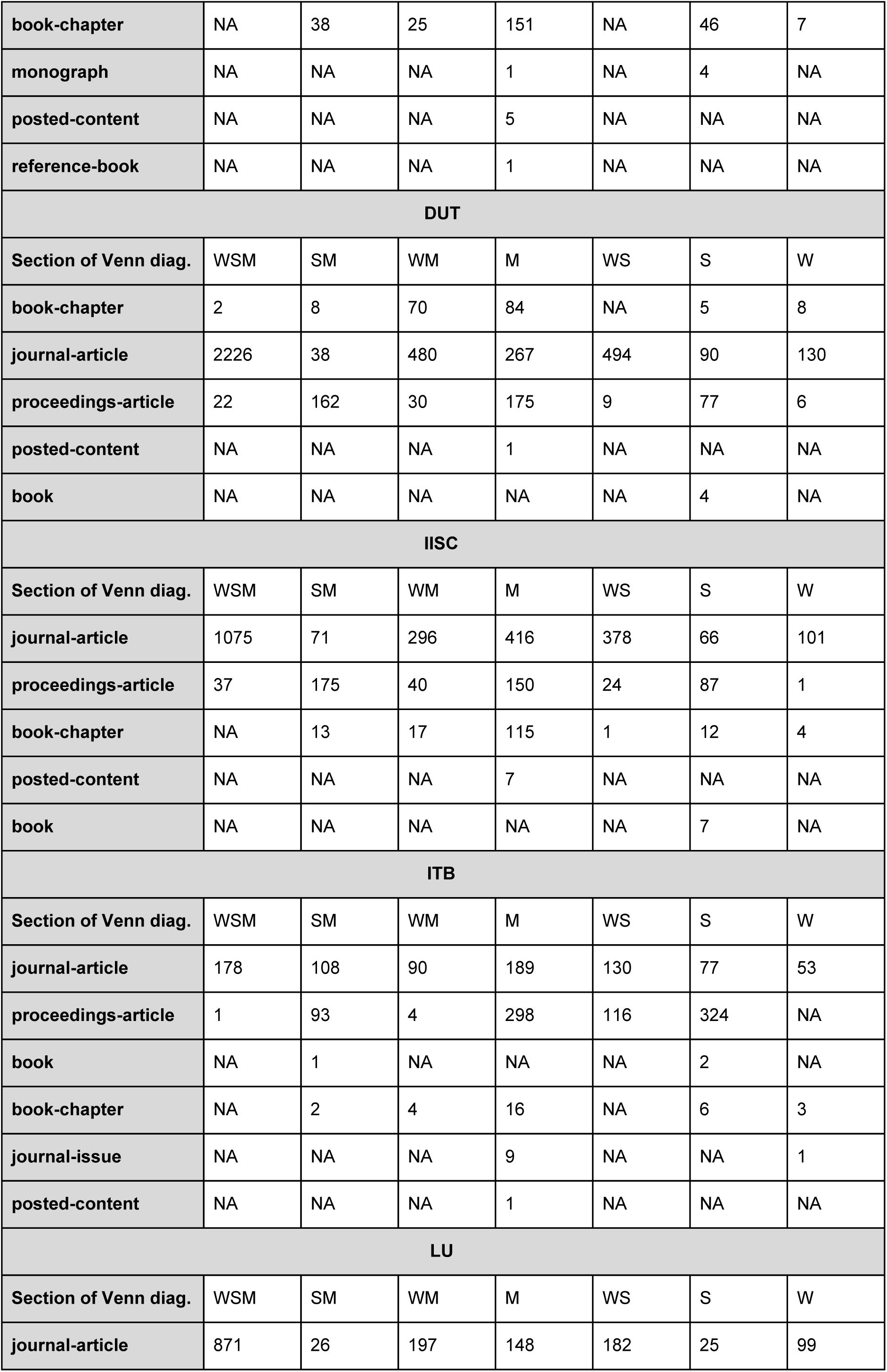
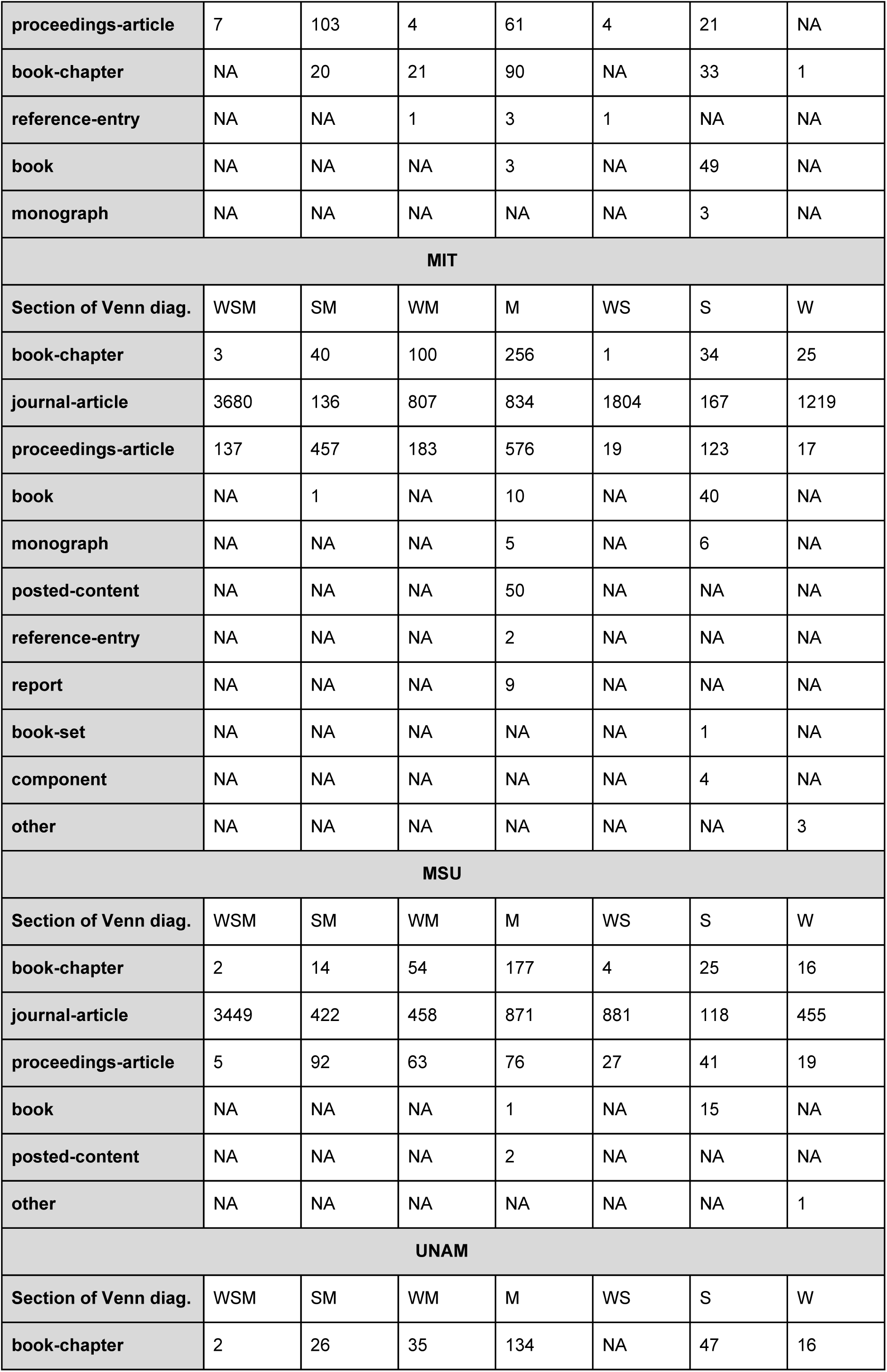
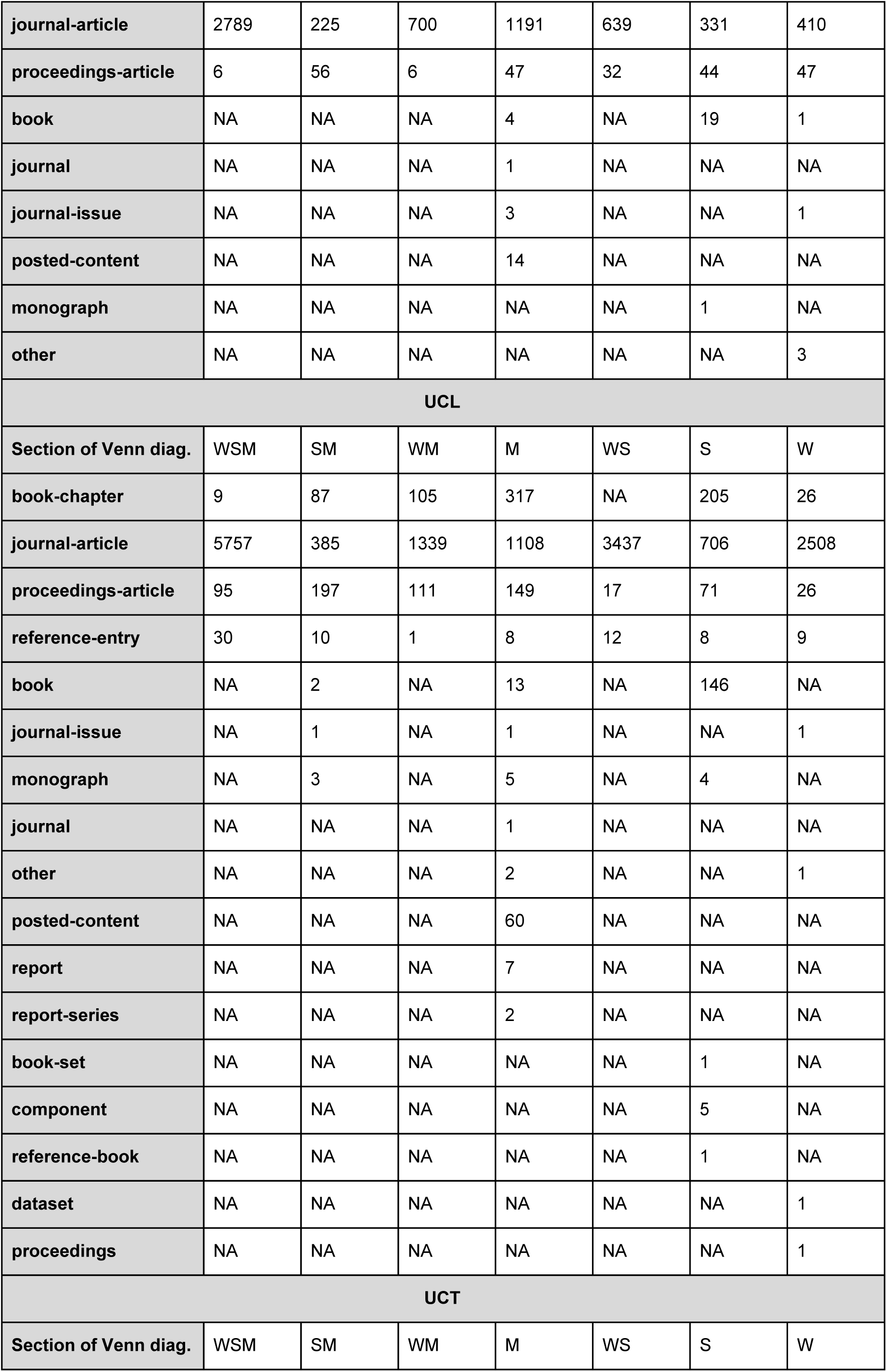
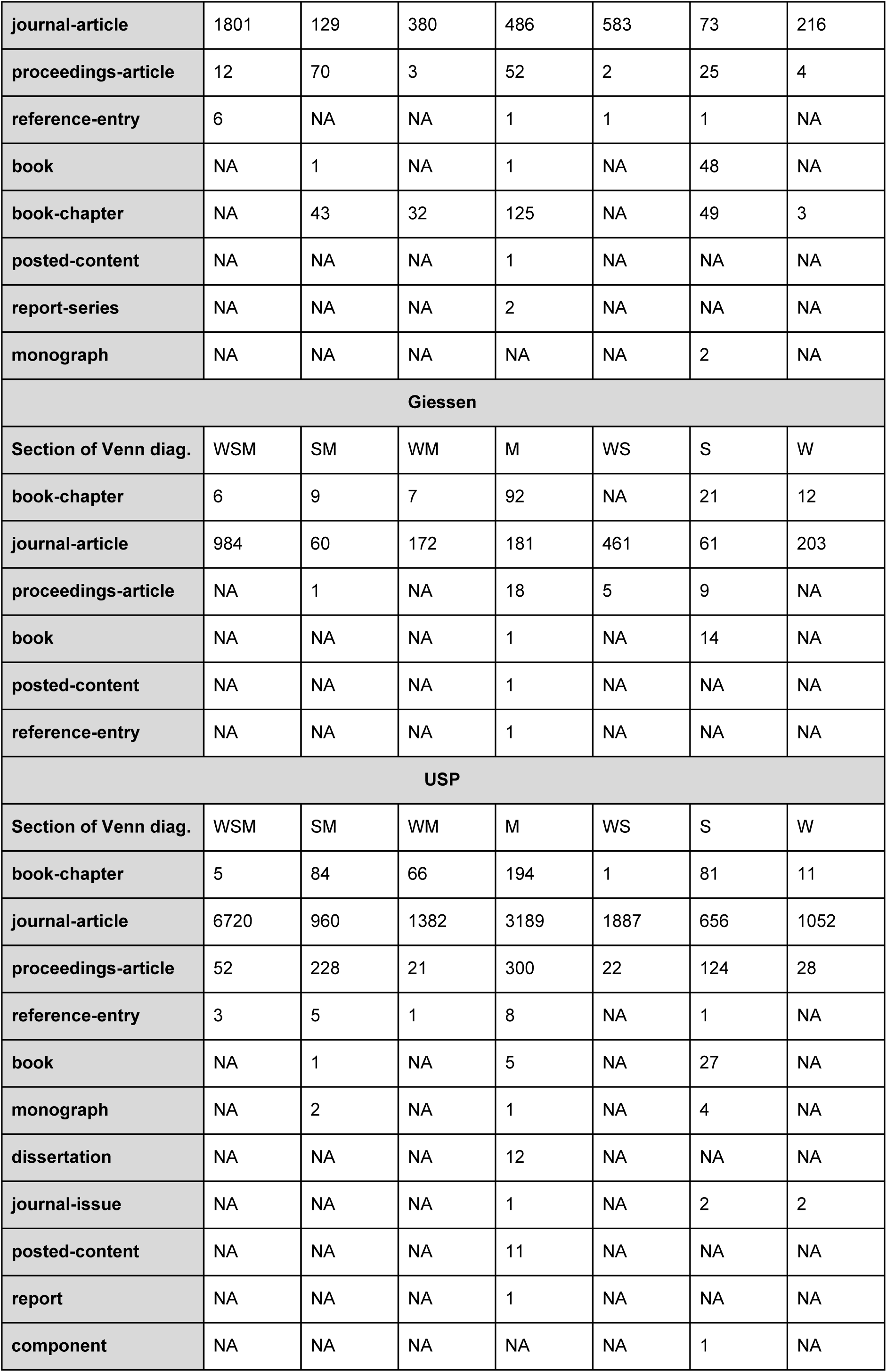
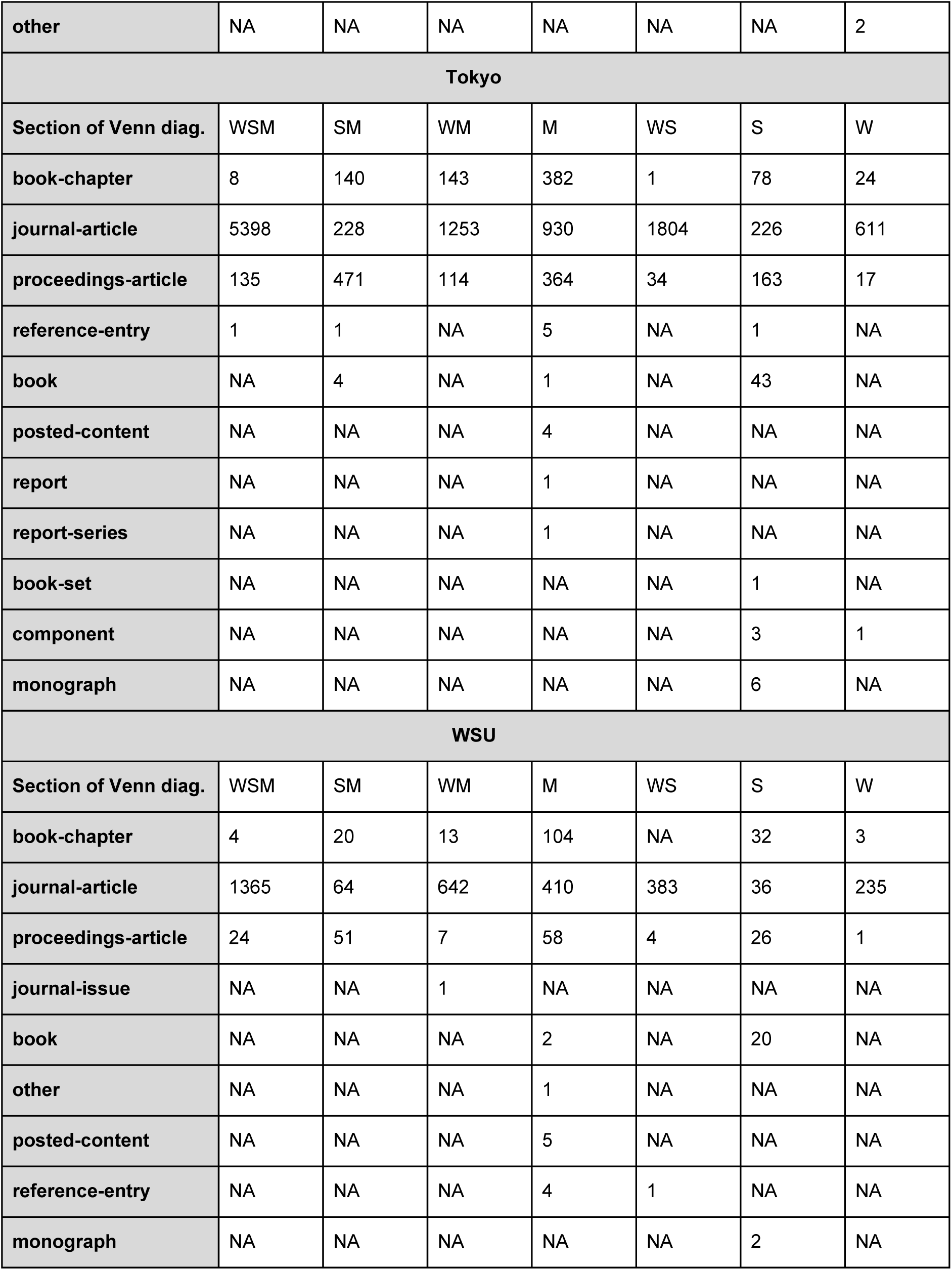

Appendix 8: Document types for individual institutions in 2016
| Cairo |  |  |  |  |  |  |  |
| --- | --- | --- | --- | --- | --- | --- | --- |
| Section of Venn diag. | WSM | SM | WM | M | WS | S | W |
| book-chapter | 1 | 10 | 23 | 106 | NA | 20 | 3 |
| journal-article | 1490 | 120 | 307 | 395 | 418 | 70 | 299 |
| proceedings-article | 22 | 113 | 7 | 129 | 8 | 151 | 4 |
| reference-entry | 4 | NA | NA | NA | 1 | 1 | NA |
| posted-content | NA | NA | NA | 2 | NA | NA | NA |
| book | NA | NA | NA | NA | NA | 3 | NA |
| component | NA | NA | NA | NA | NA | 1 | NA |
| journal-issue | NA | NA | NA | NA | NA | NA | 1 |
| Curtin |  |  |  |  |  |  |  |
| Section of Venn diag. | WSM | SM | WM | M | WS | S | W |
| journal-article | 2044 | 98 | 354 | 249 | 502 | 55 | 190 |
| proceedings-article | 4 | 61 | 4 | 86 | 6 | 38 | 6 |
| reference-entry | 1 | NA | NA | NA | NA | NA | NA |
| book | NA | 2 | NA | 3 | NA | 42 | NA |
| **book-chapter** | NA | 38 | 25 | 151 | NA | 46 | 7 |
| **monograph** | NA | NA | NA | 1 | NA | 4 | NA |
| **posted-content** | NA | NA | NA | 5 | NA | NA | NA |
| **reference-book** | NA | NA | NA | 1 | NA | NA | NA |
| **DUT** |  |  |  |  |  |  |  |
| **Section of Venn diag.** | WSM | SM | WM | M | WS | S | W |
| **book-chapter** | 2 | 8 | 70 | 84 | NA | 5 | 8 |
| **journal-article** | 2226 | 38 | 480 | 267 | 494 | 90 | 130 |
| **proceedings-article** | 22 | 162 | 30 | 175 | 9 | 77 | 6 |
| **posted-content** | NA | NA | NA | 1 | NA | NA | NA |
| **book** | NA | NA | NA | NA | NA | 4 | NA |
| **IISC** |  |  |  |  |  |  |  |
| **Section of Venn diag.** | WSM | SM | WM | M | WS | S | W |
| **journal-article** | 1075 | 71 | 296 | 416 | 378 | 66 | 101 |
| **proceedings-article** | 37 | 175 | 40 | 150 | 24 | 87 | 1 |
| **book-chapter** | NA | 13 | 17 | 115 | 1 | 12 | 4 |
| **posted-content** | NA | NA | NA | 7 | NA | NA | NA |
| **book** | NA | NA | NA | NA | NA | 7 | NA |
| **ITB** |  |  |  |  |  |  |  |
| **Section of Venn diag.** | WSM | SM | WM | M | WS | S | W |
| **journal-article** | 178 | 108 | 90 | 189 | 130 | 77 | 53 |
| **proceedings-article** | 1 | 93 | 4 | 298 | 116 | 324 | NA |
| **book** | NA | 1 | NA | NA | NA | 2 | NA |
| **book-chapter** | NA | 2 | 4 | 16 | NA | 6 | 3 |
| **journal-issue** | NA | NA | NA | 9 | NA | NA | 1 |
| **posted-content** | NA | NA | NA | 1 | NA | NA | NA |
| **LU** |  |  |  |  |  |  |  |
| **Section of Venn diag.** | WSM | SM | WM | M | WS | S | W |
| **journal-article** | 871 | 26 | 197 | 148 | 182 | 25 | 99 |
| **proceedings-article** | 7 | 103 | 4 | 61 | 4 | 21 | NA |
| **book-chapter** | NA | 20 | 21 | 90 | NA | 33 | 1 |
| **reference-entry** | NA | NA | 1 | 3 | 1 | NA | NA |
| **book** | NA | NA | NA | 3 | NA | 49 | NA |
| **monograph** | NA | NA | NA | NA | NA | 3 | NA |
| **MIT** |  |  |  |  |  |  |  |
| **Section of Venn diag.** | WSM | SM | WM | M | WS | S | W |
| **book-chapter** | 3 | 40 | 100 | 256 | 1 | 34 | 25 |
| **journal-article** | 3680 | 136 | 807 | 834 | 1804 | 167 | 1219 |
| **proceedings-article** | 137 | 457 | 183 | 576 | 19 | 123 | 17 |
| **book** | NA | 1 | NA | 10 | NA | 40 | NA |
| **monograph** | NA | NA | NA | 5 | NA | 6 | NA |
| **posted-content** | NA | NA | NA | 50 | NA | NA | NA |
| **reference-entry** | NA | NA | NA | 2 | NA | NA | NA |
| **report** | NA | NA | NA | 9 | NA | NA | NA |
| **book-set** | NA | NA | NA | NA | NA | 1 | NA |
| **component** | NA | NA | NA | NA | NA | 4 | NA |
| **other** | NA | NA | NA | NA | NA | NA | 3 |
| **MSU** |  |  |  |  |  |  |  |
| **Section of Venn diag.** | WSM | SM | WM | M | WS | S | W |
| **book-chapter** | 2 | 14 | 54 | 177 | 4 | 25 | 16 |
| **journal-article** | 3449 | 422 | 458 | 871 | 881 | 118 | 455 |
| **proceedings-article** | 5 | 92 | 63 | 76 | 27 | 41 | 19 |
| **book** | NA | NA | NA | 1 | NA | 15 | NA |
| **posted-content** | NA | NA | NA | 2 | NA | NA | NA |
| **other** | NA | NA | NA | NA | NA | NA | 1 |
| **UNAM** |  |  |  |  |  |  |  |
| **Section of Venn diag.** | WSM | SM | WM | M | WS | S | W |
| **book-chapter** | 2 | 26 | 35 | 134 | NA | 47 | 16 |
| **journal-article** | 2789 | 225 | 700 | 1191 | 639 | 331 | 410 |
| **proceedings-article** | 6 | 56 | 6 | 47 | 32 | 44 | 47 |
| **book** | NA | NA | NA | 4 | NA | 19 | 1 |
| **journal** | NA | NA | NA | 1 | NA | NA | NA |
| **journal-issue** | NA | NA | NA | 3 | NA | NA | 1 |
| **posted-content** | NA | NA | NA | 14 | NA | NA | NA |
| **monograph** | NA | NA | NA | NA | NA | 1 | NA |
| **other** | NA | NA | NA | NA | NA | NA | 3 |
| **UCL** |  |  |  |  |  |  |  |
| **Section of Venn diag.** | WSM | SM | WM | M | WS | S | W |
| **book-chapter** | 9 | 87 | 105 | 317 | NA | 205 | 26 |
| **journal-article** | 5757 | 385 | 1339 | 1108 | 3437 | 706 | 2508 |
| **proceedings-article** | 95 | 197 | 111 | 149 | 17 | 71 | 26 |
| **reference-entry** | 30 | 10 | 1 | 8 | 12 | 8 | 9 |
| **book** | NA | 2 | NA | 13 | NA | 146 | NA |
| **journal-issue** | NA | 1 | NA | 1 | NA | NA | 1 |
| **monograph** | NA | 3 | NA | 5 | NA | 4 | NA |
| **journal** | NA | NA | NA | 1 | NA | NA | NA |
| **other** | NA | NA | NA | 2 | NA | NA | 1 |
| **posted-content** | NA | NA | NA | 60 | NA | NA | NA |
| **report** | NA | NA | NA | 7 | NA | NA | NA |
| **report-series** | NA | NA | NA | 2 | NA | NA | NA |
| **book-set** | NA | NA | NA | NA | NA | 1 | NA |
| **component** | NA | NA | NA | NA | NA | 5 | NA |
| **reference-book** | NA | NA | NA | NA | NA | 1 | NA |
| **dataset** | NA | NA | NA | NA | NA | NA | 1 |
| **proceedings** | NA | NA | NA | NA | NA | NA | 1 |
| **UCT** |  |  |  |  |  |  |  |
| **Section of Venn diag.** | WSM | SM | WM | M | WS | S | W |
| **journal-article** | 1801 | 129 | 380 | 486 | 583 | 73 | 216 |
| **proceedings-article** | 12 | 70 | 3 | 52 | 2 | 25 | 4 |
| **reference-entry** | 6 | NA | NA | 1 | 1 | 1 | NA |
| **book** | NA | 1 | NA | 1 | NA | 48 | NA |
| **book-chapter** | NA | 43 | 32 | 125 | NA | 49 | 3 |
| **posted-content** | NA | NA | NA | 1 | NA | NA | NA |
| **report-series** | NA | NA | NA | 2 | NA | NA | NA |
| **monograph** | NA | NA | NA | NA | NA | 2 | NA |
| **Giessen** |  |  |  |  |  |  |  |
| **Section of Venn diag.** | WSM | SM | WM | M | WS | S | W |
| **book-chapter** | 6 | 9 | 7 | 92 | NA | 21 | 12 |
| **journal-article** | 984 | 60 | 172 | 181 | 461 | 61 | 203 |
| **proceedings-article** | NA | 1 | NA | 18 | 5 | 9 | NA |
| **book** | NA | NA | NA | 1 | NA | 14 | NA |
| **posted-content** | NA | NA | NA | 1 | NA | NA | NA |
| **reference-entry** | NA | NA | NA | 1 | NA | NA | NA |
| **USP** |  |  |  |  |  |  |  |
| **Section of Venn diag.** | WSM | SM | WM | M | WS | S | W |
| **book-chapter** | 5 | 84 | 66 | 194 | 1 | 81 | 11 |
| **journal-article** | 6720 | 960 | 1382 | 3189 | 1887 | 656 | 1052 |
| **proceedings-article** | 52 | 228 | 21 | 300 | 22 | 124 | 28 |
| **reference-entry** | 3 | 5 | 1 | 8 | NA | 1 | NA |
| **book** | NA | 1 | NA | 5 | NA | 27 | NA |
| **monograph** | NA | 2 | NA | 1 | NA | 4 | NA |
| **dissertation** | NA | NA | NA | 12 | NA | NA | NA |
| **journal-issue** | NA | NA | NA | 1 | NA | 2 | 2 |
| **posted-content** | NA | NA | NA | 11 | NA | NA | NA |
| **report** | NA | NA | NA | 1 | NA | NA | NA |
| **component** | NA | NA | NA | NA | NA | 1 | NA |
| other | NA | NA | NA | NA | NA | NA | 2 |
| **Tokyo** |  |  |  |  |  |  |  |
| **Section of Venn diag.** | WSM | SM | WM | M | WS | S | W |
| book-chapter | 8 | 140 | 143 | 382 | 1 | 78 | 24 |
| journal-article | 5398 | 228 | 1253 | 930 | 1804 | 226 | 611 |
| proceedings-article | 135 | 471 | 114 | 364 | 34 | 163 | 17 |
| reference-entry | 1 | 1 | NA | 5 | NA | 1 | NA |
| book | NA | 4 | NA | 1 | NA | 43 | NA |
| posted-content | NA | NA | NA | 4 | NA | NA | NA |
| report | NA | NA | NA | 1 | NA | NA | NA |
| report-series | NA | NA | NA | 1 | NA | NA | NA |
| book-set | NA | NA | NA | NA | NA | 1 | NA |
| component | NA | NA | NA | NA | NA | 3 | 1 |
| monograph | NA | NA | NA | NA | NA | 6 | NA |
| **WSU** |  |  |  |  |  |  |  |
| **Section of Venn diag.** | WSM | SM | WM | M | WS | S | W |
| book-chapter | 4 | 20 | 13 | 104 | NA | 32 | 3 |
| journal-article | 1365 | 64 | 642 | 410 | 383 | 36 | 235 |
| proceedings-article | 24 | 51 | 7 | 58 | 4 | 26 | 1 |
| journal-issue | NA | NA | 1 | NA | NA | NA | NA |
| book | NA | NA | NA | 2 | NA | 20 | NA |
| other | NA | NA | NA | 1 | NA | NA | NA |
| posted-content | NA | NA | NA | 5 | NA | NA | NA |
| reference-entry | NA | NA | NA | 4 | 1 | NA | NA |
| monograph | NA | NA | NA | NA | NA | 2 | NA |

## Appendix 9

(see Brookes-Kenworthy et al, 2019, for a summary and raw forms of data collected via the manual cross-validation process).

Here we proceed to validate the sources against each other, at the institutional level. The three charts in Figure A9.1 show the percentages of sampled DOIs from one source that are found in the metadata of the other two sources (i.e., a “DOI match”). This presents some indication of the amount of DOIs that are actually covered by another source, but simply did not link to the target affiliation in our data collection and validation processes (either the object did not show up in affiliation search or the object’s DOI is not passed through collection process via APIs)^56^.

**Figure A9.1:**
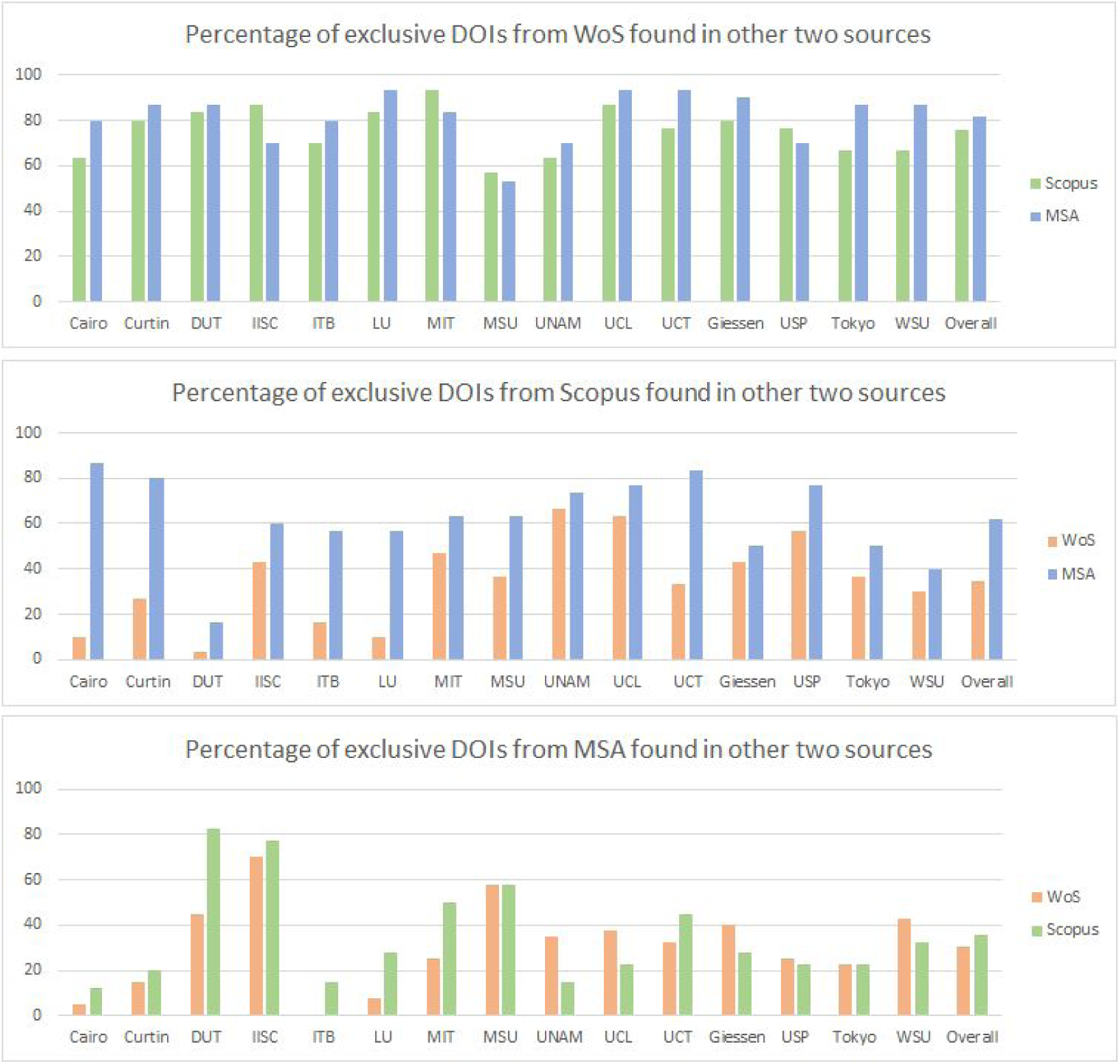
Percentage of exclusive DOIs from each source that are found in the metadata (matching DOI only) of the other two sources.

The top chart indicates that a relatively high proportion of DOIs from WoS are indexed in both Scopus and MSA. This implies that Scopus and MSA actually have good coverages of DOIs that initially seemed exclusive to WoS, but they were not assigned to the target affiliations by Scopus and MSA. In contrast, much less of DOIs from MSA can be found in WoS and Scopus (bottom chart). As for DOIs from Scopus (middle chart), relatively high proportion of them can be found in MSA, but much less so in WoS. An extreme exception is DUT, where neither WoS nor MSA have very high coverage of DOIs exclusively from Scopus (it may be worth recalling we saw earlier that Scopus had exclusive coverage of many DOIs registered with Chinese DOI agencies). Overall, it appears that MSA has a broader coverage of all DOIs, but the completeness of affiliation metadata is lower.

While a DOI may be missing from a database, the object to which the DOI is assigned to may actually still be in the database, i.e., the DOI was simply not recorded in the metadata of the research object. Hence, we also performed “title match” (instead of matching DOI strings) across sources. For each sampled DOI, we check whether the corresponding document title can be found in the other two sources. These are summarised in the three charts of Figure A9.2.

**Figure A9.2:**
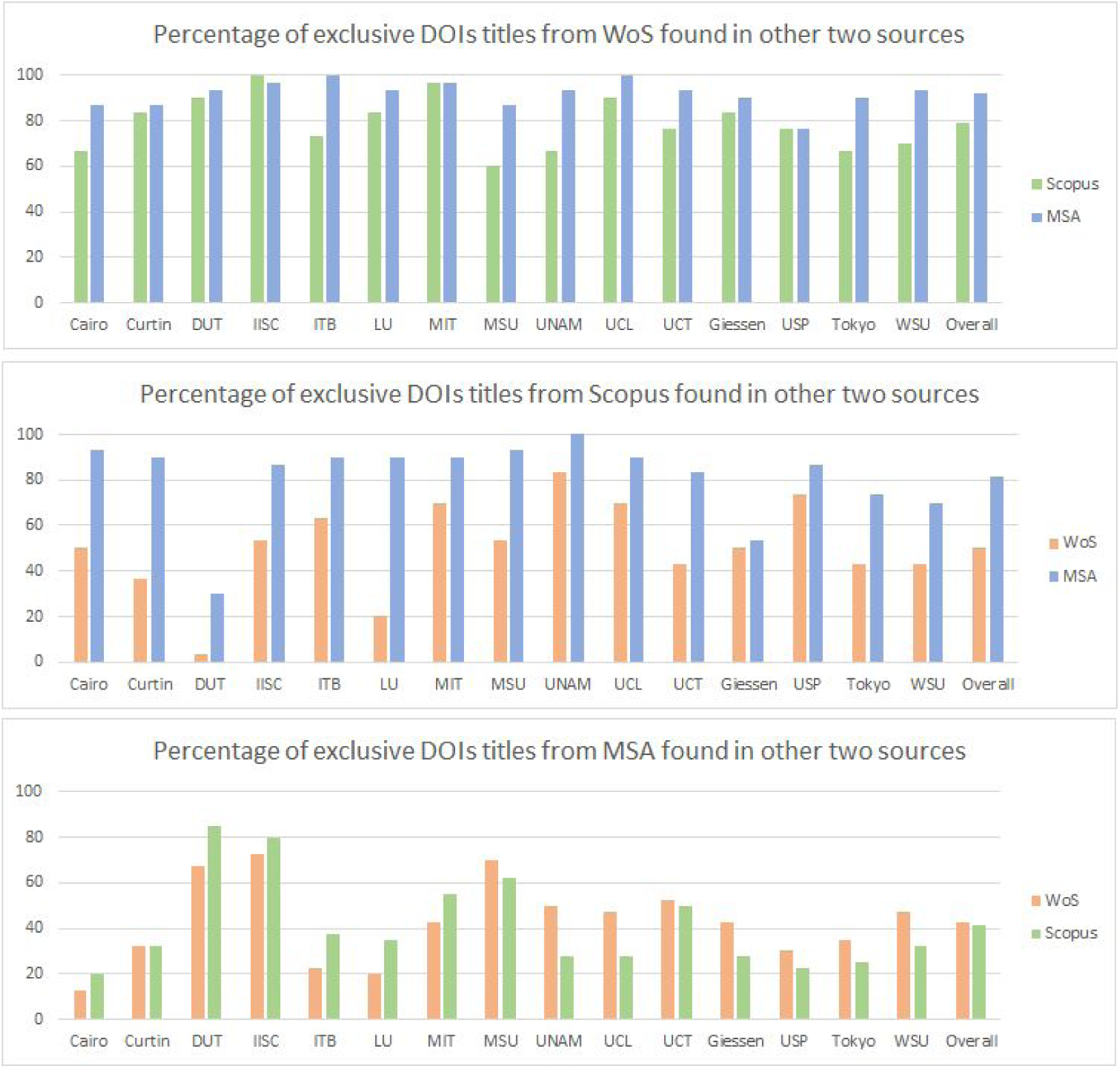
Percentage of exclusive DOIs from each source that have their corresponding title found in the other two sources.

All percentages in Figure A9.2 are equal or greater than the corresponding percentages in Figure A9.1. This is because for all cases for which the DOI was found, the corresponding title was also found (i.e., various objects have correct titles but no record of their DOIs, not the other way around). The results here also further highlight the extent of MSA’s high coverage of objects related to DOIs initially appeared to be exclusively indexed by the other two sources. Otherwise, the general pattern is similar to what we observed earlier, with WoS titles mostly covered by both MSA and Scopus, MSA having more coverage (than WoS) of Scopus titles, and both Scopus and WoS have relatively lower coverage of titles from MSA.

Having the correct affiliation recorded in metadata is not necessarily the same as having the correct affiliation linkage (e.g., an object may have the correct metadata, but does not show up in the affiliation search). As a way to gain some insight into the degree of this issue, we match the affiliation across sources. When two sources both match a DOI to its target affiliation, we refer this to as an “affiliation match”. Figure A9.3 demonstrates the findings again via three different charts. Each bar represents the percentage DOIs from one source having title match and affiliation match with another source. For example, the green bars in the top chart of Figure A9.3 denote percentages of those exclusive WoS titles that were found in Scopus and also have affiliation metadata in Scopus that match the target affiliations.

**Figure A9.3:**
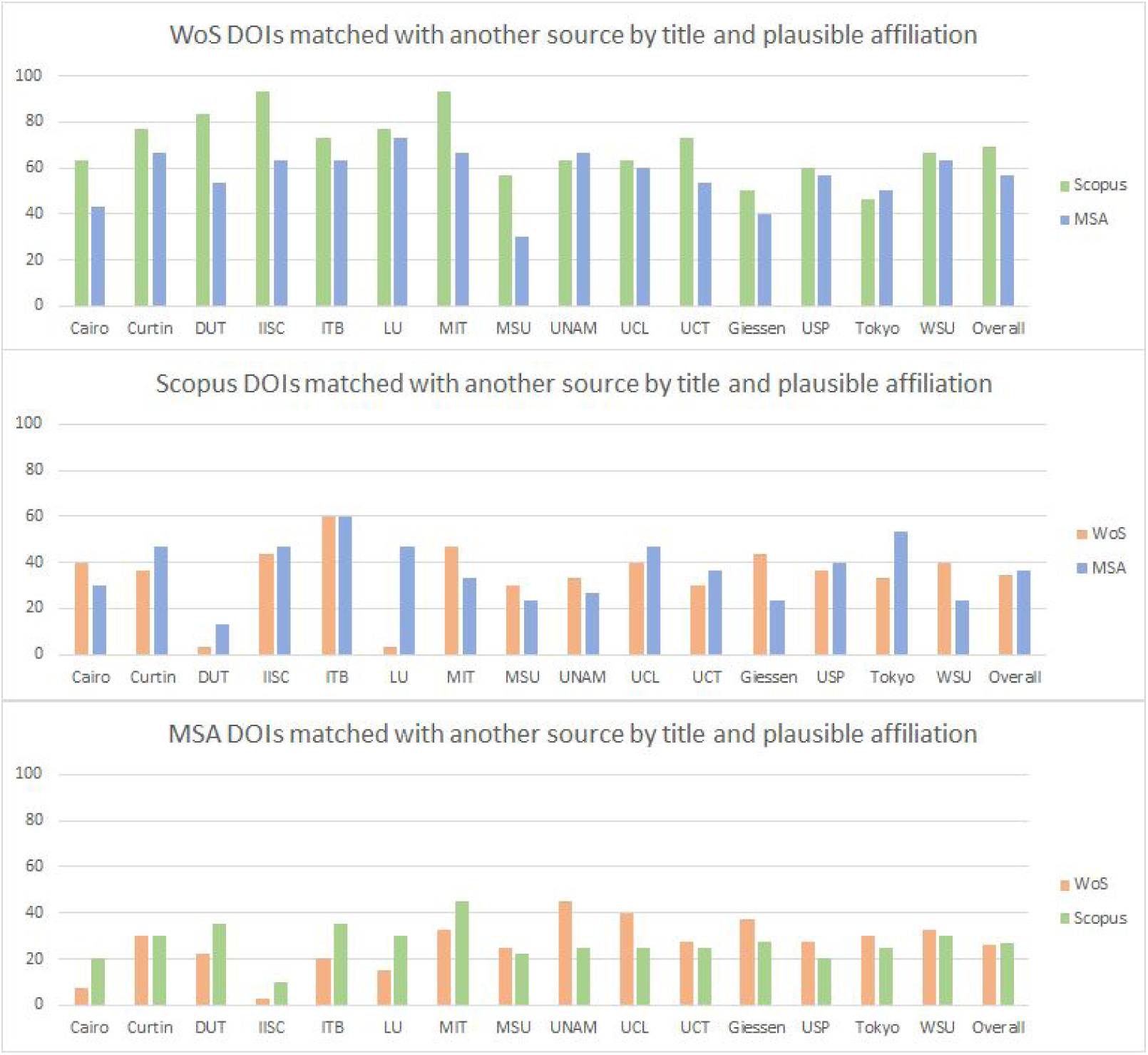
Title and plausible affiliation found in another source.

It certainly appears that more WoS DOIs actually have title and affiliation matches in the other two sources. One can also note the decrease of percentages when compared to Figure A9.2. This is a clear indication that many of these titles were simply not assigned to their target affiliations by the contrasting sources. However, the numbers here also include title matches that does not necessarily have DOI matches. This means we cannot tell how many of these should have been collected from the contrasting sources via our data collection process.

**Figure A9.4:**
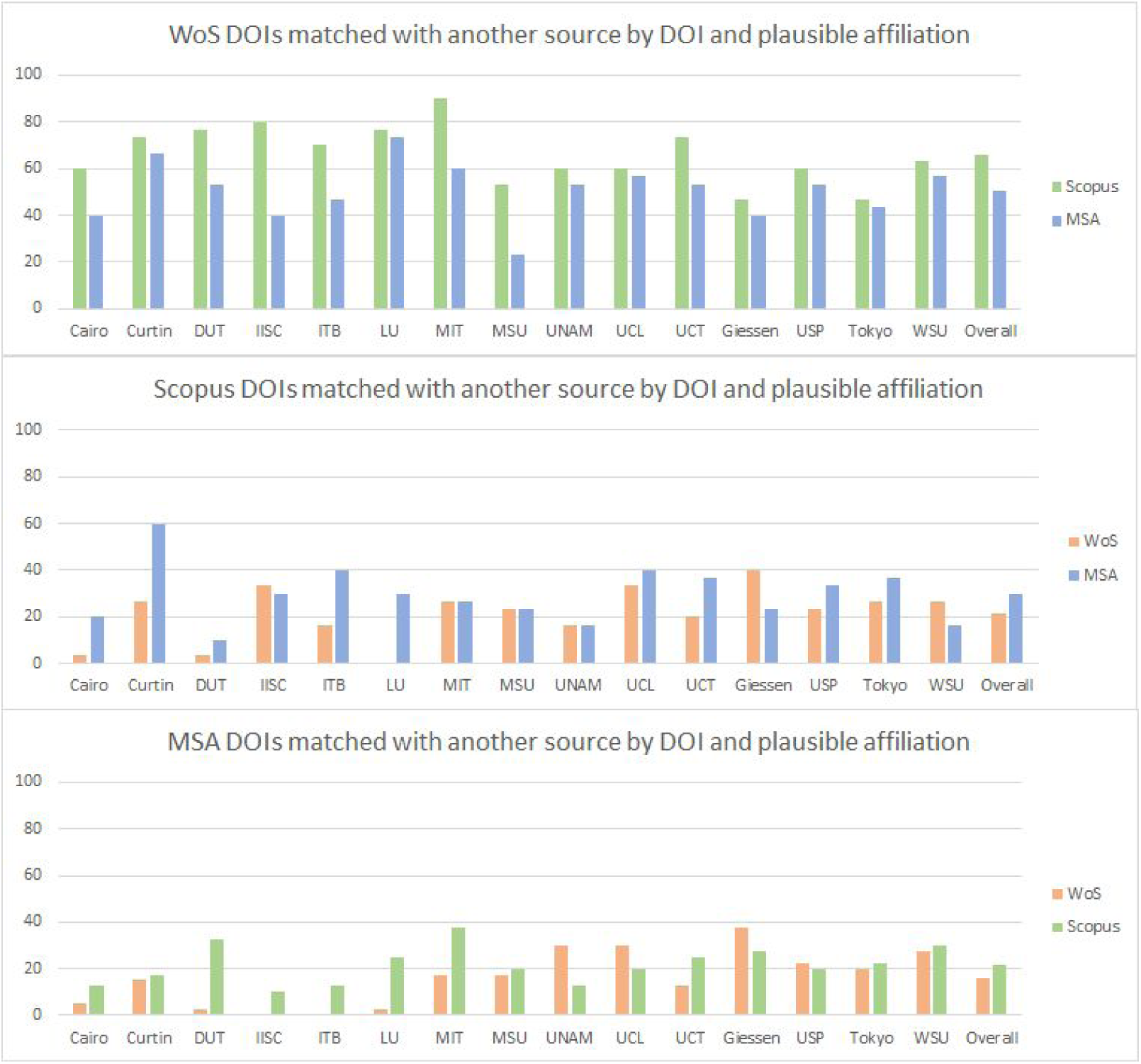
DOI and plausible affiliation found in another source.

Hence, we now filter down to objects that have both DOI matches and affiliation matches. These are presented in Figure A9.4 and are indications of numbers of DOIs that our data collection process should have captured (but did not) from each contrasting source, given they have DOIs and are plausibly affiliated to target affiliations. The reason for these to be missing from our collection process is likely^57^ to be that the affiliation linkages are broken. This could include various reasons but the most prominent one seems to be that the metadata (as per source website) is not synchronized with the API returns we gathered.

More WoS DOIs have both DOI and affiliation match by other sources, in contrast to those of Scopus and MSA.

## Appendix 10 Some examples of DOI errors in WoS, Scopus and MSA

Here we give a few examples^58^ of errors we found that are related to metadata recordings of DOIs:

- **10.1109/AUPEC.2016.07749327**: This is a conference paper affiliated to Curtin University via Scopus. However, this is not a valid DOI (according to doi.org and Crossref). A title search reveals that there is an extra “0” in the DOI and the correct DOI should be 10.1109/AUPEC.2016.7749327.
- **10.1166/jnn.2016.11757**: This is a journal paper affiliated to Loughborough University via WoS. This is not a valid DOI and the correct DOI should be 10.1166/jnn.2016.1175 (i.e., an extra digit of “7” was recorded at the end of the string).
- **10.11113/jt.v78.9016:** This is a journal article affiliated to Institut Teknologi Bandung via MSA. The DOI is found on the article website (not in the article itself). However, this is not a valid DOI as per doi.org.

## Footnotes

1 See https://www.topuniversities.com/qs-world-university-rankings/methodology, https://www.timeshighereducation.com/world-university-rankings/methodology-world-university-rankings-2018, and http://www.shanghairanking.com/ARWU-Methodology-2017.html.

2 See, for example, http://www.bbk.ac.uk/news/league-tables.

3 We have selected WoS, Scopus and MSA for our analysis because they provide structured metadata handling and comprehensive API search functions. GS is not considered due to difficulties in metadata handling and lack of API support, but it may be of interest for examination in future work (especially given the apparent large scale of coverage).

4 https://unpaywall.org/

5 https://opencitations.net/

6 As per website search or report on 7 August 2018. Numbers reported are not necessarily the same as total number of user accessible records. For estimates of user accessible records, see Gusenbauer (2018).

7 As permitted through the advanced search functions in WoS and Scopus on 7 August 2018.

8 While this article was being prepared, Elsevier announced their agreement to use Unpaywall data, see https://www.elsevier.com/connect/elsevier-impactstory-agreement-will-make-open-access-articles-easier-to-find-on-scopus and later implemented it https://blog.scopus.com/posts/scopus-makes-millions-of-open-access-articles-easily-discoverable

9 See https://dev.labs.cognitive.microsoft.com/products/5636d970e597ed0690ac1b3f

10 See https://azure.microsoft.com/en-au/pricing/details/cognitive-services/academic-knowledge-api/

11 See https://www.microsoft.com/en-us/research/project/academic/articles/sharpening-insights-into-the-innovation-landscape-with-a-new-approach-to-patents/

12 Data as of 15 August 2019.

13 Publication year defined as per source.

14 This is for all databases in WoS. Counts were obtained by querying for all research areas under each of the five broad categories (as defined by WoS) using the Advanced Search function on WoS, as at 3 August 2018.

15 This is for the core databases in WoS. Counts were obtained by querying for all research areas under each of the five broad categories (as defined by WoS) using the Advanced Search function on WoS, as at 8 August 2018.

16 Obtained by querying for each broad subject area (as defined by Scopus) through Scopus’ Advanced Search option, as at 3 August 2018.

17 MSA did not seem to have broadly defined disciplines. Counts for the 19 top-level fields of study were obtained from the Topics Analytics page (on 3 August 2018). Then we sorted their detailed disciplines into broader ones (roughly following those in WoS) as follows: Health Sciences = Medicine; Physical Sciences = Chemistry, Engineering, Computer Science, Physics, Materials Science, Mathematics, Geology; Life Sciences = Biology, Environmental Science; Social Sciences = Psychology, Geography, Sociology, Political Science, Business, Economics; Arts & Humanities = History, Art, Philosophy.

18 See Appendix 1.

19 See Appendix 2 for a list of WoS databases accessed in this study.

20 This implied a Crossref/Unpaywall DOI that does not have any citation links in OpenCitations is assumed to have zero citation for this study.

21 We use affiliation IDs from the Global Research Identifier Database (GRID, https://www.grid.ac/) as the standardised identifier for each institution. These are mapped to IDs and search terms in WoS, Scopus and MSA in Appendix 1.

22 This cross-validation process was carried out manually by a data wrangler, on a part-time basis over a few months, for which online data was accessed from 18 December 2018 to 20 May 2019.

23 Using the Crossref API for agency information.

24 See https://unpaywall.org/user-guides/research

25 Cairo University, Curtin University, Dalian University of Technology (DUT), Indian Institute of Science Bangalore (IISC), Institut Teknologi Bandung (ITB), Loughborough University (LU), Massachusetts Institute of Technology (MIT), Moscow State University (MSU), National Autonomous University of Mexico (UNAM), University College London (UCL), University of Cape Town (UCT), University of Giessen, University of Sao Paulo (USP), University of Tokyo, Wayne State University (WSU).

26 The number of DOIs that are indexed by Unpaywall.

27 The set of unique DOIs for all 15 institutions combined.

28 Dates as per source’s metadata.

29 Originally, there were 150 additional universities, but 10 were removed due to non-coverage or identification issues (e.g., multiple Scopus affiliation IDs). See Appendix 1 for the list of GRID IDs of the additional 140 universities.

30 To see whether these correlations are driven by the size of total output, we have also constructed pairwise scatterplots between the three proportions, with the points colour-coded by total output numbers. The random spread of the colours suggested the correlations are not strongly influenced by size. See Appendix 4.

31 None of the cells in these contingency tables has an expected count less than 10.

32 Using the sampling procedure for that of Fisher’s exact test with 5000 replicates. See https://www.rdocumentation.org/packages/stats/versions/3.6.1/topics/chisq.test

33 p_wos = p_scopus = p_msa generated from a uniform distribution (truncated at ⅓ and 1).

34 Hierarchical clustering is performed using hclust function (base R) with dissimilarity matrix calculated using Gower’s distance in the daisy function (R package cluster). Graphical presentations are produced using R package dendextend.

35 This is the total number of DOIs that are jointed covered by the sources listed in each column title. The numbers here differ slightly with the first Venn diagram in Figure 5 because there exists a small number of DOIs in each source that had repeated entries but fall in different years. The number of such cases for WoS, Scopus and MSA are 1, 2 and 43 respectively.

36 See Harzing & Alakangas (2017b), Hug & Brändle (2017) and https://academic.microsoft.com/faq

37 Calculated out of all DOIs, from the particular source.

38 See for example: https://www.openaire.eu/blogs/open-science-in-indonesia and https://campuspress.yale.edu/tribune/creating-an-open-access-indonesia/

39 Percentage of DOIs from WoS only that are also indexed by at least one of the two other sources but recorded a year apart (in both directions).

40 Percentage of DOIs from WoS only that are also indexed by at least one of the two other sources but recorded two years apart (in both directions).

41 See https://unpaywall.org/data-format

42 Note here the total number of DOIs are slightly lower in each part of the Venn diagram as compared to the left Venn diagram in Figure 5. This is because here we are only including DOIs that are also recorded in Unpaywall.

43 See Figure 3 for the labelling of the Venn diagram.

44 This is the number of DOIs in each source that is also indexed by Unpaywall (i.e., Crossref).

45 These are calculated using the sets of DOIs from each source that are also indexed in Unpaywall. OpenCitations and Unpaywall both use Crossref DOIs as identifiers. If we use the full set of DOIs (i.e., including non-Crossref DOIs) we get a very small increase in citation totals ranging from 0.01% to 0.03%.

46 Only Unpaywall (i.e., Crossref) DOIs are included in the calculations of average citations.

47 https://unpaywall.org/

48 This is the total number of DOIs in each source that are recorded in Unpaywall.

49 Only DOIs indexed by Unpaywall are included in the calculations.

50 This is done via doi.org as first pass, followed by manual title search online.

51 The decision of whether an affiliation is a plausible variant of the target affiliation is made somewhat subjectively but informed via simple online searches. These may include subdivisions under the target affiliation (e.g., departments, research groups, etc), aliases, etc. The strategy is that this should be a simple decision via a quick online search, otherwise a negative response is recorded.

52 There are two Venn diagrams for each institution: left = all DOIs as per WoS, Scopus and MSA; right = DOIs from the three sources that are also recorded in Unpaywall.

53 The points in the scatter plots are colour coded according to the size of output (total number of DOIs), grouped as per plot legend.

54 Parameters estimated from data. Values were generated using the mscFit command in R package fMultivar.

55 parameters estimated from data. Values were generated using the msnFit command in R package fMultivar.

56 One should also note here that having the target affiliation appearing in metadata is not necessarily the same as the object actually being linked to the target affiliation systematically (e.g., API searchable).

57 The other reason would be metadata changes between time of data collection and time of manual cross-validation. But given that we are primary using 2016 data, the scale of this is expected to be relatively small. Manual spot checks did not find any cases where the metadata appears to have changed.

58 Accessed on 22 August 2019.

